# Influenza A Viral Burst Size from Thousands of Infected Single Cells Using Droplet Quantitative PCR (dqPCR)

**DOI:** 10.1101/2024.02.23.581786

**Authors:** Geoffrey K. Zath, Mallory M. Thomas, Emma Kate Loveday, Dimitri A. Bikos, Steven Sanche, Ruian Ke, Christopher B. Brooke, Connie B. Chang

## Abstract

An important aspect of how viruses spread and infect is the viral burst size, or the number of new viruses produced by each infected cell. Surprisingly, this value remains poorly characterized for influenza A virus (IAV), commonly known as the flu. In this study, we screened tens of thousands of cells using a microfluidic method called droplet quantitative PCR (dqPCR). The high-throughput capability of dqPCR enabled the measurement of a large population of infected cells producing progeny virus. By measuring the fully assembled and successfully released viruses from these infected cells, we discover that the viral burst sizes for both the seasonal H3N2 and the 2009 pandemic H1N1 strains vary significantly, with H3N2 ranging from 10^1^ to 10^4^ viruses per cell, and H1N1 ranging from 10^1^ to 10^3^ viruses per cell. Some infected cells produce average numbers of new viruses, while others generate extensive number of viruses. In fact, we find that only 10% of the single-cell infections are responsible for creating a significant portion of all the viruses. This small fraction produced approximately 60% of new viruses for H3N2 and 40% for H1N1. On average, each infected cell of the H3N2 flu strain produced 709 new viruses, whereas for H1N1, each infected cell produced 358 viruses. This novel method reveals insights into the flu virus and can lead to improved strategies for managing and preventing the spread of viruses.

**Author summary:** Viruses infect and exploit host cells to reproduce and spread. The viral burst size, or the number of viral particles released from an infected cell, plays a critical role in understanding infection dynamics and overall viral fitness. However, accurately determining burst size for many single cells using conventional laboratory methods can be challenging. Here, we introduce dqPCR, a droplet microfluidic method for the rapid measurement of influenza virus numbers produced by thousands of individual cells. Our findings revealed that only a small proportion of infected cells are responsible for producing a significant portion of the total viral population. By utilizing this method in future studies, we can gain a deeper understanding of the role of diversity in rapidly evolving viruses.

## Introduction

The number of viral particles produced from an infected host cell is known as the viral burst size. First characterized in bacteriophage populations (1,2), burst size is an important measure of viral fitness and plays an important role in modeling of viral infection dynamics (3). To study viral infections, tissue culture systems or animal models are typically used; however, these methods provide an average measurement of infected cells and may not capture the variation between individual infected cells (4). In the case of influenza A virus (IAV) infection, individual cells exhibit significant heterogeneity in production of infectious virions (5,6), viral gene expression (6–9), and host innate immune activation (8). Previous estimates of IAV burst size indicate a similar trend; average IAV burst size measured from bulk tissue culture was estimated at 11.5 plaque forming units (pfu) per cell (10) while single cell measurements ranged from 1 to 1000 pfu/cell (5). However, these estimates relied on plaque assays and were limited to quantification of fully infectious viral particles. The plaque assays overlooked the majority of non-infectious particles that make up 90-99% of virions in the IAV population that rely on co-infection to propagate (11,12). Therefore, new techniques are needed to accurately determine the IAV burst size by capturing the full range of particle types released from single-cell infections.

Various methods have been used to measure IAV production from single-cell infections, including RT-PCR (5,6,13), RNA sequencing (6–9), flow cytometry (11), and infectivity titer (5,6,13). However, each method has limitations in accurately measuring viral burst size. Infectivity assays only detect fully infectious viral particles and cannot account for incomplete viral particles (11,12) that contribute to viral replication and host immune responses (14). RT-PCR, RNA sequencing, and flow cytometry typically measure intracellular viral replication, neglecting the packaging and release of extracellular viral particles. These methods are limited because intracellular viral transcription may correlate poorly with viral progeny production (9). Additionally, these methods can be low-throughput, capturing only a limited number of single-cell infections (5–7,9,13), and may not be sufficient to study highly diverse viral populations like IAV. Determining IAV burst size distributions, therefore, requires a high-throughput method capable of measuring extracellular viral particles released during single-cell infection. One such method is drop-based microfluidics. This flexible platform enables high-throughput, single cell virology experimentation in isolated microenvironments (15–20). Recently, we demonstrated that IAV can replicate in single cells encapsulated in drops and is comparable to bulk infection (21).

Existing drop-based microfluidic methods for quantifying genome copies rely on endpoint measurements like droplet digital PCR (ddPCR) or real-time fluorescence measurements using qPCR on a microfluidic chip. However, these approaches have limitations. ddPCR can only confirm the presence or absence of nucleic acids (22,23) and cannot quantify the contents of an individual drop. qPCR on a chip can quantify the contents of an individual drop, but requires complex microfluidic devices and results are relatively low-throughput (24–30). Improving the throughput of drop-based qPCR would allow for rapid measurement of a significantly larger number of drops, thus enabling the study of large and diverse viral populations.

In this study, we introduce a microfluidic method called droplet quantitative PCR (dqPCR), which combines our prior work (21,31) with flow-based droplet fluorescence detection. The dqPCR method is used to measure extracellular burst size distributions of thousands of IAV-infected single cells that were encapsulated into drops. Human alveolar epithelial (A549) cells were infected with either influenza A/Perth/16/2009 (H3N2; human seasonal isolate) or influenza A/California/07/2009 (H1N1; 2009 pandemic strain) at multiplicity of infection (MOI) of 0.1. The IAV burst sizes were found to be highly diverse, spanning three orders of magnitude (10^1^ to 10^4^ viruses/cell) for H3N2 and two orders of magnitude (10^1^ to 10^3^ viruses/cell) for H1N1 across thousands of single cell infection measurements. IAV production followed a negative-binomial distribution, with modeled mean burst sizes of 709 and 358 viruses/drop for H3N2 and H1N1, respectively. Notably, we found that 10% of cells produced ≈60% or ≈40% of M gene RNA during H3N2 or H1N1 infections, respectively. The dqPCR method provides a novel high-throughput approach for quantifying released viruses from infected single cells and introduces an unprecedented means for characterizing the infection dynamics of highly heterogeneous viral populations.

## Results

### Drop-based Microfluidic Platform for Measuring Viral Burst Size from Single-Cell Infections

To determine IAV burst size from single cell infections, H3N2 and H1N1 strains were first inoculated onto A549 cells in bulk culture with an MOI of 0.1 pfu/cell (Fig 1A). A low MOI was chosen to ensure that of the few cells that were infected, most cells were infected with a single virion, as variation in cellular MOI could affect our measurements due to co-infection (32). While a low MOI of 0.1 is expected to have few infected cells that produce progeny virus, the high-throughput capability of dqPCR enables the screening of tens of thousands of drops. Single infected cells were then encapsulated into 100 µm diameter microfluidic drops, such that most drops were either empty or contained a single cell (Fig 1B). Drops were incubated for 18 hours post infection (hpi) to allow for replication of IAV progeny virus (Fig 1C) with numbers comparable to bulk infection (*SI Appendix,* Fig S1). Interestingly, the supernatant collected from drops produced slightly more progeny virus than bulk (2.4× to 6.8× greater production, p << 0.05). The burst sizes of single-cell infections were quantified by the number of IAV genome copies (cpd), which can be equated to viruses per cell for each drop that contains an infected cell producing progeny virus. The genome copies were measured using a multiplexed RT-qPCR assay targeting two template sequences, the IAV M gene and cellular β-actin (*SI Appendix,* Table S1), following our previous work (31). A microfluidic chip (20) was used to split off a 1/8^th^ portion of the drop containing progeny virus and merge with a continuous flow of RT-qPCR mix (Fig 1D). This sampling method isolates extracellular viral progeny released from the infected host cell, allowing us to quantify genome copies only from fully assembled and released extracellular virions. Note that our recent work indicates minimal amplification efficiency loss when directly amplifying infected supernatant in drops compared to bulk qPCR (21). Based on this finding, we assume that using an M gene stock serves as a suitable standard for comparing our infected drop samples. Drops containing progeny virus and RT-qPCR mix were thermocycled using a standard qPCR machine and removed at successive PCR cycle numbers, *N* (Fig 1E). Normalized drop fluorescence intensities (11*R_N_, SI Appendix*, M5) were measured by flowing drops through a microfluidic device mounted on a custom optical detection microscope (33) (Fig 1F). This flow-based detection method significantly increased the throughput of our drop fluorescence measurements during burst size experiments compared to our previous static image analyses (31). Δ*R_N_* of filtered drops were converted to M gene and β-actin cpd using dqPCR (Fig 1G). A variable filter was applied to drops containing high amounts of β-actin to exclude them from the final burst size distributions (*SI Appendix*, R8). These drops were assumed to be contaminated with intracellular lysate and were not included in further analysis. The abundance of M gene RNA across thousands of single-cell infections revealed a wide distribution of IAV burst sizes (Fig 1H).

**Fig 1.**
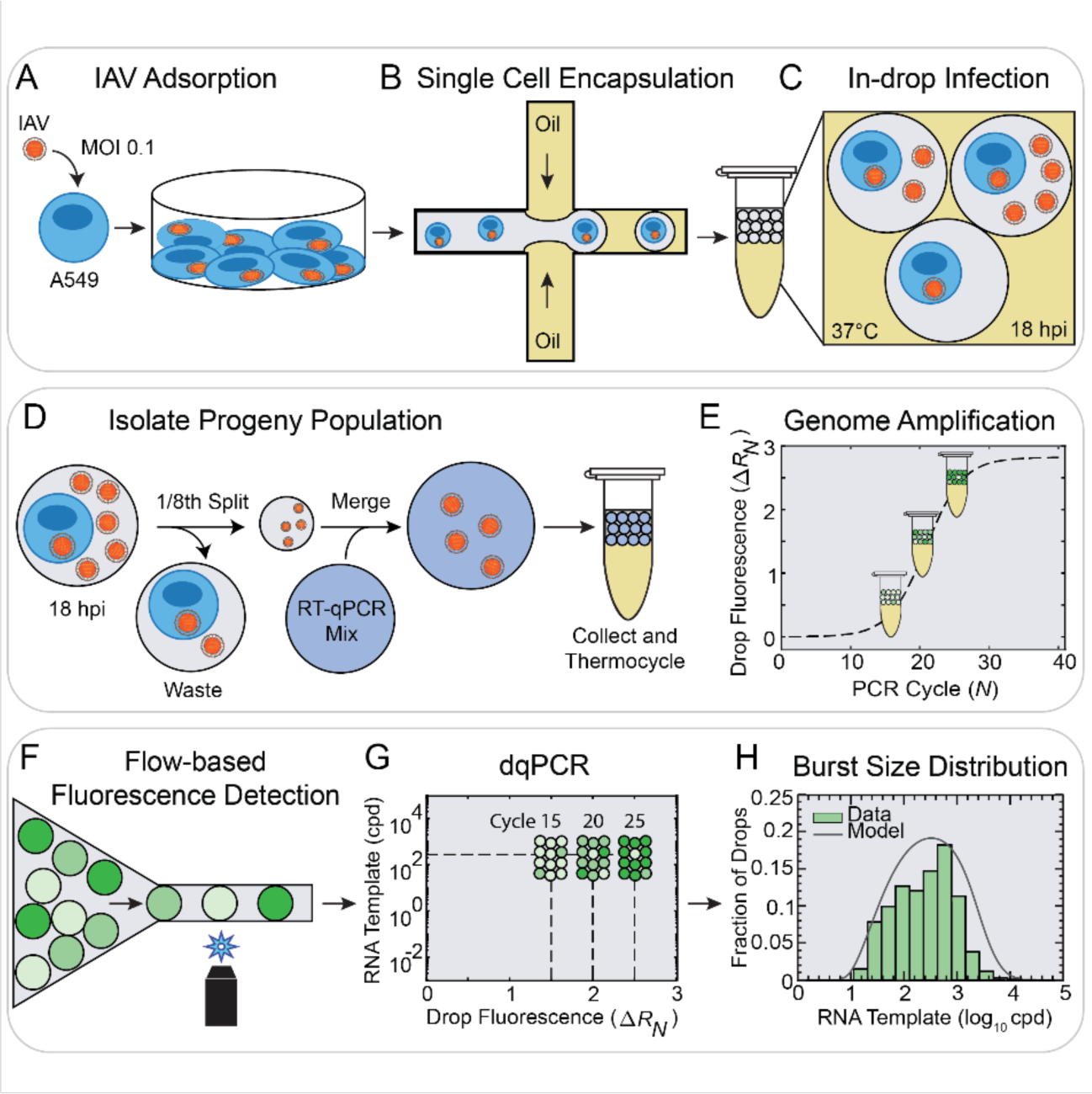
Drop-based Microfluidic Platform for Measuring Viral Burst Size from Single-Cell Infections. **(A)** IAV (H1N1 or H3N2) was inoculated onto plated A549 cells at an MOI of 0.1 pfu/cell. **(B)** Single infected cells were encapsulated into 100 µm drops using a flow-focusing device. **(C)** Drops were incubated at 37 °C for 18 hpi to allow for replication of IAV progeny virus. **(D)** In-drop infections were injected into a microfluidic device that splits 1/8^th^ of the extracellular progeny virus from the host cell and merges these viruses with a continuous flow of multiplexed RT-qPCR mix targeting IAV M gene RNA and cellular β-actin mRNA. **(E)** Drops were thermocycled in a standard qPCR machine and sampled at discontinuous *N*. **(F)** 11*R_N_* was measured serially using flow-based detection. **(G)** 11*R_N_* values were converted to RNA cpd using dqPCR. **(H)** IAV burst size distributions of M gene RNA cpd from thousands of single-cell infections (green bars). The mean burst size of each distribution was calculated from a parametric model (gray curve) fitted to our data.

### Droplet Quantitative PCR (dqPCR) Method for Converting Drop Fluorescence to Nucleic Acid Concentration

The dqPCR method allows for the construction of standard curves that directly relate 11*R_N_* (normalized drop fluorescence) to M gene RNA copy number (cpd). In traditional RT-qPCR, standard curves plot the cycle threshold (*Ct*) at which reporter fluorescence rises above background for a dilution series of known RNA template concentrations (*SI Appendix*, M7). This method requires continuous measurements of fluorescence amplification across all *N*. However, in dqPCR, drops are sampled discontinuously and only require evaluation of sample fluorescence at a single *N*.

To construct standard curves at a single *N* using dqPCR, we generated a reference amplification curve for a known RNA template concentration amplified in drops (Fig 2A, solid blue curve, 1.71 × 10^2^ M gene cpd). Drop 11*R_N_* was measured at *N* = 1, 10, 13, 16, 19, 22, 28 and 40 (Fig 2A, black dots). A continuous amplification curve was fit to the discontinuous drop fluorescence measurements using sigmoidal curve fitting using PCR efficiency (SCF-E) (see *SI Appendix*, M8). The model determined four PCR efficiency (*E_N_*) parameters that best fit the 11*R_N_* values of the sampled *N* (*E_max_*, *E_min_*, *N_0.5_*, and *k*). Once fit, these parameters were used to calculate the *E_N_* curve for all *N* (Fig 2A, dashed black curve). The *E_N_* curve was then used to produce a 11*R_N_* reference amplification curve that included non-sampled *N* (Fig 2A, solid blue curve). SCF-E was validated for bulk RT-qPCR of M gene RNA (*SI Appendix,* Fig S5A) across five orders of magnitude (*SI Appendix*, R2 and Table S2). Additionally, SCF-E was validated for drops of 1.71 × 10^2^ M gene cpd (*SI Appendix*, Fig S12A).

**Fig 2.**
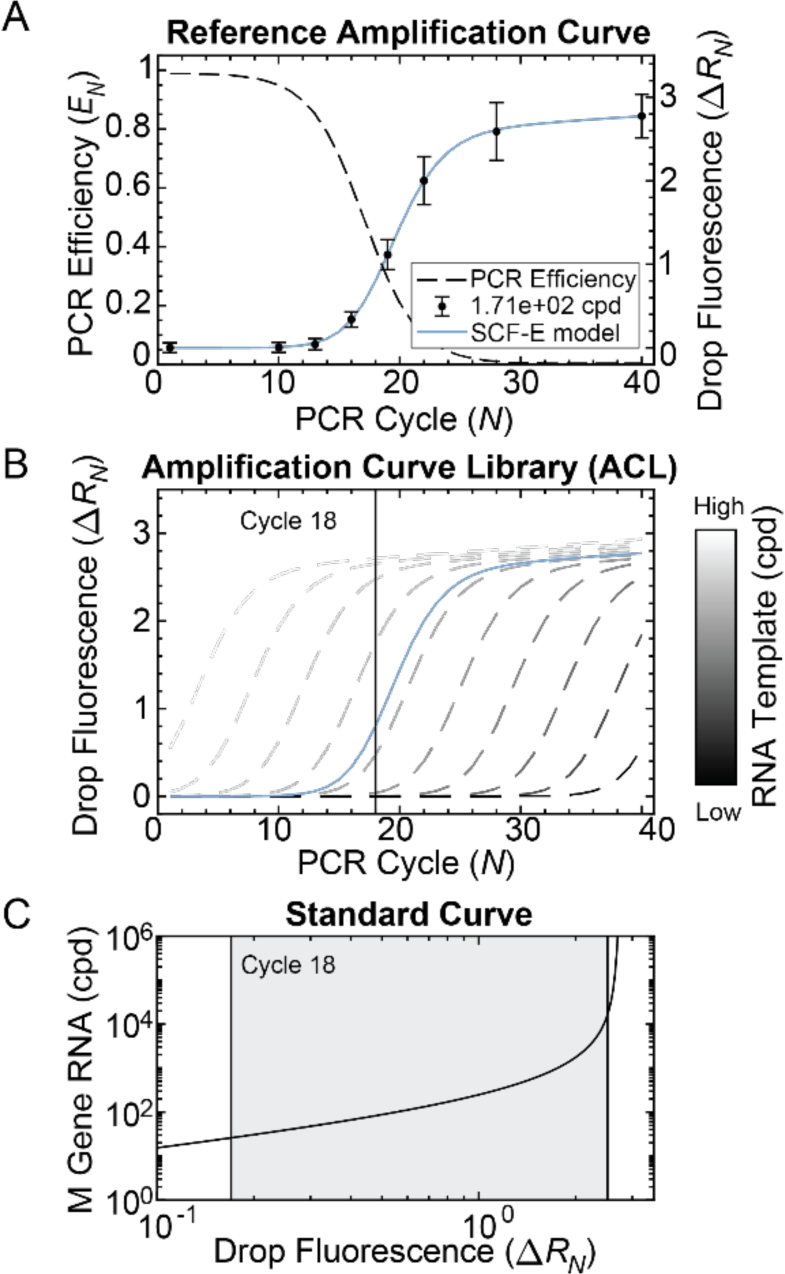
Droplet Quantitative PCR (dqPCR) Model for Converting Drop Fluorescence to Nucleic Acid Concentration. **(A)** M gene RNA (1.71 x 10^2^ cpd) was amplified in 50 µm drops. 11*R_N_* was detected at *N* = 1, 10, 13, 16, 19, 22, 28, 40 and displayed as the mean (black dots) with error bars representing one standard deviation. Approximately 1000 drops were detected at each cycle. A continuous reference amplification curve (blue solid curve) was reconstructed from discontinuous 11*R_N_* measurements using SCF-E. The *E_N_* curve (black dashed curve) was fit to 11*R_N_* at sampled cycle numbers to calculate 11*R_N_* at non-sampled cycle numbers. **(B)** The efficiency parameters fit from the reference amplification curve (blue solid curve) were used to model 1000 virtual amplification curves (gray dashed curves, subset of 10 shown), creating a high-resolution dilution series called an Amplification Curve Library (ACL), representing 1000 unique M gene RNA concentrations. **(C)** Standard curves were constructed using 11*R_N_* from all virtual curves at a single *N* (*i.e.*, cycle 18). The valid range of the standard curve (gray shaded area) is determined by the min and max 11*R_N_* of the reference curve.

The *E_N_* parameters obtained from the reference amplification curve were used to model virtual amplification curves, creating a high-resolution dilution series referred to as an Amplification Curve Library (ACL) (*SI Appendix*, M9). The ACL simulated 1000 virtual amplification curves, each with a unique RNA template concentration (Fig 2B, dashed gray curves, subset of 10 shown). Once constructed, the ACL was used to generate a standard curve that related RNA template concentration to 11*R_N_* at a single *N* (Fig 2C). This was done by recording 11*R_N_* of every virtual curve at a single *N* (*e.g.*, *N* = 18) where unknown drops were sampled (Fig 2B, black vertical line). The valid range for conversion with the standard curve (Fig 2C, gray shaded area) was determined by the 11*R_N_* values of the reference curve (Fig 2A). The maximum valid 11*R_N_* was calculated as the mean minus the standard deviation of 11*R_N_* at *N* = 40, which removes drops where amplification has plateaued. The minimum valid 11*R_N_* was set at the 99th percentile of 11*R_N_* at *N* = 1 to remove background fluorescence. ACL standard curves were shown to be robust across *N* for both bulk (*SI Appendix*, R4) and drops (*SI Appendix,* Fig S12B). Additionally, we tested ACL standard curves for human and plant DNA template sequences (*SI Appendix*, R5) and found that 11*R_N_* accurately converted to genome concentrations within a two-fold change of the expected means. Together, the SCF-E model for generating reference amplification curves and the ACL method of constructing standard curves form the dqPCR analysis pipeline that enables the conversion of drop fluorescence to cpd (Fig 2).

### Resolving IAV M Gene RNA Concentrations from Single and Mixed Droplet Populations

To validate the capability of dqPCR to resolve a range of RNA concentrations in drops, we conducted a control experiment amplifying three known concentrations of M gene RNA (1.71 × 10^1^, 1.71 × 10^2^, and 1.71 × 10^3^ cpd) in 50 μm drops (Fig 3A). These RNA concentrations were selected because they fall with the expected range of IAV single-cell burst sizes from previous estimates (5,9,10). 11*R_N_* was measured for each group individually at multiple intermediate *N* and converted to M gene cpd using dqPCR (*SI* Appendix, Table S6). A new reference amplification curve of 1.71 × 10^2^ M gene cpd was generated for each M gene group, as PCR kinetics can vary from day to day (*SI Appendix,* Fig S13). The measured mean M gene cpd at each sampled *N* for all concentrations fell within a 2-fold change of the expected concentration (*SI Appendix,* Fig S14). A linear mixed-effects model showed minimal deviation from the expected concentration at different cycle numbers (<1% difference/cycle, *SI Appendix*, R6).

**Fig 3.**
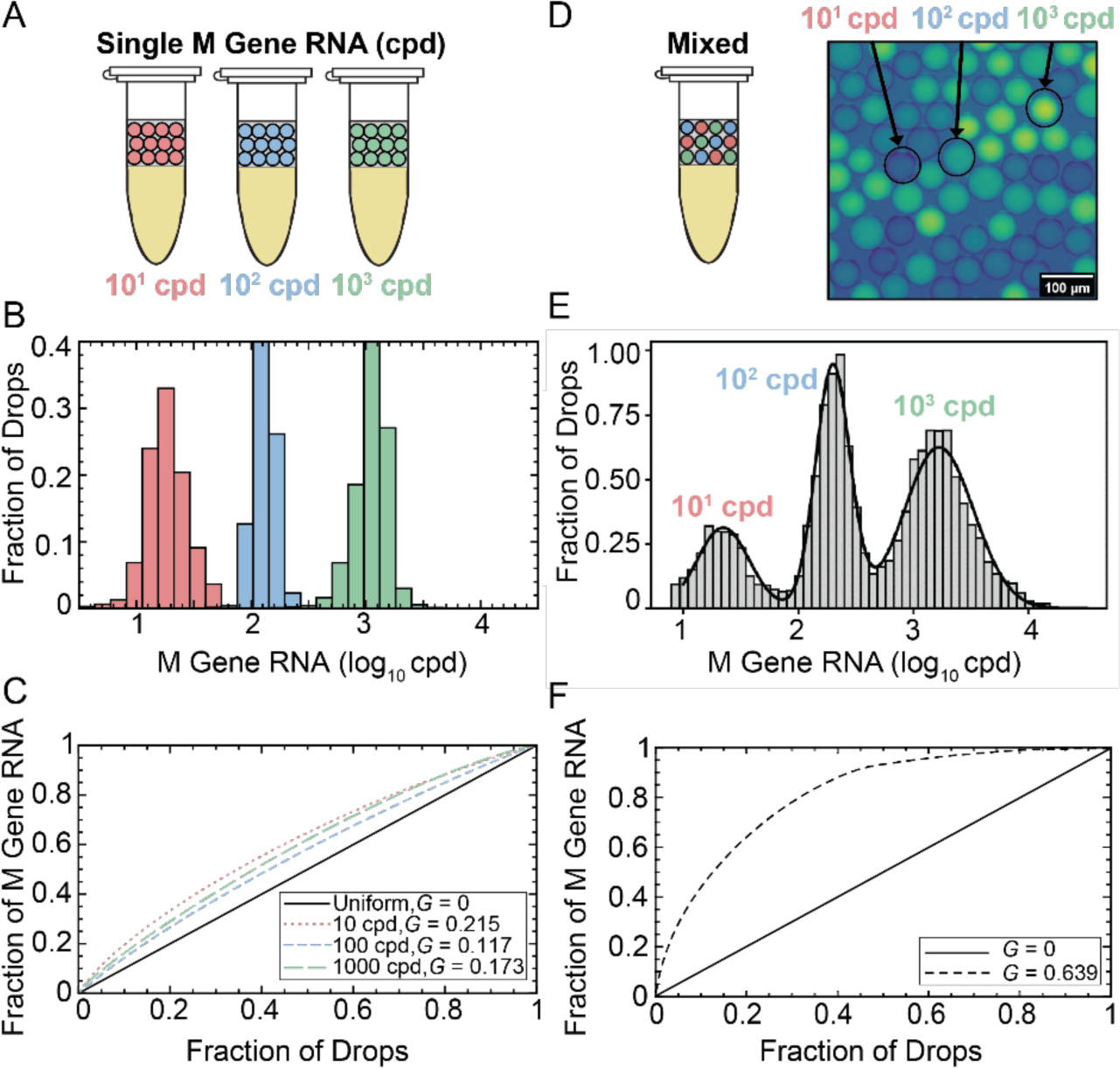
Resolving IAV M Gene RNA Concentrations from Single and Mixed Droplet Populations. **(A)** Three known concentrations of M gene RNA, 1.71 × 10^1^ cpd (red), 1.71 × 10^2^ cpd (blue), and 1.71 × 10^3^ cpd (green) were amplified in 50 µm drops. **(B)** 11*R_N_* was detected at multiple different *N* (Table S6) within each group and converted to M gene cpd. M gene cpd measurements were pooled to a single distribution for each group, with means of 2.03 × 10^1^ cpd (red, *n* = 4,513 drops); 1.31 × 10^2^ (blue, *n* = 15,899 drops); 1.16 × 10^3^ (green, *n* = 65,249 drops). **(C)** The uniformity of each distribution was quantified using *G*. **(D)** Three M gene RNA concentrations in drops were collected in a mixed sample. **(E)** Mixed 11*R_N_* were detected at *N* = 16-19, converted to M gene cpd, and pooled to a single distribution (grey bars, *n* = 68,748). Three individual peaks were isolated from the mixed distribution using GMM analysis (black curve) (Table S7). **(F)** The heterogeneity of the mixed population compared to the individual populations in (C) was significantly greater with *G* = 0.639.

Thus, M gene cpd measurements from all *N* were pooled together into single distributions with means of 2.03 × 10^1^ cpd (red, *n* = 4,513 drops), 1.31 × 10^2^ (blue, *n* = 15,899 drops), 1.16 × 10^3^ (green, *n* = 65,249 drops) (Fig 3B). Distributions were centered around the expected mean and exhibited a linear relationship between measured and expected cpd, falling within a 2-fold change of the expected mean (*SI Appendix*, Fig S17). The heterogeneity of each distribution was examined using the Gini coefficient (*G*) (Fig 3C); a value typically used to measure the degree of wealth inequality and ranges from 0 to 1 (34). A *G* value of 0 indicates total equality, where each cell produces the same number of viruses, while a *G* value of 1 signifies absolute inequality, where one cell produces all the viruses. Thus, a higher *G* corresponds to a more heterogeneous distribution. The uniformity of each M gene distribution (Fig 3C) was as follows: 10^1^ cpd (*G* = 0.215), 10^2^ cpd (*G* = 0.117) or 10^3^ cpd (*G* = 0.173). These values represent the baseline level of noise for single concentrations M gene RNA in drops (∼*G* < 0.22) (Fig 3C).

To further confirm the ability of dqPCR to resolve a mixed drop population, we conducted another control experiment in which the same three known concentrations of M gene RNA (1.71 × 10^1^, 1.71 × 10^2^, or 1.71 × 10^3^ cpd) were separately encapsulated into 50 µm drops and collected as a mixed sample prior to thermocycling (Fig 3D). 11*R_N_* of the mixed drops was detected at *N* = 16-19 and converted to M gene cpd, enabling high-throughput analysis with over 69,000 drops sampled (Fig 3E). The data from the mixed sample yielded three individual distributions and were isolated using a Gaussian mixture model (GMM) (*SI Appendix*, R7 and Table S7). *G* for the mixed drop sample containing 10^1^ to 10^3^ M gene cpd was 0.639 (Fig 3F), demonstrating greater heterogeneity compared to the measurement of single RNA concentrations in drops, as expected. These results demonstrate the ability to resolve three M gene RNA concentrations from a mixed concentration of drops.

### H3N2 and H1N1 Single-Cell Infection Burst Size Distributions

To determine the burst size distributions of H3N2 and H1N1 populations, we measured M gene RNA abundance in >10^4^ drops containing infected cells producing progeny virus. Infected drops were processed by isolating 1/8^th^ of a drop containing progeny viruses and merging it with RT-qPCR mix. The merged drops were thermocycled and analyzed using flow-based fluorescence detection. To ensure that the experimental burst size measurements only included extracellular viral progeny and did not include intracellular viral RNA from cell lysate, a multiplexed RT-qPCR assay was used to detect the presence of an abundant cellular protein, β-actin (*SI Appendix,* Fig S18). Drops with high levels of β-actin were filtered out from the final burst size distributions using a variable filter (*SI Appendix*, R8). Filtered drops were quantified at *N* = 16, 19, 22, 25, and 28 (*SI Appendix,* Table S8) using dqPCR and pooled to create single burst size distributions. For each biological replicate, a new SCF-E reference curve (*SI Appendix,* Fig S20) was used for dqPCR (*SI Appendix,* Table S9). As an additional control to ensure that the measurements included only extracellular viral progeny, single IAV-infected cells in drops were treated with oseltamivir, a drug preventing the cleavage of assembled viral particles from the host membrane. Infected cells treated with oseltamivir exhibited M gene RNA abundance comparable to mock infected cells at 18 hpi. (SI Appendix, R9, Fig S21).

Experimental IAV burst sizes revealed increased progeny virus production in the H3N2 population compared to H1N1, as observed from a random sample of 100 drops and plotting in ascending order of M gene RNA (Fig 4A). The burst size distributions were highly diverse, ranging from ∼10^1^ to ∼10^3^ M gene cpd for H1N1 (*n* = 5,585 drops) and an order of magnitude higher, from ∼10^1^ to ∼10^4^ M gene cpd, for H3N2 (*n* = 9,620 drops) (Fig 4B). The mean of the H3N2 distribution was 1.0 × 10^3^ M gene cpd (Fig 4B, blue bars). This fell outside of the interquartile range (IQR) of the distribution, with lower (25^th^ percentile) and upper (75^th^ percentile) bounds of 93 and 890 M gene cpd, respectively (*SI Appendix,* Table S8). The mean can fall outside of the IQR in a highly skewed distribution, which occurs when a small fraction of the infected cell population produces a greater proportion of virus. The mean of the H1N1 burst size distribution was 4.8 × 10^2^ M gene cpd (Fig 4B, red bars), falling within the IQR of 88 and 630 M gene cpd (*SI Appendix,* Table S8). This suggests less heterogeneity in the H1N1 population, which is further supported by examining the uniformity of each distribution using *G* (Fig 4C). The H3N2 distribution is less uniform (*G_H3N2_* = 0.705) compared to H1N1 (*G_H1N1_* = 0.586). In both viruses, a minor population of infected cells produces a large fraction of progeny virus. For H3N2, 10% of infected cells produced ≈60% of measured progeny virus (Fig 4C, blue dotted line). For H1N1, 10% of infected cells produced ≈40% of progeny virus (Fig 4C, red dashed line).

**Fig 4.**
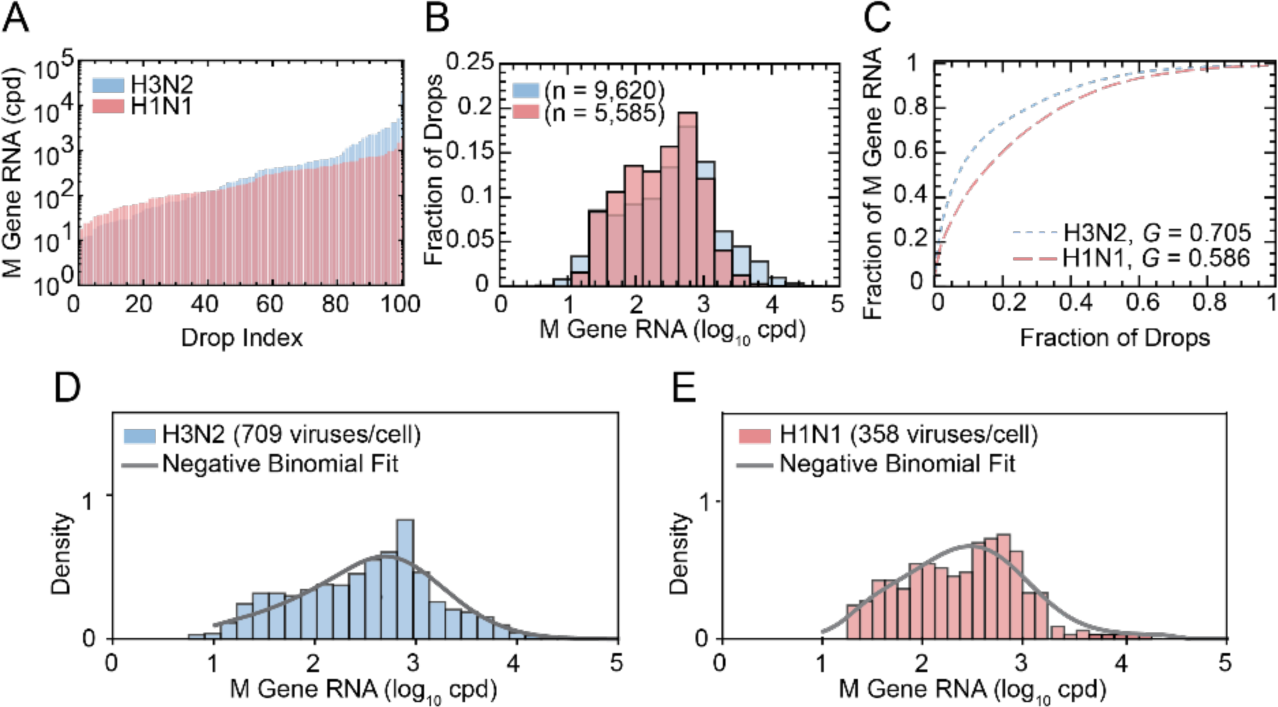
H3N2 and H1N1 Single-Cell Infection Burst Size Distributions. IAV burst sizes of H3N2 (blue) and H1N1 (red) were sampled at *N* = 16, 19, 22, 25, and 28 across three biological replicates (Table S8). **(A)** Low to high M gene RNA (cpd) from a random sample of drops (*n* = 100) for H3N2 (blue) and H1N1 (red). **(B)** Single burst size distributions for H3N2 (blue, *n* = 9,620) and H1N1 (red, *n* = 5,585) populations. The measured mean burst sizes were 1.0 × 10^3^ M gene cpd (H3N2) and 4.8 × 10^2^ M gene cpd (H1N1). **(C)** The H3N2 distribution (*G* = 0.705) was less uniform than H1N1 (*G* = 0.586). **(D-E)** Distributions were fitted to a negative binomial (grey curve, Table S10). The modeled mean burst sizes were (D) 709 cpd (H3N2) and (E) 385 cpd (H1N1).

To estimate the most accurate mean burst size, we apply a statistical model (*SI Appendix*, R10) that considers both measurement noise and distribution shape. The model demonstrated that the burst size data is best fit by a negative-binomial distribution (Fig 4D-E). The shape parameter of the negative binomial distribution was estimated to be 0.48 and 0.49 for H3N2 and H1N1, respectively (Table S10). This indicates extreme cell-to-cell variability in burst sizes for both H3N2 and H1N1 populations, as the negative-binomial distribution is considered highly dispersed when the shape parameter is less than 1. Using the negative binomial fit, we estimated that the modeled mean burst sizes for each population were 709 viruses/cell for H3N2 and 358 viruses/cell for H1N1 (Fig 4D-E).

## Discussion

We introduce a droplet microfluidic method called dqPCR to quantify viral burst size from thousands of single infected cells in microfluidic droplets. dqPCR is a quantitative PCR technique for droplets that does not require specialized microfluidic devices or continuous measurements of a sample. This has led to a drastic improvement in throughput compared to previous on-chip qPCR work (24–30). Compared to ddPCR, which measures only single genome copies per drop, dqPCR can resolve a range of genome copies from a single copy to up to thousands of copies per drop.

Estimating viral burst sizes using population averaging masks cell-to-cell variability. Additionally, single-cell burst size estimates based upon infectivity, like the plaque assay, do not account for the production of incomplete particles that are abundant in IAV populations (11,12). Here, we account for both infectious and non-infectious particles by quantifying viral genome abundance. Moreover, we account for single cells in a population without averaging, to investigate whether one cell produces more progeny or if all cells produce equal progeny. We measured IAV burst sizes across >10^4^ drops containing cells producing H3N2 or H1N1 progeny virus. This has yielded the largest estimate of single-cell viral burst size to date. The range of our burst size measurements spanned from 10^1^ to 10^4^ viruses/drop, which is an order of magnitude wider than previous, lower throughput (*n* = 430) estimates of H1N1 infectious virus production in single cells (10^1^ to 10^3^ pfu/cell) (5). This is likely due to our method that measures all viral genome copies rather than infectious particles. A mean burst size of 709 and 358 viruses/cell was modeled for H3N2 and H1N1 strains, respectively, from highly dispersed negative-binomial distributions. The mean burst size for H3N2 (709 viruses/cell) compares similarly with a previous mean estimate of 962 viruses/cell for H3N2 in a bulk culture (10). One caveat of this study is that some drops contained more than one cell per drop. Across both populations, for a drop that contained a cell or cells, an average of 86% of these drops were loaded with 1 cell, 12% with 2 cells, and 2% with 3 cells or more (*SI Appendix,* Fig S23). Another limitation of this study is that the cells were not synchronized to the same growth stage, which may influence viral numbers. Future work can improve upon our methods to encapsulate exactly one cell per drop (35,36) and explore cell synchronization.

The observed heterogeneity in the single cell burst sizes (*G*_H3N2_ = 0.705 and *G*_H1N1_ = 0.586) indicates that a minor population (10%) of infected host cells produced a large fraction of progeny virus (≈60% for H3N2 and ≈40% for H1N1) (Fig 4C). While our burst size measurements quantify the number of M gene segments, similar heterogeneity has been observed when quantifying the number of HA and NA segments using single-cell sequencing (*G* = 0.78), where 6.5% of infected cells generated over 50% of the counted progeny virus (9), although this study examined fewer single cells (*n* = 92). Additionally, these trends align with single-cell transcriptomic studies of IAV infection, which show that 10-15% of infected cells are responsible for 50% of viral gene expression (7), although levels of intracellular transcripts are not well correlated to progeny virus production (9).

This study is the first to compare viral burst sizes of two different clinically relevant human IAV strains and provides the first burst size estimate for the H3N2 strain, revealing differences in progeny production between the two strains. Future research could investigate specific viral and host factors that contribute to the observed heterogeneity in viral output, both within and between IAV strains, as well as the extent to which *in vitro* measurements reflect burst sizes across different host cell types *in vivo*. As dqPCR is a multiplexed assay, it can be adapted to simultaneously measure multiple template sequences, such as other genome segments within IAV or both virus and host templates. In the case of IAV, dqPCR could be applied to investigate segment reassortment (37) or the frequency of incomplete viral genomes (10–12), or it could be extended to study genome recombination in other viruses (18). The dqPCR method presented here not only expands our current toolset for high-throughput single-cell characterization, but also enables the detection of nucleic acid abundance on an unprecedented scale. Future research can leverage the highly resolved single-cell data sets provided by dqPCR to further characterize host-virus interactions, improve diagnostics, or develop more effective vaccines and anti-viral treatments.

## Materials and methods

### Virus Strains and Cell Lines

Influenza A virus (IAV) strains H1N1 (A/California/07/09) and H3N2 (A/Perth/16/2009) were obtained as seed stock from Dr. Christopher Brooke (University of Illinois Urbana-Champaign) and expanded in Madin-Darby canine kidney cells (MDCK) and human alveolar epithelial cells (A549) obtained from ATCC (CCL-34, CCL-185). MDCK cells were maintained in Dulbecco’s Modified Eagle medium (Corning) supplemented with 10% fetal bovine serum (FBS) (HyClone) and 1× penicillin-streptomycin (pen/strep) (Fisher Scientific). A549 cells were maintained in Hams F-12 medium (Corning), also supplemented with 10% FBS and 1× pen/strep. IAV seed stocks were generated using the JetPRIME kit (Polyplus) to co-transfect Human kidney epithelial-like cells (293T) with 8 plasmids each carrying the sequence of one IAV genome segment: pDZ::PB2, pDZ::PB1, pDZ::PA, pDZ::HA, pDZ::NP, pDZ::NA, pDZ::M, and pDZ::NS. Plaque isolates from transfection supernatants were then amplified in MDCK cells to produce IAV seed stocks. The seed stock virus population (P1) was expanded into working stocks (P2), which were used for all experiments, and generated by infecting MDCK cells with seed stock at an MOI of 0.01 pfu/cell. Working stock titers were determined by plaque assay on MDCK cells. IAV burst sizes were measured from single-cell infection of A549 cells at an MOI of 0.1 pfu/cell.

### Microfluidic Device Fabrication

Microfluidic devices were fabricated using polydimethylsiloxane (PDMS) (Sylgard 184) by patterning SU-8 photoresist on silicon wafers using standard soft photolithography.

### IAV Infection

A549 cells were seeded onto 6-well plates at a concentration of 1 × 10^6^ cells/well and incubated at 37 °C with 5% CO_2_ for 24 hrs. Enough wells were plated to collect a final cell suspension of 2 × 10^6^ cells/mL following infection. For our experiments, we plated cells onto 6 wells for infection samples and 4 wells for mock infection controls. Confluent cell monolayers were washed twice with 1× PBS (Corning) and infected at an MOI of 0.1 with H1N1 or H3N2 working stocks suspended in A549 infection media (Ham’s F-12 medium supplemented with 1 mM HEPES (HyClone), 1× pen/strep, 0.2% Bovine Serum Albumin (BSA) (MP Biomedical) and 1 µg/mL TPCK-trypsin (Worthington Biomedical)). Infected cells were incubated at 37 °C for 1 hr. Mock infected cells were treated with media without the virus and otherwise processed the same as infected cells. After 1 hr, supernatants were removed and replaced with 2 mL of fresh media for an additional 1 hr incubation period at 37 °C. Following the second incubation period, cells were washed twice with 1× PBS. Cells were then detached from wells using 300 µL of 0.25% Trypsin containing 1× EDTA (Fisher Scientific) for 5 min. Next, 700 µl of A549 culture media (Hams F-12 medium supplemented with 10% FBS and 1× pen/strep) was added to the cells to neutralize trypsinization. Cells were collected into two final suspensions, infected and mock, and the wells were washed one more time with 500 µl of A549 culture media. Cell suspensions were centrifuged at 500 × *g* for 5 min to pellet, washed with 1× PBS, and centrifuged a second time. The washed cell pellets were each resuspended in infection media (Ham’s F-12 media supplemented with 1 mM HEPES, 1× pen/strep and 0.2% BSA) at a final concentration of 2 × 10^6^ cells/mL. For each IAV strain (H1N1 or H3N2), we sampled drop and bulk infections, with mock infection controls at the following timepoints: drop infection (0 and 24 hpi), bulk infection (0 and 24 hpi), mock drop infection (24 hpi), mock bulk infection (24 hpi). This required final cell suspension volumes of 3 mL for infected cells (2 mL for drop infections, 1 mL for bulk infections) and 2 mL for mock-infected cells (1.5 mL for mock drop infections, and 500 µl for mock bulk infection). For bulk infection and mock samples, 500 µl of cell suspensions were re-plated onto 3 wells for a final concentration of 1 × 10^6^ cells/well. Re-plated cells were either incubated at 37 °C with 5% CO_2_ for 24 hpi or immediately frozen at −80 °C for 0 hpi. Drop infection protocols are described in the following section.

### Droplet Encapsulation of Infected Cells

Single A549 cells infected with either H1N1 or H3N2 were encapsulated in 100 µm diameter drops using a flow-focusing drop-making device. Mock-infected cells were encapsulated using the same procedure. A 2 mL suspension of 2 × 10^6^ cells/mL in media was loaded into a 3 mL syringe and injected into the device at 1000 µL/hr using syringe pumps (New Era NE-1000). Simultaneously, 4.5 mL of 1.5 wt% (w/w) solution of PEG-PFPE_2_-based surfactant (RAN Biotechnologies, #008-FluoroSurfactant) in fluorinated HFE 7500 oil (3M) was loaded into a 5 mL syringe and injected at 3000 μL/hr. Drops were collected for 30 min into a 5 mL syringe. Syringes were capped and incubated at 37 °C and 5% CO_2_ for 24 hpi to allow for viral replication in drops. Drop infections sampled at 0 hpi were immediately frozen overnight at −80 °C.

### Droplet Quantitative PCR (dqPCR) of Isolated Progeny Viruses

A multiplexed RT-qPCR assay was used to simultaneously amplify both IAV M gene RNA and cellular β-actin mRNA (31). Drops containing infected cells were split and merged with RT-qPCR mix. Drops were thermocycled and drop fluorescence was measured using epifluorescence or flow-based fluorescence detection.

More detailed methods are in the *SI Appendix, Supplementary Materials and Methods*.

## Data availability

Data supporting this study are included within the article and/or supporting materials. The datasets generated during and/or analyzed during the current study are available on the Chang Lab GitHub repository at https://github.com/chang-lab-mayo-clinic/Influenza-A-Burst-Size.

## Acknowledgements

This work was supported by Defense Advanced Research Projects Agency (DARPA) grant W911NF-17-2-0034 to C.B.C., R.K., C.B.B. (https://www.darpa.mil/), National Institutes of Health (NIH) National Institute of Allergy and Infectious Diseases (NIAID) grant 1R56AI156137-01 to C.B.C. (https://www.niaid.nih.gov/), and by National Science Foundation (NSF) CAREER grant 1753352 to C.B.C. (https://www.nsf.gov/). The funders had no role in study design, data collection and analysis, decision to publish, or preparation of the manuscript. We thank Humberto Sanchez for help with protocols and Carter Hoffman for helpful discussions.

## Author contributions

**Conceptualization:** Geoffrey K. Zath, Ruian Ke, Christopher B. Brooke, and Connie B. Chang

**Data Curation:** Geoffrey K. Zath and Mallory M. Thomas

**Formal Analysis:** Geoffrey K. Zath, Mallory M. Thomas, Steven Sanche, and Ruian Ke

**Funding Acquisition:** Ruian Ke, Christopher B. Brooke, and Connie B. Chang

**Investigation:** Geoffrey K. Zath, Mallory M. Thomas, Emma Kate Loveday, and Dimitri A. Bikos

**Methodology:** Geoffrey K. Zath, Mallory M. Thomas, Emma Kate Loveday, Dimitri A. Bikos, and Connie B. Chang

**Project Administration:** Geoffrey K. Zath, Connie B. Chang

**Resources:** Ruian Ke, Christopher B. Brooke, and Connie B. Chang

**Software:** Geoffrey K. Zath

**Supervision:** Connie B. Chang

**Validation:** Geoffrey K. Zath and Mallory M. Thomas

**Visualization:** Geoffrey K. Zath and Mallory M. Thomas

**Writing – Original Draft:** Geoffrey K. Zath

**Writing – Review & Editing:** Geoffrey K. Zath, Mallory M. Thomas, and Connie B. Chang with additional input from all authors.

## Supporting information captions

### Supplementary materials and methods

(M1) RT-qPCR Template and Primer Sequences. (M2) Microfluidic Device Fabrication.

(M3) Bulk RT-qPCR.

(M4) Droplet Quantitative PCR (dqPCR) of Isolated Progeny Viruses.

(M5) Drop Fluorescence Imaging.

(M6) Drop Flow-based Fluorescence Detection.

(M7) Cycle Threshold (*Ct*) Method of Creating Standard Curves for RT-qPCR.

(M8) Sigmoidal Curve Fitting using PCR Efficiency (SCF-E).

(M9) Amplification Curve Library (ACL) Method of Constructing Standard Curves for RT-qPCR.

## Supplementary results

(R1) Comparing M Gene Abundance During Bulk and Drop Infections.

(R2) Validating SCF-E with Bulk RT-qPCR of IAV M Gene.

(R3) Setting a PCR Efficiency Constant (*E_N_*) for the ACL.

(R4) Validating ACL Standard Curves for Multiple PCR Cycle Numbers.

(R5) Validating ACL Standard Curves for Multiple Template Sequences.

(R6) Validating dqPCR for Multiple PCR Cycle Numbers.

(R7) Extracting Individual Distributions from Mixed Droplet Detection Data.

(R8) Filtering of Drops Containing Cell Lysate from Burst Size Distributions.

(R9) Oseltamivir (OST) treatment of IAV Infected Cells During Drop Infections.

(R10) Modeling IAV Burst Size Distributions from Droplet RNA Concentration.

## Supplementary figures

**Figure S1. Comparison of IAV M Gene Abundance During Bulk and Drop Infections.** IAV strains (H1N1 and H3N2) were separately used to infect A549 cells under bulk cell culture and single cell encapsulation in 100 µm diameter microfluidic drops. M gene abundance (copies/µL) was measured using a bulk RT-qPCR assay from the supernatant of both bulk and drop infections at 0 and 18 hpi. Experimental details can be found in **SI R1**. Each bar represents the pooled data from three replicate experiments. Error bars represent one standard deviation.

**Figure S2. Sigmoidal Curve Fitting using PCR Efficiency (SCF-E).** A four-parametric sigmoid function was used to model decreasing PCR efficiency (*E_N_*) at each cycle number (*N*) (Eq. S3). This model was fitted to discontinuous fluorescence measurements (*F_N_*) of a known RNA template concentration (*C_RNA, ref_*) to generate a continuous reference amplification curve (Eq. S4 and S5). Reference curves used in burst size experiments were generated with *C_RNA, ref_* = 1.71 × 10^2^ RNA (cpd). In the SCF-E model, *E_N_* is the PCR efficiency at cycle *N; E_max_* is the maximum PCR efficiency at cycle number 1 and ranges from of 0.9 to 1; *E*_min_ is the minimum PCR efficiency at cycle number 40 and ranges from 0 to 0.1; *N*_0.5_ is the cycle number at which *E_N_* equals 0.5; and *k* is the shape parameter of the curve. As *k* increases, the shape of the curve flattens, and as *k* decreases, the shape of the curve becomes steeper. Thirty values of each of the four efficiency parameters (*E*_max_, *E*_min_*, N_0.5_, and k*) were used to test 8.1×10^5^ (30^4^) curve fits to experimental fluorescence measurements. The set of efficiency parameters yielding the best fit were used to construct the SCF-E reference curve (solid blue curve).

**Figure S3. Translating a Single SCF-E Reference Curve into 1000 Virtual Amplification Curves.** ACLs consist of virtual amplification curves that share the same *E*_max_, *E*_min_, and *k* parameters as the reference curve but have unique *N_0.5_* values. This is done by substituting *N_0.5, ref_* from Eq. S5 with *N_0.5, virt_* in Eq. S7. We input 1000 evenly spaced *N_0.5, virt_* values between cycle numbers *N* = 1 to 40, resulting in 1000 virtual amplification curves, each associated with a unique theoretical RNA concentration (*C_RNA, virt_*). Virtual curves with *N_0.5, virt_* < *N_0.5, ref_* (leftmost pink solid curve) correspond to higher RNA concentrations compared to the reference curve, whereas curves with *N_0.5, virt_* > *N_0.5, ref_* (rightmost yellow solid curve) represent lower RNA concentrations than the reference (middle blue solid curve). The efficiency curve for reference and virtual curves is illustrated as a dashed blue line.

**Figure S4. Calculating RNA Template Concentration of the Virtual Amplification Curves.** We relate the known RNA concentration of the reference curve (*C_RNA, ref_*) to the unknown RNA concentrations of the virtual curves (*C_RNA, virt_*) based on their position on the x-axis (cycle number, *N*). In Eq. S10, we define a relationship between the known cycle numbers where PCR efficiency is equal to 0.5 for the reference (*N_0.5, ref_*) and virtual (*N_0.5, virt_*) curves and their respective RNA concentrations (*C_RNA_*). By rearranging Eq. S10, we can calculate the unknown *C_RNA, virt_* using Eq. S12. To perform the calculation, the slope of the line (*m*) in Eq. S13 is determined using a PCR efficiency constant (*E_N_*). As indicated by Eq. S13, an increase in *E_N_* leads to a decrease in *m* (dotted black line), expanding the range of *C_RNA, virt_* in the ACL (from solid triangle to open triangle, and from solid square to open square). This relationship is visually illustrated here with Eq. S8.

**Figure S5. Constructing a Standard Curve from the ACL. (A)** Construction of an amplification curve library (ACL) (Fig. S3). Three *C_RNA, virt_* within the ACL are displayed here (pink, blue, yellow), with the leftmost pink curve representing a higher *C_RNA, virt_* compared to the rightmost yellow curve. **(B)** ACL standard curves (solid line, dashed line, dotted line) are constructed from the *F_N_* of *C_RNA, virt_* (pink, blue, yellow) at a particular cycle number (15, 20, 25). The cycle number chosen for the ACL standard curve corresponds to the cycle number at which unknown RNA concentrations are sampled. In this method, only virtual curves in the exponential or linear phase at the selected cycle number can be included in the standard curve (grey shaded regions in **A** and **B**). This is because the early amplification and plateau regions of multiple virtual curves may overlap at a single cycle number. To ensure that only useable regions are selected for the standard curve, we establish threshold values based on the fluorescence of the reference curve. The upper threshold is set at 1 standard deviation below the *F_N_* at cycle 40, and the lower threshold is set at the 99^th^ percentile of the *F_N_* at cycle 1. For example, as amplification proceeds for a particular *C_RNA, virt_* (pink curve) in **A**, the *F_N_* increases and intersects cycles 15, 20, and 25 at the three points (squares). However, only two of those points are within the valid range of the ACL standard curves in **B.**

**Figure S6. Comparison of SCF-E and SCF Methods of Fitting PCR Amplification Curves.** Fluorescence intensities (11*R_N_*) of bulk RT-qPCR amplification of six concentrations of IAV M gene RNA (Table S2, 10-fold dilutions from 2.62 × 10^4^ to 2.62 × 10^9^ copies/µL) were measured at *N =* 1 to 40. The concentration of each dilution decreases from left to right when viewing the amplification curves. **(A)** Amplification curves generated with the SCF-E model (dotted gray curves) had an average fit of *R*^2^ = 0.9996 across all RNA concentrations (Table S3). **(B)** Amplification curves generated with the SCF model (dotted gray curves) had an average fit of *R*^2^ = 0.9975 across all RNA concentrations (Table S3). **(C)** For a single M gene RNA concentration (2.62 × 10^7^ copies/µL), the SCF-E model provided a better fit (*R^2^* = 0.9996) than SCF (*R^2^* = 0.9972). The difference in fit is most noticeable at the lower (11*R_N_* < 1) and higher (11*R_N_* > 4) fluorescence values, representing the exponential and plateau phases of PCR amplification (magnified insets). All error bars represent one standard deviation from the average fluorescence measured across three technical replicates.

**Figure S7. Setting a PCR Efficiency Constant (*E_N_*) for the ACL.** To calculate *E_N_* at *N = N_0.5_ – 3,* we determined *N_0.5_* of the SCF-E reference curve, which corresponds to *N* where *E_N_* = 0.5 **(A to B)**. *N_0.5_ – 3* represents three cycles prior to *N_0.5_* **(C)**. We recorded the *E_N_* at *N_0.5_ – 3* **(D)**. When *N = N_0.5_ – 3* is inserted into Eq. S3, the term *N_0.5_* cancels out, resulting in a constant value of -3 for (*N – N_0.5_*). The *E_N_* calculated in this step is used in Eq. S13 to determine *m*.

**Figure S8. Linear regression of measured to expected RNA concentrations from Table S4.** We compared cycle numbers (*N*) at which PCR efficiency (*E_N_*, Eq. S3) of the reference curve is used to calculate the constant *m* (Eq. S13), for assigning RNA template concentrations to virtual curves in the ACL. We tested the cycle number at which PCR efficiency is equal to 0.5 (*N_0.5_*) and the five cycles preceding it (*N = N_0.5_ -1 to N = N_0.5_ -5*), to construct six ACLs for comparison. From each ACL, a standard curve was constructed at PCR cycle number 20. We evaluated the dynamic range of each standard curve in Table S4. The ACL standard curve with the widest dynamic range and the highest degree of accuracy (*R*^2^) between measured (*C_RNA, ACL_*) and expected (*C_RNA, expected_*) M gene RNA concentrations was built using *E_N_* at *N = N_0.5_ – 3.* For this ACL, the dynamic range of the standard curve was 4.70 log_10_ RNA copies/μL (Table S4, dynamic range of 2.62 × 10^4^ to 1.31 × 10^9^ copies/μL) with *R*^2^ = 0.990 between *C_RNA, ACL,_* and *C_RNA, expected_*. Therefore, the *E_N_* at *N = N_0.5_ – 3* was used to construct all amplification curve libraries in this work.

**Figure S9. Linear regression of measured to expected RNA concentrations from Table S5.** We compared the dynamic range of ACL standard curves from different cycle numbers (*N* = 17 to 23) to the Ct standard curve. Standard curves were used to convert fluorescence to template concentration for a known dilution series of IAV M gene RNA (Table S2, three replicates of 21 concentrations between 2.62 × 10^4^ to 2.62 × 10^9^ copies/uL). Standard curve dynamic range and degree of accuracy (*R*^2^) between measured (*C_RNA, ACL_*) and expected (*C_RNA, expected_*) M gene RNA concentrations were evaluated in Table S5.

**Figure S10. Variability in M gene RNA Concentrations Measured with ACL Standard Curves by PCR Cycle Number.** We examined variability between standard curves obtained from different ACL cycle numbers using two measures: **(A)** coefficient of variation (*CV* %) and **(B)** percent difference (%). Standard curves were constructed from ACL cycle numbers (*N* = 17 to 23, individual plots) and used to convert fluorescence to RNA concentration (*C_RNA, ACL_*) across 23 IAV M gene RNA concentrations (Table S2). **(A)** *CV* (%) is the ratio of the standard deviation to the mean *C_RNA, ACL_*, and it quantifies the variability *within* replicates of the same M gene RNA concentration. A linear regression (dotted line) was conducted to compare *CV* (%) across M gene RNA concentrations. The results revealed a weak linear relationship (*R^2^* < 0.28) for all cycle numbers. **(B)** Percent difference (*%*) measures the variability *between C_RNA, ACL_* and the expected RNA concentration (*C_RNA, expected_*). Similarly, a weak relationship (*R^2^* < 0.36) was observed between the starting RNA concentration and percent difference (%) for all cycle numbers. To assess the variability in ACL standard curves, a comparison was made with *Ct* standard curves (A and B, first upper left plot), for *CV* (%) and percent difference (%). The analysis indicated no linear response for *CV* (%) (*R^2^* = 0.0236) and a weak linear response for percent difference (%) (*R^2^* = 0.2477) across the M gene RNA concentrations.

**Figure S11: Validating ACL Standard Curves for Multiple Template Sequences.** Standard curves constructed using the **(A)** ACL method or **(B)** Ct method were used to convert fluorescence to template concentration (*C_RNA/DNA, ACL_*) for a known dilution series of three different template sequences: IAV M gene RNA (green/gray dots, ranging from 2.62 × 10^4^ to 2.92 × 10^9^ copies/μL), human MYCN gene DNA (blue squares, ranging from 1.88 × 10^0^ to 1.88 × 10^3^ copies/μL), and soybean Lectin endogene DNA (red circles, ranging from 6.4 x 10^0^ to 4.0 x 10^3^ copies/μL). Each dilution of the templates had varying numbers of technical replicates: 3 for the M gene, 94 for the MYCN, and 18 for the Lectin dataset. The M gene ACL was built with a 10^7^ copies/μL reference curve fitted with either the SCF-E (green dots) or SCF (grey dots) model.

The standard curve from the M gene ACL was constructed at a single cycle number (*N =* 20). The MYCN libraries were built with a 1,875 copies/μL reference curve, and the Soybean ACL was built with an 800 copies/μL reference curve, both fitted using the SCF-E model. The standard curves from these ACLs were constructed at *N =* 27 (MYCN) and *N =* 29 (Lectin). An *R^2^* value quantified the linear response between measured *C_RNA/DNA, ACL_* to expected concentrations (*C_RNA/DNA, expected_*) across template concentrations. Dotted lines indicate a 2-fold change from *C_RNA/DNA, expected_,* and error bars represent one standard deviation from the mean.

**Figure S12. Droplet Quantitative PCR (dqPCR) of the IAV M gene. (A)** SCF-E reference amplification curve generated for 1.71 x 10^2^ cpd of IAV M gene RNA amplified in 50 µm drops. Drop fluorescence intensity (11*R_N_*) was detected at PCR cycle numbers *N* = 1, 13, 16, 19, 22, 25, 28, 40 and displayed as a mean (black dots) and one standard deviation (error bars) (≈1000 or more drops per cycle number). The estimated PCR efficiency (*E_N_*) curve (dashed blue line) used during SCF-E produced a well-fitting reference amplification curve (dotted red line, *R^2^* = 0.9995 (B) dqPCR was validated for a single concentration of M gene RNA (1.71x10^2^ cpd) across three biological replicates. The samples were collected at various cycle numbers: rep 1 (red squares) at *N* = 19 to 21 and rep 2 (blue circles) and rep 3 (green triangles) at *N =* 19 to 22. Drop fluorescence was converted to M gene RNA concentration (*Crna, acl*) within a 2-fold change of the expected mean (dotted lines) for all cycle numbers. Dashed lines indicate a fixed effects model with upper and lower bounds set as the 95% confidence intervals. **(C)** Representative epifluorescence images of 50 μm diameter drops at PCR cycle numbers 1, 19, and 40.

**Figure S13. SCF-E Reference Amplification Curves used in** Figure 3B. Three known concentrations of M gene RNA, 1.71 × 10^1^ cpd (red dots), 1.71 × 10^2^ cpd (blue dots), and 1.71 × 10^3^ cpd (green dots), were amplified in 50 µm drops. Drop fluorescence (11*R_N_*) was detected at multiple PCR cycle numbers (Table S6, note that cycles 19-21 in the 10 cpd group were below background and not included in the table) and converted to M gene cpd using dqPCR (Fig. 3B).

For each experiment run on different days, a new SCF-E reference curve was generated from the amplification of 1.71 × 10^2^ cpd (red dotted line, blue dashed line, green solid line).

**Figure S14. Using dqPCR to Quantify Three IAV M gene RNA Concentrations at Multiple Cycle Numbers.** Three known IAV M gene RNA concentrations, 1.17 × 10^1^ cpd (10 cpd, red circles), 1.17 × 10^2^ cpd (100 cpd, blue crosses), and 1.17 × 10^3^ cpd (1000 cpd, green squares) were amplified in 50 µm drops. Drop fluorescence (11*R_N_*) was detected at multiple PCR cycle numbers (Table S6) and converted to M gene cpd (*C_RNA, ACL_*) using dqPCR. *C_RNA, ACL_* measurements were pooled together and displayed as a single distribution in Fig. 3B. Dashed lines indicate the expected M gene cpd, and dotted lines represent a 2-fold change from the expected mean.

**Figure S15. Analysis of Variance (ANOVA) for dqPCR of Three IAV M Gene RNA Concentrations.** Three known IAV M gene RNA concentrations, 1.17 × 10^1^ cpd (10 cpd plot), 1.17 × 10^2^ cpd (100 cpd plot), and 1.17 × 10^3^ cpd (1000 cpd plot), were amplified in 50 µm drops.

Drop fluorescence (11*R_N_*) was detected at multiple PCR cycle numbers (Table S6) and converted to M gene cpd (*C_RNA, ACL_*) using the dqPCR model. *C_RNA, ACL_* measurements were pooled together and displayed as a single distribution in Fig. 3B. To examine the contribution of error within individual PCR cycle numbers to overall error in the pooled distributions, we performed an Analysis of Variance (ANOVA) test on a random sample of 100 drops from each group. **(A)** The 10 cpd group, quantified at PCR cycle numbers 22 – 25, had mean *C_RNA, ACL_* values that differed by cycle number (ANOVA p-value <0.05). This variability due to cycle number accounted for 8% of the total error in the pooled distribution. **(B)** The 100 cpd group, quantified at PCR cycle numbers 19-22, had mean *C_RNA, ACL_* values that differed by cycle number (ANOVA p-value <0.05). This variability due to cycle number accounted for 59% of the total error in the pooled distribution. (C) The 1000 cpd group, quantified at PCR cycle numbers 16-22, had mean *C_RNA, ACL_* values that differed by cycle number (ANOVA p-value <0.05). This variability due to cycle number accounted for 7% of the total error in the pooled distribution.

**Figure S16. Linear Mixed Effects Model (LME) for dqPCR of Three IAV M Gene RNA Concentrations.** Three known IAV M gene RNA concentrations, 1.17 × 10^1^ cpd (10 cpd, red circle), 1.17 × 10^2^ cpd (100 cpd, blue cross), and 1.17 × 10^3^ cpd (1000 cpd, green triangle), were amplified in 50 µm drops. Drop fluorescence (11*R_N_*) was detected at multiple PCR cycle numbers (Table S6) and converted to M gene cpd (*C_RNA, ACL_*). *C_RNA, ACL_* measurements were pooled together and displayed as a single distribution in Fig. 3B. To further examine the contribution of error within individual PCR cycle numbers to overall error in the pooled distributions, we calculated the percent difference (*%*) (Eq. S14) between the measured *C_RNA, ACL_* values and their expected M gene cpd (*C_RNA, expected_*), across cycle numbers. This analysis informed the validity of pooling conversions measured from multiple PCR cycles in our burst size experiments. Percent differences were compared using a linear mixed effects model (LME) with a 95% confidence interval (CI) (dotted lines). In the LME, we set cycle as the fixed effect variable and *C_RNA, expect_* as the random effect variable. Both variables were log-transformed during the model fit, as replication cycles correspond to RNA copy numbers on a log scale. The fixed effects coefficients described a negative relationship between the slope and percent difference from the expected RNA concentration of -0.85% per cycle number (p-value <0.05, SE = 0.01). While the p-value indicates significance, the magnitude of the relationship is small (∼1%/cycle), maintaining that it is valid to pool dqPCR data from different cycle numbers.

**Figure S17. Using dqPCR to Quantify IAV M gene RNA Concentration, Pooled Cycle Numbers.** Three known IAV M gene RNA concentrations, 1.17 × 10^1^ cpd, 1.17 × 10^2^ cpd, 1.17 × 10^3^ cpd, were amplified in 50 µm drops. Drop fluorescence (11*R_N_*) was detected at multiple PCR cycle numbers (Table S6) and converted to M gene cpd (*C_RNA, ACL_*) using dqPCR. *C_RNA, ACL_* measurements were pooled together and plotted against the expected M gene cpd (*C_RNA, expected_*). Two biological replicates (black circle and green triangle) are shown. The solid line represents a 1:1 relationship between *C_RNA, ACL_* and *C_RNA, expected_*, dashed lines indicate a 2-fold change from the expected mean.

**Figure S18. β-actin mRNA abundance increases with cell concentration.** A bulk RT-qPCR assay was performed to amplify β-actin mRNA from lysed A549 cells after 24 hr of incubation. There was a strong linear relationship between β-actin and cell concentration (dashed black line, *R^2^* = 0.995), validating its use as a reliable marker for the detection of intracellular lysate in drops during our burst size experiments.

**Figure S19. Multiplexed dqPCR for Detection of IAV M gene and Cellular β-actin. (A)** Both M gene RNA (red) and β-actin mRNA (black) were detected by multiplexed dqPCR from H1N1 infection in drops (n=2000). Data is presented in ascending order of RNA concentration, prior to β-actin filtering (indicated by the dashed black line). **(B)** Showing a zoomed-in region of (A), with drop indexes 1900 to 2000.

**Figure S20. SCF-E Reference Curves for Burst Size Replicate Experiments.** For each burst size replicate experiment (Table S8), a new reference amplification curve was constructed for dqPCR (Table S9). To generate these curves, a known concentration of M gene RNA (1.71 x 10^2^ cpd) was amplified in 100 µm drops. Drop fluorescence intensity (11*R_N_*) was measured at PCR cycle numbers *N* = 1, 16, 19, 22. 25, 28, 31 and 40. The 11*R_N_* of drops from each cycle number are shown as the mean (black dots) with error bars representing one standard deviation. SCF-E was used to create a continuous reference amplification curve (orange dashed line) from discontinuous 11*R_N_* measurements, using an estimate for the PCR efficiency (blue dashed line).

**Figure S21. Oseltamivir (OST) treatment of IAV infected cells during drop infections.** MDCK cells infected with IAV H1N1 were encapsulated into 100 µm diameter microfluidic drops. During encapsulation, infected cells were either suspended in standard droplet infection media (+H1N1 -OST) or in droplet infection media treated with 10 µM concentration of oseltamivir (+H1N1 +OST). Mock infected cells encapsulated in standard infection media (-H1N1 –OST) were used as a negative control. M gene abundance (copies / µL) was measured using a bulk RT-qPCR assay from the supernatant of broken drops sampled at 0 and 18 hpi. Each bar represents the pooled data from three technical replicates. Significance stars (*** *p* < 0.001 and **** *p* < 0.0001) obtained by a two-sample Student’s t-test.

**Figure S22.** Best Fit Model to Estimate IAV Burst Size Distributions from Table S10. We used a simulation-based approach to estimate the IAV burst distributions shown in Figure 4D. We considered three distribution models: lognormal (solid gray curve), Poisson (dashed gray curve), and negative-binomial distribution (solid blue or red curve, corresponding to H3N2 and H1N1 distributions, respectively). The parameters for each distribution are provided in Table S10. Based on the analysis, burst sizes were found to best fit a negative binomial distribution.

**Figure S23. Cell Loading During Burst Size Replicate Experiments**. **(A)** Images of 100 µm drops re-injected into a microfluidic device were captured at the beginning and end of each burst size replicate experiment to monitor cell loading. Drops containing a single cell are indicated by blue arrows. **(B)** Across all H3N2 and H1N1 replicate experiments, cell loading was determined by counting the percentage of drops containing cells (mean = 25% ± 4%). Cell loading in drops was assumed to follow a Poisson distribution. For a population of drops with 25% containing cells, the estimated Poisson mean (λ) is 0.29 cells/drop. In this case, ≈75% of drops are empty, ≈21.5% of drops contain one cell, ≈3% contain two cells, and ≈0.5% of drops contain three or more cells. The bar graph displays the drop counts for the replicates from left to right, which are 753, 679, 538, 719, 774, and 582.

**Table S1. RT-qPCR Template and Primer Sequences**.

All sequences are listed in the 5’ to 3’ direction. The M gene template control was constructed as linear dsDNA containing a T7 promoter (<u>underlined</u>), forward and reverse primer binding sites (*italicized*), a probe binding site (**bold**), and a partial M gene sequence. The M gene dsDNA was *in vitro* transcribed prior to use as a positive control. The M gene TaqMan probe carried a FAM fluorophore with a BHQ1 quencher. The β-actin template control was constructed as a dsDNA plasmid containing forward and reverse primer binding sites (*italicized*), and a probe binding site (**bold**), and a partial β-actin mRNA sequence. The β-actin TaqMan probe carried a Cy5 fluorophore with a BHQ2 quencher.

**Table S2. IAV M Gene RNA Concentrations Amplified by Bulk RT-qPCR and Used to Validate dqPCR.**

**Table S3. SCF-E and SCF Curve Fitting Parameters for Figure S6.**

**Table S4. Dynamic Range of ACL Standard Curves Built with Different *E_N_* Constants.** We compared cycle numbers (*N*) at which PCR efficiency (*E_N_*) of the reference curve is used to calculate the constant *m* (and *E_N_*). This constant determines the RNA template concentration assigned to virtual curves in the ACL. *E_N_* is calculated using Eq. S3 and *m* is calculated using Eq. S13. We used the cycle number at which PCR efficiency is equal to 0.5 (*N_0.5_*) and the five cycles preceding it (*N = N_0.5_ -1 to N = N_0.5_ -5*), to construct six ACLs for comparison. From each ACL, a standard curve was generated at PCR cycle number 20. This standard curve was then used to convert fluorescence (*F_N_*) to template concentration (*C_RNA_*) for a known dilution series of M gene RNA (Table S2, three replicates of 21 concentrations between 2.62 × 10^4^ to 2.62 × 10^9^ copies/µL). The minimum (C_RNA, min_) and maximum (C_RNA, max_) concentrations in the dilution series, whose measured RNA concentration fell within a 2-fold change of the expected concentrations, marked the dynamic range of each standard curve. The *R^2^* value quantifies the linear regression (Fig. S8) between the measured (*C_RNA, ACL_*) and expected (*C_RNA, expected_*) RNA concentrations. The ACL built with *E_N_* at *N = N_0.5_ – 3* produced a cycle 20 standard curve with the highest dynamic range and the greatest degree of accuracy (*R^2^*). Therefore, an offset of -3 cycles was used to construct all ACLs in this work.

**Table S5. Dynamic Range of ACL Standard Curves from Different ACL Cycle Numbers.** We compared the dynamic range of standard curves created using the amplification curve library (ACL) method to that of the cycle threshold (*Ct*) method. Standard curves were used to convert fluorescence to template concentration for a known dilution series of IAV M gene RNA (Table S2, three replicates of 21 concentrations between 2.62 × 10^4^ to 2.62 × 10^9^ copies/µL). In the ACL method, multiple standard curves were constructed from individual cycle numbers (*N* = 17 to 23). On the other hand, in the *Ct* method, a single standard curve is built from multiple *Ct* values, with one *Ct* value corresponding to each concentration of M gene RNA. The dynamic range of each standard curve was determined by identifying the minimum (*C*_RNA, min_) and maximum (*C*_RNA, max_) concentrations within the dilution series that yielded measured RNA concentration falling within a 2-fold change of the expected concentrations. The *R^2^* value quantifies the linear response (Fig. S9) between the measured (*C_RNA, ACL_*) and expected (*C_RNA, expected_*) RNA concentrations.

**Table S6. Using dqPCR to Quantify IAV M Gene RNA Concentrations at Multiple Cycle Numbers.**

Three known IAV M gene RNA concentrations, 1.71 × 10^1^ cpd (10^1^), 1.71 × 10^2^ cpd (10^2^), and 1.71 × 10^3^ cpd (10^3^), were amplified in 50 µm drops. Drop fluorescence (11*R_N_*) was detected at multiple PCR cycle numbers and converted to M gene cpd (*C_RNA, AC_*_L_) using dqPCR. *C_RNA, ACL_* measurements from different cycle numbers were pooled together and presented as a single distribution in Figure 3B. To assess the variability between individual cycle numbers and the pooled distribution, we analyzed the standard deviation of the mean *C_RNA, AC_*_L_ and the coefficient of variation (*CV %*) across the number of sampled drops (*n*) at each *N*.

**Table S7. Maximum Likelihood Estimates for a Gaussian Mixture Model (GMM) Used to Extract Individual Distributions from Mixed Droplet Detection Data.**

To use dqPCR for measuring IAV burst size in a heterogeneous population of single-cell infections, we validated the method’s ability to isolate individual distributions from a sample containing multiple M gene RNA concentrations. We applied a Gaussian mixture model (GMM) on dqPCR data obtained from three known IAV M gene RNA concentrations: 1.17 × 10^1^ cpd, 1.17 × 10^2^ cpd, and 1.17 × 10^3^ cpd, mixed together in a single sample (data shown in Figure 3E). Model assumptions are described in **SI R7**. In the model, *x_i_* represents the number of M gene RNA copies (cpd). Each Gaussian has a different weight (*w_i_*) corresponding to the probability that a random drop in the mixture has *x_i_* copies, a predicted mean (*B_i_*, representing measurement bias) and variance (*σ^2^_i_*, representing measurement noise).

**Table S8. Burst Size Replicate Experiments.**

IAV burst sizes were measured from thousands of individual drops sampled at cycle numbers *N* = 16, 19, 22, 25, and 28 in three biological replicates. The measurements were used to determine the number of IAV M gene RNA and cellular β-actin mRNA (cpd). Drops containing high β-actin cpd (Fig. S19) were assumed to contain non-packaged, intracellular, M gene RNA and were subsequently filtered from the final burst size distributions (Filtered Drops, *SI Appendix* R8). We present the average measured burst size for each replicate, along with the interquartile range (IQR) that represents the lower 25^th^ and upper 75^th^ percentile of the distributions. The heterogeneity of each distribution was examined using the Gini coefficient (*G*). If *G* = 0, the distribution is completely homogeneous, meaning that each cell produces the same number of viruses. Conversely, if *G* = 1, the distribution is completely heterogeneous, where one cell produces all the viruses. Thus, a higher *G* corresponds to a more heterogeneous distribution.

**Table S9. dqPCR Parameters Used in Burst Size Replicate Experiments.**

The parameters used to create reference standard curves using SCF-E (Fig. S20, orange dotted curves) for each burst size experiment.

**Table S10. Best Fit Model to Estimate IAV Burst Size in** Fig. 4D.

We used a simulation-based approach to estimate the IAV burst distributions shown in Figure 4D. We considered three possible distributions: lognormal, Poisson, and negative-binomial, all with unknown parameters. First, we simulated the viral burst size based on the assumed distribution. For each simulated value of burst size *x*, we introduced measurement noise by assuming a log-normal distribution. The mean of the log-normal distribution was determined by the bias function *σ(x)*, and the standard deviation was determined by *σ(x)*. Next, we computed a density function using a kernel density estimation with a Gaussian kernel to represent the resulting distribution. We then used this density function to calculate the log-likelihood of the observed data. We estimated the parameters for each distribution and for each dataset (H3N2 or H1N1) by maximizing the likelihood of observations reported in Fig. 4D.

