## Supplementary Information for "Influenza A Viral Burst Size from Thousands of Infected Single Cells Using Droplet Quantitative PCR (dqPCR)"

**Table of Contents**

- Supplementary Materials and Methods M1 to M9
- Supplementary Results R1 to R10
- Supplementary Figures S1 to S23
- Supplementary Tables S1 to S10
- References for Supplementary Information

#### Supplementary materials and methods

**(M1) RT-qPCR Template and Primer Sequences.** Detection of IAV and host cells was facilitated by reverse transcription quantitative polymerase chain reaction (RT-qPCR) of IAV matrix protein genomic RNA (M gene) and cellular β-actin mRNA. M gene RNA was amplified with forward/reverse primers and a Taqman probe (FAM/BHQ1) that target a highly conserved region shared across IAV strains (1) (Table S1). β-actin mRNA was amplified with forward/reverse primers and a Taqman probe (Cy5/BHQ2) shown to be robust against multiple eukaryotic cell lines (2) (Table S1). All primers and probes were purchased from Eurofins Operon as 100 µM stocks. Known concentrations of the template sequences were used as positive controls for RT-qPCR validation throughout this work. The M gene control was first constructed as linear dsDNA (IDT) containing a T7 promoter (underlined), forward and reverse primer binding sites (italicized), and a probe binding site (bold) (Table S1). M gene RNA was *in vitro* transcribed (IVT) from the ssDNA gblock using the MEGAscript T7 RNA Synthesis Kit (Ambion, #AM1333), and purified over a GE Illustra Sephadex G-50 NICK column. The β-actin control (Table S1) was constructed as a dsDNA plasmid (pCAG-mGFP-Actin, Addgene #21948), also containing forward and reverse primer binding sites (italicized) and a probe binding site (bold), which was directly amplified by PCR. The concentrations (copies/µL) of template sequence controls in working stocks were quantified with a NanoDrop spectrophotometer. For droplet experiments, template sequence concentrations in copies per drop (cpd) were calculated by multiplying copies/µL by the volume (µL) of a 50 µm ($6.54 \times 10^{-5}$ µL) or 100 µm (or $5.24 \times 10^{-4}$ µL) diameter drop.

**(M2) Microfluidic Device Fabrication.** Microfluidic devices were fabricated in polydimethylsiloxane (PDMS) (Sylgard 184) using soft lithography. Three device geometries were designed using AutoCAD (Autodesk): a flow-focusing device for encapsulation of infected cells in 100 µm drops, a split-and-merge device used to isolate replicated virus from the host cell and merge with RT-qPCR mix (3), and a flow-based imaging device (4) for drop fluorescence detection. Device master molds were fabricated by patterning SU-8 photoresist (Microchem, SU-8 3050) on silicon wafers (University Wafer, #447) with photolithography. PDMS was prepared at a

10:1 mass ratio of polymer to cross-linking agent and poured onto the device master molds. Air was purged from the uncured PDMS by placing the filled mold in a vacuum chamber for at least 1 hr. The PDMS was cured in an oven at 55 °C for 24 hrs and then removed from the mold with a scalpel. Inlet and outlet ports were punched into PDMS slabs with a 0.75 mm diameter biopsy punch (EMS Rapid-Core, Electron Microscopy Sciences). The split-and-merge device was comprised of two layers bonded together after plasma treatment (Harrick Plasma, PDC-001) for 30 s at medium power and 700 mTorr oxygen pressure. The double-layer PDMS split-and-merge device and single-layer PDMS flow-focusing device were each bonded to 2 × 3 in glass slides (VWR micro slides, #48382-179) after plasma treatment for 60 s at high power and 400 mTorr oxygen pressure. Microelectrodes were embedded in the split-and-merge device by injecting molten solder (Indium Corporation of America, #53307) into the electrode channels, and terminated with a pin terminal (Phoenix Contact, #1945151). Both devices were made hydrophobic by flowing Aquapel (Pittsburgh Glass Works) through the channels, followed by blowing the channels with air passed through a 0.2-µm filter (GVS ABLUO 25 mm, Fisher Scientific) before baking the devices in an oven at 55 °C for 1 hr for further drying.

**(M3) Bulk RT-qPCR.** Bulk RT-qPCR reactions were completed using the SuperScript III Platinum One-Step RT-qPCR kit (Invitrogen 11732020) with a final reaction volume of 25 µL. Working stocks of M gene primers and FAM TaqMan probe (Table S1) were prepared at 25 µM and 10 µM, respectively. The bulk RT-qPCR master mix contained final concentrations of 400 nM primers, 200 nM probe, 0.05 µM ROX reference dye, 2.0 mM $MgSO_4$, 1.0% w/v Tween-20 (Calbiochem 655204-100mL), 0.8 µg/µL BSA (Fisher BP675-1), 1.0 M betaine (Sigma B0300-1VL), 0.32 U/µL SUPERase RNase Inhibitor (Invitrogen AM2694), and 1 µL of SuperScript III RT/Platinum Taq mix (Invitrogen 11732020). The additives (Tween-20, BSA, betaine) increase stability of dqPCR reactions, as determined by our previously published protocol (5). Thermocycling was performed using a standard qPCR machine (QuantStudio 7, Applied Biosystems) with the following conditions: 1 cycle for 30 mins at 60 °C, 1 cycle for 2 mins at 95

°C, and 40 cycles between 15 s at 95 °C and 1 min at 60 °C. Reference amplification curves for the bulk RT-qPCR assay were constructed from a dilution series of M gene IVT RNA (Table S2).

**(M4) Droplet Quantitative PCR (dqPCR) of Isolated Progeny Viruses.** A multiplexed dqPCR assay was used to amplify both IAV M gene RNA and cellular β-actin mRNA in drops containing progeny virus. The dqPCR master mix contained the same reagent concentrations as the bulk RT-qPCR, except for an increased concentration of ROX reference dye at 0.15 μM. Working stocks of the M gene primers and FAM TaqMan probe, as well as the β-actin primers and Cy5 TaqMan probe (Table S1), were prepared at concentrations of 25 μM and 10 μM, respectively. Droplets containing infected cells were re-injected into a split-and-merge device (3) at a flow rate of 500 μL/hr with oil (3.0 wt% RAN surfactant in HFE 7500) flowing at 2000 μL/hr to space them apart. In the device, a split junction was used to isolate progeny viruses from the host cell. Approximately 1/8$^{th}$ volume of the drop containing progeny virus was split and sent to the merge junction. To separate split drops before the merge junction, a second spacer oil (3.0 wt% RAN surfactant in HFE 7500) was injected into the device at a flow rate of 500 μL/hr. At the merge junction, the RT-qPCR master mix was injected into the split drop at a flow rate of 438 μL/hr. Merging was facilitated by an embedded microelectrode that provided a constant 25 kHz, 200 V square wave signal to destabilize the drop interface and allow merging with the RT-qPCR mix. The microelectrode was connected to a high voltage amplifier (Trek Model, #2220-CE) controlled by a custom LabVIEW program. Merged drops were collected in PCR tubes for 2.5 min each to sample approximately 20 μL of drops. PCR tubes were stored on a cold block until ready for thermocycling. Drops were thermocycled in a standard qPCR machine (QuantStudio 7, Applied Biosystems) with the following conditions: 1 cycle for 30 min at 60 °C, 1 cycle for 2 min at 95 °C, and 40 cycles between 15 sec at 95 °C and 1 min at 60 °C. Drops were removed from the thermocycler at cycle numbers $N = 1$ and $N = 40$, to measure baseline and terminal fluorescence, as well as multiple intermediate cycle numbers falling within the exponential to linear region of PCR amplification curves.

Reference amplification curves for the dqPCR assay were constructed by encapsulating $10^6$ copies/µL of M gene IVT RNA and β-actin plasmid (Table S1) into 50 µm drops using a flow-focusing device, which corresponds to $10^2$ cpd of M gene and β-actin. Drops containing M gene or β-actin template sequence controls were collected in 3 mL syringes for 36 min to generate 600 µL of drops and kept on ice until thermocycling with the above conditions. The method for constructing reference amplification curves from dqPCR of template sequence controls is further described in **SI M8.**

**(M5) Drop Fluorescence Imaging.** As a positive control, thermocycled drops were imaged to capture five fields of view on an inverted epifluorescence microscope (Nikon Ti2-E) at 10× magnification (NA 0.3). Brightfield and fluorescence images were captured with a sCMOS camera (Hamamatsu, ORCA-Flash 4.0 v3) for each reporter fluorescence, FAM TaqMan for M gene RNA (FITC channel), Cy5 TaqMan probe for β-actin RNA (Cy5 channel), as well as for the ROX reference dye (Texas Red channel). Drop fluorescence intensities are reported as $\Delta R_N$, which is the ratio of the M gene (FAM) or β-actin (Cy5) reporter florescence to the reference dye fluorescence (ROX) at each sampled cycle number (*N*), normalized to the baseline at *N* = 1. Here, $R_N = (F_{N,\ FAM}$ or $F_{N,\ Cy5})/F_{N,\ ROX}$ and $\Delta R_N = R_N - R_{N=1}$.

**(M6) Drop Flow-based Fluorescence Detection.** Thermocycled drops were injected into a continuous-flow microfluidic detection device (4) at a flowrate of 200 µL/hr, along with fluorinated spacer oil (HFE 7500) injected at a flowrate of 800 µL/hr. Drop fluorescence intensity ($\Delta R_N$) was detected in high-throughput using a Nikon Ti-U inverted microscope custom-modified with three lasers, a set of dichroic mirrors, and two photomultiplier tubes (PMTs, Hamamatsu H10723-20). The beams of the 22 mW 488 nm laser (Thorlabs MCLS1), 25 mW 561 nm laser (Cobolt Jive), and 20 mW 642 nm laser (Thorlabs MCLS1) were aligned and coupled into the backport of the microscope where they were focused to a spot by the 40× objective (NA 0.60). The flowing of drops across the laser spot resulted in fluorescence detection by three PMTs split into three channels using dichroic and bandpass filters, Ch1: 520/40 nm (green), Ch2: 600/40 nm (red), and

Ch3: 670/10 nm (far-red). A field programmable gate array (FPGA, National Instruments NI-7852R) was used to control the PMT gains and record fluorescence measurements using LabVIEW 2015. A custom MATLAB (R2020a) script was used to process and analyze the drop fluorescence detection data.

**(M7) Cycle Threshold (*Ct*) Method of Creating Standard Curves for RT-qPCR.** In standard bulk RT-qPCR, the nucleic acid template concentration of an unknown sample is determined by comparing its cycle threshold (*Ct*) value to a standard curve made from a dilution series of known template concentrations. The standard curve follows Eq. S1:

$$\boldsymbol{Ct = m * log_{10}(C_{RNA}) + b}$$

**(Eq. S1)**

Cycle threshold (*Ct*) is the PCR cycle number at which template amplification fluorescence rises above background fluorescence, $C_{RNA}$ is the known template concentration, *m* is the slope of the *Ct* versus $\log_{10}(C_{RNA})$ line, and *b* is the y-intercept of the line.

This method assumes that PCR efficiency at the *Ct* value ($E_{Ct}$) is constant, regardless of starting template concentration. An optimized PCR assay should have an $E_{Ct}$ between 90 – 110% for accurate quantification of template concentration. $E_{Ct}$ is calculated using *m* of the *Ct* versus $\log_{10}(C_{RNA})$ line, as shown in Eq. S2:

$$\boldsymbol{E_{Ct} = 10^{\frac{-1}{m}} - 1}$$

**(Eq. S2)**

For an ideal assay with $E_{Ct}$ = 100%, the Ct values of a 10-fold dilution series are spaced approximately 3.3 cycles apart ($m \cong -3.3$).

**(M8) Sigmoidal Curve Fitting using PCR Efficiency (SCF-E).** To track RNA amplification during dqPCR, we reconstructed continuous RT-qPCR amplification curves from discontinuous drop fluorescence measurements, as shown in Fig. 2A. Amplification curves can be fit to discontinuous drop fluorescence measurements using a sigmoidal curve fitting (SCF) model. Here, we employ SCF that relates reaction efficiency to fluorescence intensity to accurately model the shape of RT-qPCR amplification curves. This method is called “Sigmoidal Curve Fitting Using PCR Efficiency,” or SCF-E. We used SCF-E to generate reference amplification curves of a known RNA template concentration during dqPCR.

RT-qPCR amplification curves can have an asymmetric shape due to a decrease in PCR reaction efficiency after each cycle number. At early cycles, PCR efficiency is near 100%, relating to an exact doubling of nucleic acid template sequence at each cycle, and approaches 0% at later cycles as PCR reagents become consumed (6). This relationship leads to a steeper slope at the start of the curve during exponential amplification and a flatter slope near the final plateau phase (7). To model asymmetric amplification curves from fluorescence data alone, SCF methods typically set a limit on the number of cycles that are fit (8) or add asymmetric fitting terms (9,10). Alternative methods first model PCR efficiency with a sigmoidal (11) or bilinear (7) equation, and use this to construct a fluorescence amplification curve. In our SCF-E model, asymmetric RT-qPCR amplification curves are fit to experimental fluorescence measurements using a four-parametric sigmoid function that includes a PCR efficiency parameter, $E_N$, as shown in Eq. S3:

$$\boldsymbol{E_N = \frac{E_{max}}{1 + e^{\left(\frac{N - N_{0.5}}{k}\right)}} + E_{min}}$$

**(Eq. S3)**

$E_N$ is the PCR efficiency at cycle $N$, $E_{max}$ is the maximum PCR efficiency in the range of 0.9 to 1, $E_{min}$ is the minimum PCR efficiency in the range of 0 to 0.1, and $N_{0.5}$ is the cycle at which $E_N$ equals 0.5. The parameter $k$ determines the shape of the amplification curve, with larger values flattening the curve and smaller values steepening the curve. An example of the PCR efficiency curve is illustrated in Fig. S2 (blue dashed curve).

During RT-qPCR, RNA amplification is detected with a complementary fluorescent probe during thermocycling. The rate of fluorescence accumulation during RT-qPCR is determined by the reaction efficiency, as described by Eq. S4:

$$\boldsymbol{F_{N+1} = F_N(1 + E_N)}$$

**(Eq. S4)**

$F_N$ is the fluorescence intensity of the amplification curve at cycle $N$, and $F_{N+1}$ is the fluorescence intensity of the amplification curve at cycle $N+1$. The fluorescence amplification curve is also illustrated in Fig. S2 (blue solid curve).

Eqs. S3 and S4 are combined to yield Eq. S5, the **SCF-E model**:

$$\boldsymbol{F_{N+1} = F_N\left(1 + \frac{E_{max}}{1 + e^{\left(\frac{N-N_{0.5}}{k}\right)}} + E_{min}\right)}$$

**(Eq. S5)**

To generate an RT-qPCR amplification curve of $F_N$ at all PCR cycle numbers ($N$ = 1 to 40), we vary the efficiency parameters in Eq. S3 ($E_{max}$, $E_{min}$, $N_{0.5}$, and $k$) to find the best fit to our experimental $F_N$ values, measured at only a few PCR cycle numbers. Thirty values of each of the four efficiency parameters are used to test $8.1 \times 10^5$ ($30^4$) curve fits. The set of parameters that yield the best fit (highest $R^2$ value) to experimental $F_N$ measurements are used to construct the RT-qPCR amplification curve. An initial input for $F_N$ of any non-zero integer is chosen for the fit; here, we used $F_N = 1$. A calibration factor ($CF$), shown in Eq. S6, is used to normalize $F_N$ at $N$ =

40 in the SCF-E model curve ($F_{endpoint,\ model}$) to the experimentally measured $F_N$ at $N$ = 40 ($F_{endpoint,\ measured}$):

$$CF = F_{endpoint,measured}/F_{endpoint,model}$$

**(Eq. S6)**

**(M9) Amplification Curve Library (ACL) Method of Constructing Standard Curves for RT-qPCR.** We generated a high-resolution dilution series of 1000 RNA template concentrations to construct standard curves for conversion of drop fluorescence to RNA concentration. This is performed by first generating a single reference amplification curve for a known IAV M gene RNA concentration ($10^2$ cpd) (Fig. 2A), that is fit using the SCF-E model (Eq. S5). The reference amplification curve is then translated along its *x*-axis by varying the SCF-E $N_{0.5}$ parameter 1000 times, between cycles 1 to 40, to build what we call an "Amplification Curve Library" (ACL) of 1000 virtual curves (Fig. 2B, plot with subset of 10 curves). The ACL is then used to construct a virtual standard curve from a single PCR cycle number (Fig. 2C). This method for generating standard curves is described in detail below in three sections.

***(1) Translating a Single SCF-E Reference Curve into 1000 Virtual Amplification Curves:*** The ACL is constructed by expanding a single reference amplification curve, fit with Eq. S5, into 1000 virtual curves. This is done by substituting $N_{0.5,\ ref}$, fit from Eq. S5, with $N_{0.5,\ virt}$, to make Eq. S7:

$$F_{N+1} = F_N\left(1 + \frac{E_{max}}{1 + e^{\left(\frac{N - N_{0.5,virt}}{k}\right)}} + E_{min}\right)$$

**(Eq. S7)**

The translation of the reference curve into virtual curves using Eq. S7, is illustrated in Fig. S3. Here, $F_N$ and $E_N$ are plotted on the left and right *y*-axes, $N$ on the *x*-axis (Fig. S3). The solid blue curve is an example of a reference curve fit using the SCF-E model (Eq. S5) and the dashed blue curve is the corresponding PCR efficiency curve, as previously described (Fig. S2). The reaction efficiency parameters in Eq. S7 ($E_{max}$, $E_{min}$, $k$) are taken from the reference curve fit (Eq. S5) and are constant across all the virtual curves. In the ACL, each virtual curve is defined by a different cycle number at which PCR reaction efficiency equals 0.5 ($N_{0.5,\ virt}$). We input 1000 evenly spaced $N_{0.5,\ virt}$ values, between cycle numbers $N$ = 1 to 40, into Eq. S7 to produce 1000 unique virtual curves. Two virtual curves are displayed in Fig. S3, to illustrate that $N_{0.5,\ virt} < N_{0.5,\ ref}$ (pink curve) is associated with virtual RNA concentrations ($C_{RNA,\ virt}$) greater than the reference ($C_{RNA,\ ref}$). Conversely, if $N_{0.5,\ virt} > N_{0.5,\ ref}$ (yellow curve) then $C_{RNA,\ virt} < C_{RNA,\ ref}$.

***(2) Calculating RNA Template Concentration of the Virtual Amplification Curves:***

$C_{RNA,\ virt}$ is determined by its *x*-axis distance from $C_{RNA,\ ref}$. Since the *x*-axis of the ACL is cycle number, we can relate the virtual and reference curves using the cycle numbers at which PCR efficiency equals 0.5, or $N_{0.5,\ virt}$ and $N_{0.5,\ ref}$. The $N_{0.5,\ virt}$ is a fixed array of 1000 intervals between cycle numbers $N$ = 1 to 40, as previously mentioned (Eq. S7), while $N_{0.5,\ ref}$ is a single cycle number on the reference curve (fit from Eq. S5). The relationship between an amplification curve's $N_{0.5}$ value and its $C_{RNA}$ value yields a line that follows Eq. S8 below, which is analogous to the *Ct* method (Eq. S1), with $N_{0.5}$ used in place of *Ct*.

$$\boldsymbol{N_{0.5} = m * log_{10}(C_{RNA}) + b}$$

**(Eq. S8)**

We expand this equation for $N_{0.5,\ virt}$ and $N_{0.5,\ ref}$ and take the difference with Eq. S9:

$$\boldsymbol{N_{0.5,virt} - N_{0.5,ref} = [m * log_{10}(C_{RNA,virt}) + b] - [m * log_{10}(C_{RNA,ref}) + b]}$$

**(Eq. S9)**

Eq. S9 is rearranged in Eqs. S10-11 in order to solve for $C_{RNA,\ virt}$ in Eq. 12:

$$\boldsymbol{log_{10}\left(C_{RNA,virt}\right) = log_{10}\left(C_{RNA,ref}\right) + \frac{N_{0.5,virt} - N_{0.5,ref}}{m}}$$

**(Eq. S10)**

$$\boldsymbol{f = log_{10}\left(C_{RNA,ref}\right) + \frac{N_{0.5,virt} - N_{0.5,ref}}{m}}$$

**(Eq. S11)**

$$\boldsymbol{C_{RNA,virt} = 10^{f}}$$

**(Eq. S12)**

Note that the value of $C_{RNA,\ virt}$ varies with $m$ from Eq. S8 when used in Eq. S11. To calculate $m$, Eq. S2 is rearranged into Eq. S13:

$$\boldsymbol{m = \frac{1}{log_{10}(E_N + 1)}}$$

**(Eq. S13)**

$m$ is dependent upon $E_N$ of the reference curve at a single cycle number, calculated with Eq. S3. As $E_N$ increases, $m$ decreases, and the range of $C_{RNA,\ virt}$ values in the ACL widens (Fig. S4). To determine the $E_N$ constant used to calculate $m$ in Eq. S13, we empirically compared standard curves from libraries built with different $E_N$ values **(SI R3)**. Briefly, $E_N$ corresponding to $N = N_{0.5} - 3$ was used to determine virtual RNA concentration for all ACLs used in this work.

***(3) Constructing a Standard Curve from the ACL:*** To determine the RNA concentrations of unknown samples based on their fluorescence intensity, we utilize the ACL (Fig. S5A) to construct standard curves that relate $C_{RNA}$ to $F_N$ (Fig. S5B). The standard curve is constructed by selecting one PCR cycle number in the ACL (Fig. S5A, vertical line) and relating

$F_N$ at that cycle number to $C_{RNA, virt}$ of curves intersecting the line. The usable region of the standard curve (Fig. S5B, grey box) generally corresponds to the exponential to linear region of the virtual amplification curves (Fig. S5A, grey box). Outside of these regions, $F_N$ has either not risen above baseline fluorescence at early cycle numbers or has reached saturation at late cycle numbers. Threshold values are set for $F_N$ to define these usable regions, where the upper threshold is set at 1 standard deviation below the $F_N$ at $N = 40$, and the lower threshold is set as the 99$^{th}$ percentile of the $F_N$ at $N = 1$, similar to the thresholds generally set by commercial qPCR machines.

### Supplementary results

**(R1) Comparing M Gene Abundance During Bulk and Drop Infections.** To demonstrate that encapsulation in drops does not affect IAV infection dynamics (12) during our burst size experiments, we compared the M gene RNA abundance in drop and bulk infections using a bulk RT-qPCR assay (Fig. S1). For bulk infections, we collected both the supernatant and cellular monolayer from each well and prepared them for bulk RT-qPCR. The 0 hpi bulk samples were frozen overnight at -80 °C while the 18 hpi bulk samples were incubated overnight at 37 °C. Preparation of drop infections for bulk RT-qPCR involved freezing a 300 µL sample of drops at -80 °C for 30 mins to break the emulsion. 200 µL of the broken emulsion was collected for further processing. Collected bulk and drop infections were clarified by centrifugation at 500 × *g* for 5 min, and the resulting supernatant was sampled for RT-qPCR targeting the IAV M gene. For both strains, there was greater virus production in drops compared to bulk infection (Fig. S1, 18 hpi), when measured with a two-sample Student's t-test ($p << 0.05$). The relative increase in virus production, between drop and bulk infection, was 6.8× for H3N2 and 2.4× H1N1 infections. On average, H3N2 production in drops was 5.0× greater than H1N1 ($p << 0.05$).

**(R2) Validating SCF-E with Bulk RT-qPCR of IAV M Gene.** Using bulk RT-qPCR amplification of the IAV M gene across five orders of magnitude (Table S2, 10 fold dilutions of $2.62 \times 10^4$ to $2.62 \times 10^9$ copies/µL), we validated the use of our SCF-E model (Fig. S6A) compared to the standard SCF model (Fig. S6B) (8,13). SCF-E fit parameters are listed in Table S3. The SCF-E model provided a better fit across all six RNA concentrations compared to SCF (Table S3, $R^2$ of SCF-E vs. SCF). The differences in fit were most pronounced in the exponential and plateau regions of RT-qPCR amplification curves, as demonstrated in Fig. S6C for 2.62 × $10^7$ copies/µL (magnified insets).

**(R3) Setting a PCR Efficiency Constant ($E_N$) for the ACL.** To calculate $C_{RNA,\ virt}$ in the ACL, we needed to determine the $E_N$ constant used to calculate $m$ in Eq. S13. $E_N$ is the PCR efficiency at each cycle number ($N$ = 1 to 40) in the SCF-E reference curve (Eq. S5) and $m$ is the slope of the $C_{RNA}$ vs. $N_{0.5}$ line (Eq. S8). To determine which of the 40 possible reference curve $E_N$ values to use in Eq. S13, we constructed ACLs with different $E_N$ values and compared the dynamic range of their standard curves (Table S4). A schematic of the iterative process for determining $E_N$ at different $N$ values is displayed in Fig. S7.

We first tested $E_N$ at $N = N_{0.5} – 0$, using a SCF-E reference curve of $10^7$ copies/µL of IAV M gene RNA amplified by bulk RT-qPCR. This yielded $E_N$ = 0.514 and corresponding $m$ = 5.55 (Table S4). These values were used to build an ACL, from which we constructed a standard curve at cycle 20. The standard curve was used to convert $F_N$ to $C_{RNA}$ for a known dilution series of IAV M gene RNA (Table S2, 3 replicates of 21 concentrations between 2.62 × $10^4$ to 2.62 × $10^9$ copies/µL) amplified by bulk RT-qPCR. To determine the accuracy of the standard curve, we performed a linear regression between measured ($C_{RNA,\ ACL}$) and expected ($C_{RNA,\ expected}$) RNA concentrations (Fig. S8, $N = N_{0.5} – 0$, $R^2$ = 0.886). Samples in the dilution series with a $C_{RNA,\ ACL}$ value within 2-fold change of $C_{RNA,\ expected}$ were included in the dynamic range of the standard curve (Table S4). For $E_N$ at $N = N_{0.5} – 0$, the dynamic range of the cycle 20 standard curve was small, a value of 1.60 $\log_{10}$ RNA copies/µL (Table S4, dynamic range of 3.28 × $10^6$ to 1.31 × $10^8$ copies/µL). Thus, to widen the dynamic range, we tested higher $E_N$ values at five cycles preceding $N_{0.5}$ (Table S4, $N = N_{0.5} – 1$, $N = N_{0.5} – 2$, $N = N_{0.5} – 3$, $N = N_{0.5} – 4$, and $N = N_{0.5} – 5$). The ACL standard curve with the widest dynamic range and highest degree of accuracy ($R^2$)

between $C_{RNA,\ ACL}$ and $C_{RNA,\ expected}$ was built with $N = N_{0.5} – 3$, where $E_N$ = 0.745. For this ACL, the dynamic range of the standard curve was 4.70 $\log_{10}$ RNA copies/µL (Table S4, dynamic range of $2.62 \times 10^4$ to $1.31 \times 10^9$ copies/µL) with $R^2$ = 0.990 between $C_{RNA,\ ACL}$ and $C_{RNA,\ expected}$. Therefore, the $E_N$ corresponding to $N = N_{0.5} – 3$ was used to build all amplification curve libraries in this work.

**(R4) Validating ACL Standard Curves for Multiple PCR Cycle Numbers.** To pool $C_{RNA,\ ACL}$ values sampled at multiple PCR cycle numbers, we validated that ACL standard curves converted $F_N$ to $C_{RNA,\ ACL}$ within an acceptable error range of 2-fold change from $C_{RNA,\ expected}$, at any cycle number. This was done using a known dilution series of IAV M gene RNA amplified by bulk RT-qPCR (Table S2, 3 replicates of 21 concentrations between $2.62 \times 10^4$ to $2.62 \times 10^9$ copies/µL). Each ACL was built from a SCF-E reference curve of $2.62 \times 10^7$ M gene copies/µL. Standard curves were constructed at ACL $N$ = 17 to 23, and used to converted $F_N$ to $C_{RNA,\ ACL}$ for the above dilution series. These standard curve cycle numbers were chosen as they typically fall in the linear region of our M gene reference curves. To determine how well $C_{RNA,\ ACL}$ matched to $C_{RNA,\ expected}$, we performed a linear regression on these values (Fig. S9). Samples in the dilution series whose $C_{RNA,\ ACL}$ fell within a 2-fold change of $C_{RNA,\ expected}$ were included in the dynamic range for each standard curve (Table S5). We found a wide dynamic range (Table S5, dynamic ranges from 4.00 to 4.70) and a strong linear relationship between the $C_{RNA,\ ACL}$ and $C_{RNA,\ expected}$ (Fig. S9, $R^2$ ranging from 0.800 to 0.991) for standard curves from all tested cycle numbers.

We validate that the performance of ACL standard curves at different cycle numbers is comparable to that of *Ct* standard curves used in typical bulk RT-qPCR. With a *Ct* standard curve, we convert $F_N$ to $C_{RNA,\ Ct}$ for the same dilution series of IAV M gene RNA (Table S2). This conversion resulted in a wide dynamic range of 5 orders of magnitude and a strong linear response between $C_{RNA,\ Ct}$ and $C_{RNA,\ expected}$ of $R^2$ = 0.991 (Fig. S9, under *Ct* plot; Table S5 under *Ct* method).

To further quantify the variability between standard curves from different ACL cycle numbers, we calculated the coefficient of variation (*CV*) of $C_{RNA,\ ACL}$ replicates (Fig. S10A) and percent difference (*%*) between $C_{RNA,\ ACL}$ and $C_{RNA,\ expected}$ (Fig. S10B). *CV* (%) is the ratio of the standard deviation to the mean $C_{RNA,\ ACL}$, and is used to describe variability *within* replicates of the same M gene RNA concentration. We found non-linear relationships (Fig. S10A, black dotted lines with $R^2$ values < 0.28) between *CV* (%) and M gene RNA concentration at all sampled cycle numbers. We also calculated the percent difference (*%*) to describe variability *between* $C_{RNA,\ ACL}$ and $C_{RNA,\ expected}$ using Eq. S14:

$$\%\,\boldsymbol{difference} = \frac{\boldsymbol{log}(\boldsymbol{C_{RNA,ACL}}) - \boldsymbol{log}(\boldsymbol{C_{RNA,expected}})}{\boldsymbol{log}(\boldsymbol{C_{RNA,expected}})} * \mathbf{100}$$

**(Eq. S14)**

Again, we found a non-linear relationship across RNA concentrations (Fig. S10B, black dotted lines with $R^2$ values < 0.36). We note that variability in conversion with ACL standard curves is comparable to that of the *Ct* method, using *CV* (*%*) (Fig. S10A, *Ct*, $R^2$ = 0.02) and percent difference (*%)* (Fig. S10B, *Ct*, $R^2$ = 0.25).

**(R5) Validating ACL Standard Curves for Multiple Template Sequences.** ACL standard curves were also validated for two additional template sequences, the human MYCN gene DNA and soybean Lectin endogene DNA. These were compared to the previously discussed M gene dataset (Fig. S11A). The M gene dataset consisted of 21 RNA concentrations (Table S2, $3.28 \times 10^3$ to $2.62 \times 10^9$ copies/μL) used in previous positive control and validation experiments. The human MYCN gene and soybean Lectin endogene DNA datasets were taken from previously published studies on the evaluation of qPCR curve analysis methods (14) and enhanced analysis of qPCR data using a variable efficiency model (7). The MYCN dataset consisted of four DNA concentrations (ranging from $1.88 \times 10^0$ to $1.88 \times 10^3$ copies/μL) quantified with a SYBR Green qPCR assay. The Lectin dataset consisted of five dilutions (ranging from $6.4 \times 10^0$ to $4.0 \times 10^3$

copies/μL), also quantified with a SYBR Green qPCR assay. For each of the three template sequences, we constructed an ACL library from reference curves of $10^7$ copies/μL for the M gene, 1875 copies/μL for the MYCN gene, and 800 copies/μL for the Lectin endogene. The standard curve used in the M gene dataset was constructed from ACL cycle 20. The standard curves used in MYCN and Lectin datasets were from ACL cycles 27 and 29, respectively.

ACL conversions of $F_N$ to template concentration ($C_{RNA/DNA,\ ACL}$) for the IAV M gene RNA (Fig. S11A, green dots), human MYCN DNA (Fig. S11A, blue squares), and soybean Lectin endogene DNA (Fig. S11A, red circles) datasets were plotted as a linear regression against the known template concentrations of each dilution ($C_{RNA/DNA,\ expected}$). For each dilution, there were 3, 94, or 18 technical replicates for the M gene, MYCN, or soybean datasets, respectively. There was a clear linear relationship between $C_{RNA/DNA,\ ACL}$ and $C_{RNA/DNA,\ expected}$ for all datasets (Fig. S11A, $R^2$ > 0.9640). Only 2 of the 30 total $C_{RNA/DNA,\ ACL}$ values (from all datasets) fell outside a 2-fold change (Fig. S11A, black dotted lines) from $C_{RNA/DNA,\ expected}$. Of the 28 $C_{RNA/DNA,\ ACL}$ values (93% of the data) that fell within a 2-fold change, there was a maximum fold change of 1.7.

To further demonstrate the improved fit of reference amplification curves with the SCF-E model (Eq. S5) as opposed to the SCF model (8,13), the M gene dataset was also converted using a reference curve fit with the SCF model (Fig. S11A, gray dots). There was a weaker linear relationship between $C_{RNA,\ ACL}$ and $C_{RNA,\ expected}$ ($R^2$ = 0.8487), with 8 of the 21 $C_{RNA/DNA,\ ACL}$ values falling outside of the 2-fold from $C_{RNA,\ expected}$.

Furthermore, we compared ACL standard curves to $Ct$ standard curves for all three template sequences. Conversion with $Ct$ standard curves showed a similar linear relationship between $C_{RNA/DNA,\ ACL}$ and $C_{RNA/DNA,\ expected}$ for all datasets (Fig. S11B, $R^2$ > 0.9912). This time, all measured $C_{RNA/DNA,\ ACL}$ values fell within a 2-fold change of the expected mean with a maximum fold change of 1.9.

**(R6) Validating dqPCR for Multiple PCR Cycle Numbers.** To examine the contribution of error within individual PCR cycle numbers to overall error in the dqPCR distributions (Fig. 3B), we performed an Analysis of Variance (ANOVA) test (Fig. S15) on random samples of 100 drops from each M gene distribution in Fig. 3B. For the 17.1 M gene cpd group (sampled at $N$ = 22 to 25), measured cpd differed by cycle number (p-value <0.05) and accounted for 8% of the total error in pooled measurements. For the 171 M gene cpd group (sampled at $N$ = 19 to 22), measured cpd differed by cycle number (p-value <0.05) and accounted for 59% of the total error in pooled measurements. For the 1710 M gene cpd group (sampled at $N$ = 6 to 22), M gene measured cpd differed by cycle number (p-value <0.05) and accounted for 7% of the total error in pooled measurements. The higher error in the 171 cpd was an outlier, perhaps due to experimental variability from cycle 19, as error was reduced to 18% when that cycle was excluded; nevertheless, the other concentrations showed relatively lower error. To address this concern, we further examined the reported variability between cycle numbers using another metric, the linear mixed effects model (LME, Fig. S16). The percent difference (%) (Eq. S14) between the measured and expected M gene cpd was calculated at each sampled cycle number. The LME used cycle number as the fixed effect variable and expected M gene cpd as the random effect variable. Both variables were log-transformed during the model fit, as replication cycles correspond to RNA copy numbers on a log scale. The fixed effects coefficients described a negative relationship between the slope and percent difference (%) from the expected M gene cpd of -0.85% per cycle number (p-value <0.05, SE = 0.01). While the p-value indicates significance, the magnitude of the relationship between cycle and converted cpd is small (~1%), reinforcing the validity of pooling data converted at different cycle numbers together. Furthermore, when we pool cycle numbers in our control experiment (Fig. 3B), distributions were centered around the expected mean (Table S6) and had a linear relationship between measured and expected cpd, that also fell within a 2-fold change from the expected mean (Fig. S17).

**(R7) Extracting Individual Distributions from Mixed Droplet Detection Data.** To use dqPCR for measuring IAV burst size across a population of single-cell infections, which we expected to be heterogeneous, we validated that our method allows for isolation of individual distributions from a sample containing multiple M gene RNA concentrations. This was done by applying a Gaussian mixture model (GMM) (Table S7) to the distributions shown in Fig. 3E.

We began by fitting three log-normal distributions to the three individual concentrations of M gene cpd that make up the mixed drop distribution in Fig. 3E. We let $x_i$ be M gene cpd such that, at $x_1 = 1.71 \times 10^1$, $x_2 = 1.71 \times 10^2$, or $x_3 = 1.71 \times 10^3$cpd. The dqPCR model attributes measurement $y_i$ to each drop. Because of measurement noise, $x_i$ and $y_i$ are generally not equal. Mathematically, we represent the relationship between the measurement $y_i$ of a drop containing $x_i$ cpd, using Eq. S15:

$$\mathbf{log_{10}}(\boldsymbol{y_i}) = \mathbf{log_{10}}(\boldsymbol{x_i}) + \boldsymbol{E_i}$$

**(Eq. S15)**

Here, $E_i$ is the measurement noise. Inspired by the shape of the distribution in Fig. 3E, we assumed that $E_i$ follows a Gaussian distribution with mean $B_i$ (representing measurement bias) and variance $\sigma_i^2$ (representing the amount of measurement noise). Each Gaussian has a different weight corresponding to the probability that a random drop in the mixture has $x_i$ copies. Hence, to fit data in Fig. 3E, we needed to estimate these weights $w_i$ along with the $B_i$ and $\sigma_i^2$ values. The maximum likelihood estimates are represented in Table S7. In general, our model describes the measured distribution well, as seen from the fit in Fig. 3E.

Results from Table S7 suggest that the measurement bias $\widehat{B_i}$ decreases linearly with increases in the cpd on a log scale. We therefore used a linear function to describe the relationship in Eq. S16:

$$\boldsymbol{B(x)} \approx \boldsymbol{-0.06} \times \mathbf{log_{10}}(\boldsymbol{x}) + \boldsymbol{0.20}$$

**(Eq. S16)**

Here, the constants in Eq. S16 are estimated from the values reported in Table S7.

For the standard deviation of the measurement noise, $\widehat{\sigma_i}$, we notice that the change is non-monotonic with respect to increases in cpd. We therefore fitted a quadratic curve to describe their relationship in Eq. S17:

$$\boldsymbol{\sigma(x) = 0.63 - 0.47x + 0.11x^2}$$

**(Eq. S17)**

Here, the constants are also estimated from the values reported in Table S7.

**(R8) Filtering of Drops Containing Cell Lysate from Burst Size Distributions.** To ensure that burst size measurements are solely from extracellular viral particles, and do not include intracellular viral RNA from cell lysate, we developed a multiplexed dqPCR assay that simultaneously detects cellular β-actin mRNA and IAV M gene RNA. β-actin was chosen as a cellular indicator due to its abundance as a structural protein, its highly conserved sequence between cell types, and its linear relationship to cell concentration (Fig. S18). For each burst size replicate experiment (Table S8), two reference amplification curves were generated using $10^2$ cpd of M gene and β-actin template controls (Table S1) in 50 μm drops. As expected, drops containing high β-actin mRNA also exhibited high concentrations of M gene RNA (Fig. S19). These drops were presumed to contain non-packaged, intracellular M gene RNA. To remove these drops from the burst size data, we used one of two thresholds. The threshold was set at either the top 12.5% of drops containing β-actin, corresponding to the microfluidic chip split ratio, or above a β-actin concentration of $2 \times 10^3$ cpd, corresponding to the amount of β-actin released from a lysed cell (slope of Figure S18, ≈$2 \times 10^3$ β-actin copies per cell). The threshold which removed the most drops was chosen to ensure that drops containing lysed cells were excluded from the data.

**(R9) Oseltamivir (OST) treatment of IAV infected cells during drop infections.**

To demonstrate that burst size measurements include only RNA from extracellular viral particles, we measured IAV production from single cells treated with oseltamivir acid (OST), the active metabolite of the antiviral drug oseltamivir. OST functions as a neuraminidase inhibitor, preventing the cleavage of assembled IAV particles from the host cell membrane. MDCK cells infected with IAV H1N1 were encapsulated into 100 μm diameter microfluidic drops using methods for 'IAV Infection' and 'Droplet Encapsulation of Infected Cells' described in the main text. During encapsulation, infected cells were either suspended in standard droplet infection media (+H1N1 -OST) or in droplet infection media treated with 10 μM concentration of oseltamivir (+H1N1 +OST). Mock infected cells encapsulated in standard infection media (-H1N1 -OST) were used as a negative control. M gene abundance (copies/μL) was measured using a bulk RT-qPCR assay from the supernatant of broken drops sampled at 0 and 18 hpi. The 0 hpi bulk samples were frozen overnight at -80 °C while the 18 hpi bulk samples were incubated overnight at 37 °C. Preparation of drop infections for bulk RT-qPCR involved freezing a 400 μL sample of drops at -80 °C for 30 mins to break the emulsion. 300 μL of the broken emulsion was collected for further processing. Collected bulk and drop infections were clarified by centrifugation at 500 × *g* for 5 min, and the resulting supernatant was sampled for RT-qPCR targeting the IAV M gene. IAV production in drops had a mean of 1.930 x $10^3$ M gene copies/μl at 18 hpi, across three technical replicates (Figure S21). In the presence of oseltamivir**,** IAV production was detected in only one of three technical replicates, and measured 4.004 copies/μl. This was a significant decrease according to a two-sample Student's t-test ($p$ <<0.05). These results support that burst size measurements include only IAV particles released from the host cell membrane and are not contaminated by exosomal RNA or other forms of extracellular contamination, as oseltamivir prevents viral budding and release.

**(R10) Modeling IAV Burst Size Distributions from Droplet RNA Concentration.** The measured burst sizes distributions in Fig. 4B are compounded by two underlying distributions, the viral burst size distribution and the measurement noise. To infer the viral burst size distribution, we first estimated the measurement noise by fitting log-normal distributions to fluorescence data for a mixed sample of three known M gene RNA concentrations in drops (Fig. 3E, $1.71 \times 10^1$, $1.71 \times 10^2$, or $1.71 \times 10^3$ cpd). We found that the standard deviation of the distributions ranged from 0.15 - 0.30 on a $\log_{10}$ scale (Table S7) and used this to describe the measurement noise.

We then fit a model incorporating the measurement noise and three parametric distributions, Poisson, negative-binomial and log-normal (Fig. S22), against the burst size measurements in Fig. 4B. First, we simulated the viral burst size from the assumed distribution. For each simulated value of burst size $x$, we generated simulated measurement noise assuming a log-normal distribution with a mean defined by the bias function $B(x)$ and a standard deviation defined by $\sigma(x)$. Then, we computed a density function (kernel density with a Gaussian kernel) for the resulting distribution and used this function to compute the log-likelihood of observations. We estimated the parameters of each distribution and for each of the H3N2 or H1N1 populations by maximizing the likelihood of observations reported in Fig. 4B. To reduce the computational cost, we first obtained rough parameter estimates by simulating 10,000 values from each potential burst size distribution. The ensuing estimates were then refined by repeating the above procedure using 100,000 values while constraining parameter values to a domain closer to the rough estimates.

Comparing the goodness-of-fit of the three modeled distributions to the burst size data using Akaike Information Criterion (AIC) scores, we determined that the burst size measurements were best described by a negative-binomial distribution (Table S10 and Fig. 4D). The estimated mean burst sizes of these negative binomial distributions were 709 and 358 for H3N2 and H1N1, respectively. The shape parameter of each negative binomial distribution was estimated to be 0.48 and 0.49 for H3N2 and H1N1, respectively. Note that the negative-binomial distribution is considered highly dispersed when the shape parameter is less than 1. Therefore, there exists large heterogeneity in the burst size distribution for both H1N1 and H3N2.

#### Supplementary figures

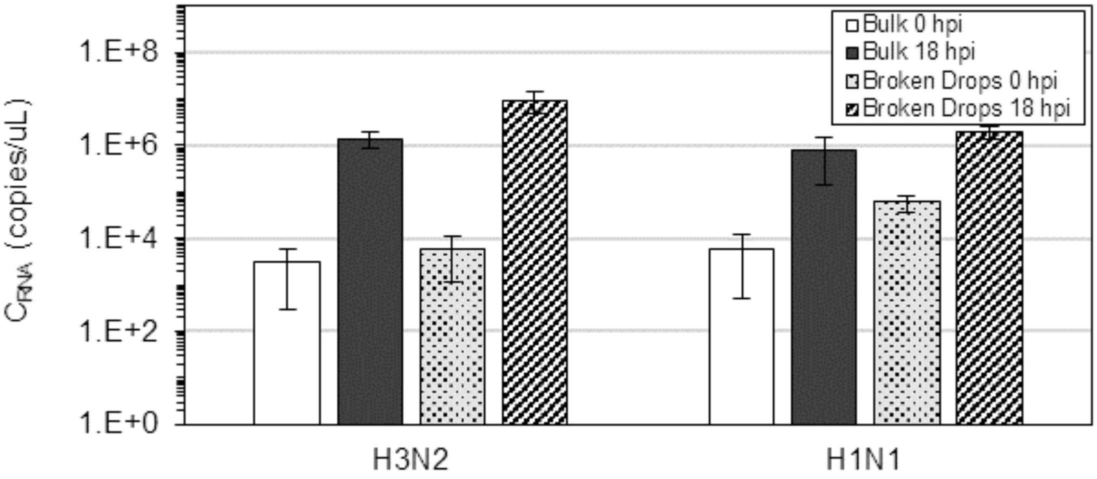


**Figure S1. Comparison of IAV M Gene Abundance During Bulk and Drop Infections.** IAV strains (H1N1 and H3N2) were separately used to infect A549 cells under bulk cell culture and single cell encapsulation in 100 µm diameter microfluidic drops. M gene abundance (copies/µL) was measured using a bulk RT-qPCR assay from the supernatant of both bulk and drop infections at 0 and 18 hpi. Experimental details can be found in **SI R1**. Each bar represents the pooled data from three replicate experiments. Error bars represent one standard deviation.

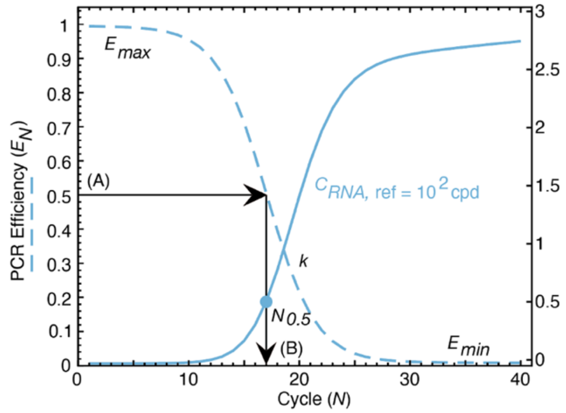


(Eq. S3) $$E_N = \frac{E_{max}}{1 + e^{\left(\frac{N - N_{0.5}}{k}\right)}} + E_{min}$$

(Eq. S4) $$F_{N+1} = F_N(1 + E_N)$$

(Eq. S5) $$F_{N+1} = F_N\left(1 + \frac{E_{max}}{1 + e^{\left(\frac{N - N_{0.5}}{k}\right)}} + E_{min}\right)$$

**Figure S2. Sigmoidal Curve Fitting using PCR Efficiency (SCF-E).** A four-parametric sigmoid function was used to model decreasing PCR efficiency ($E_N$) at each cycle number ($N$) (Eq. S3). This model was fitted to discontinuous fluorescence measurements ($F_N$) of a known RNA template concentration ($C_{RNA,\ ref}$) to generate a continuous reference amplification curve (Eq. S4 and S5). Reference curves used in burst size experiments were generated with $C_{RNA,\ ref} = 1.71 \times 10^2$ RNA (cpd). In the SCF-E model, $E_N$ is the PCR efficiency at cycle $N$; $E_{max}$ is the maximum PCR efficiency at cycle number 1 and ranges from of 0.9 to 1; $E_{min}$ is the minimum PCR efficiency at cycle number 40 and ranges from 0 to 0.1; $N_{0.5}$ is the cycle number at which $E_N$ equals 0.5; and $k$ is the shape parameter of the curve. As $k$ increases, the shape of the curve flattens, and as $k$ decreases, the shape of the curve becomes steeper. Thirty values of each of the four efficiency parameters ($E_{max}$, $E_{min}$, $N_{0.5}$, and $k$) were used to test $8.1\times10^5$ ($30^4$) curve fits to experimental fluorescence measurements. The set of efficiency parameters yielding the best fit were used to construct the SCF-E reference curve (solid blue curve).

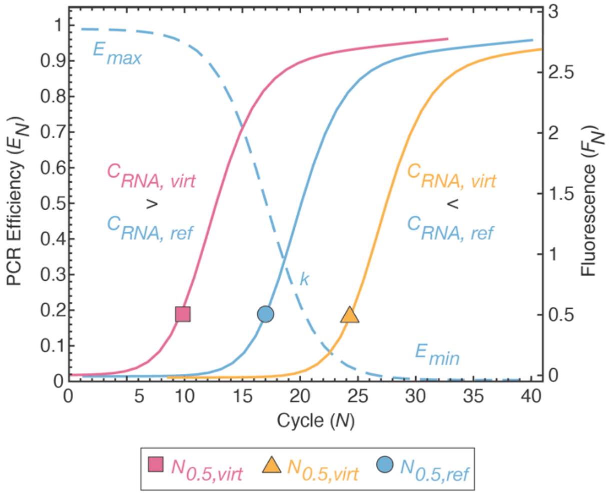


**(Eq. S7)**

$$F_{N+1} = F_N\left(1 + \frac{E_{max}}{1 + e^{\left(\frac{N - N_{0.5,virtual}}{k}\right)}} + E_{min}\right)$$

**Figure S3. Translating a Single SCF-E Reference Curve into 1000 Virtual Amplification Curves.** ACLs consist of virtual amplification curves that share the same $E_{max}$, $E_{min}$, and $k$ parameters as the reference curve but have unique $N_{0.5}$ values. This is done by substituting $N_{0.5, ref}$ from Eq. S5 with $N_{0.5, virt}$ in Eq. S7. We input 1000 evenly spaced $N_{0.5, virt}$ values between cycle numbers $N$ = 1 to 40, resulting in 1000 virtual amplification curves, each associated with a unique theoretical RNA concentration ($C_{RNA, virt}$). Virtual curves with $N_{0.5, virt} < N_{0.5, ref}$ (leftmost pink solid curve) correspond to higher RNA concentrations compared to the reference curve, whereas curves with $N_{0.5, virt} > N_{0.5, ref}$ (rightmost yellow solid curve) represent lower RNA concentrations than the reference (middle blue solid curve). The efficiency curve for reference and virtual curves is illustrated as a dashed blue line.

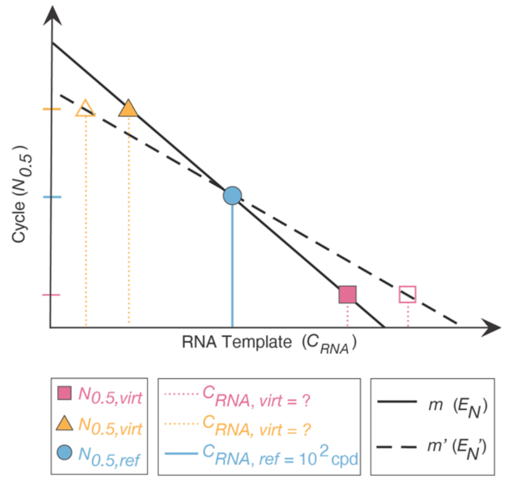


**(Eq. S8)**

$$N_{0.5} = m * log_{10}(C_{RNA}) + b$$

**(Eq. S10)**

$$log_{10}(C_{RNA,virt}) = log_{10}(C_{RNA,ref}) + \frac{N_{0.5,virt} - N_{0.5,ref}}{m}$$

**(Eq. S13)**

$$m = \frac{1}{log_{10}(E_N + 1)}$$

**Figure S4. Calculating RNA Template Concentration of the Virtual Amplification Curves.** We relate the known RNA concentration of the reference curve ($C_{RNA,\ ref}$) to the unknown RNA concentrations of the virtual curves ($C_{RNA,\ virt}$) based on their position on the x-axis (cycle number, *N*). In Eq. S10, we define a relationship between the known cycle numbers where PCR efficiency is equal to 0.5 for the reference ($N_{0.5,\ ref}$) and virtual ($N_{0.5,\ virt}$) curves and their respective RNA concentrations ($C_{RNA}$). By rearranging Eq. S10, we can calculate the unknown $C_{RNA,\ virt}$ using Eq. S12. To perform the calculation, the slope of the line (*m*) in Eq. S13 is determined using a PCR efficiency constant ($E_N$). As indicated by Eq. S13, an increase in $E_N$ leads to a decrease in *m* (dotted black line), expanding the range of $C_{RNA,\ virt}$ in the ACL (from solid triangle to open triangle, and from solid square to open square). This relationship is visually illustrated here with Eq. S8.

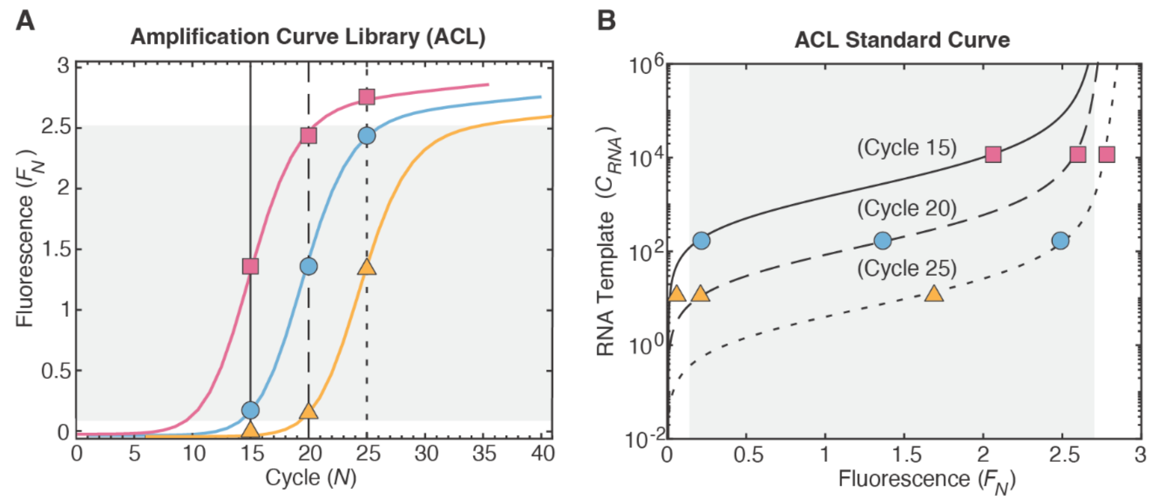


**Figure S5. Constructing a Standard Curve from the ACL. (A)** Construction of an amplification curve library (ACL) (Fig. S3). Three $C_{RNA,\ virt}$ within the ACL are displayed here (pink, blue, yellow), with the leftmost pink curve representing a higher $C_{RNA,\ virt}$ compared to the rightmost yellow curve. **(B)** ACL standard curves (solid line, dashed line, dotted line) are constructed from the $F_N$ of $C_{RNA,\ virt}$ (pink, blue, yellow) at a particular cycle number (15, 20, 25). The cycle number chosen for the ACL standard curve corresponds to the cycle number at which unknown RNA concentrations are sampled. In this method, only virtual curves in the exponential or linear phase at the selected cycle number can be included in the standard curve (grey shaded regions in **A** and **B**). This is because the early amplification and plateau regions of multiple virtual curves may overlap at a single cycle number. To ensure that only useable regions are selected for the standard curve, we establish threshold values based on the fluorescence of the reference curve. The upper threshold is set at 1 standard deviation below the $F_N$ at cycle 40, and the lower threshold is set at the 99$^{th}$ percentile of the $F_N$ at cycle 1. For example, as amplification proceeds for a particular $C_{RNA,\ virt}$ (pink curve) in **A**, the $F_N$ increases and intersects cycles 15, 20, and 25 at the three points (squares). However, only two of those points are within the valid range of the ACL standard curves in **B.**

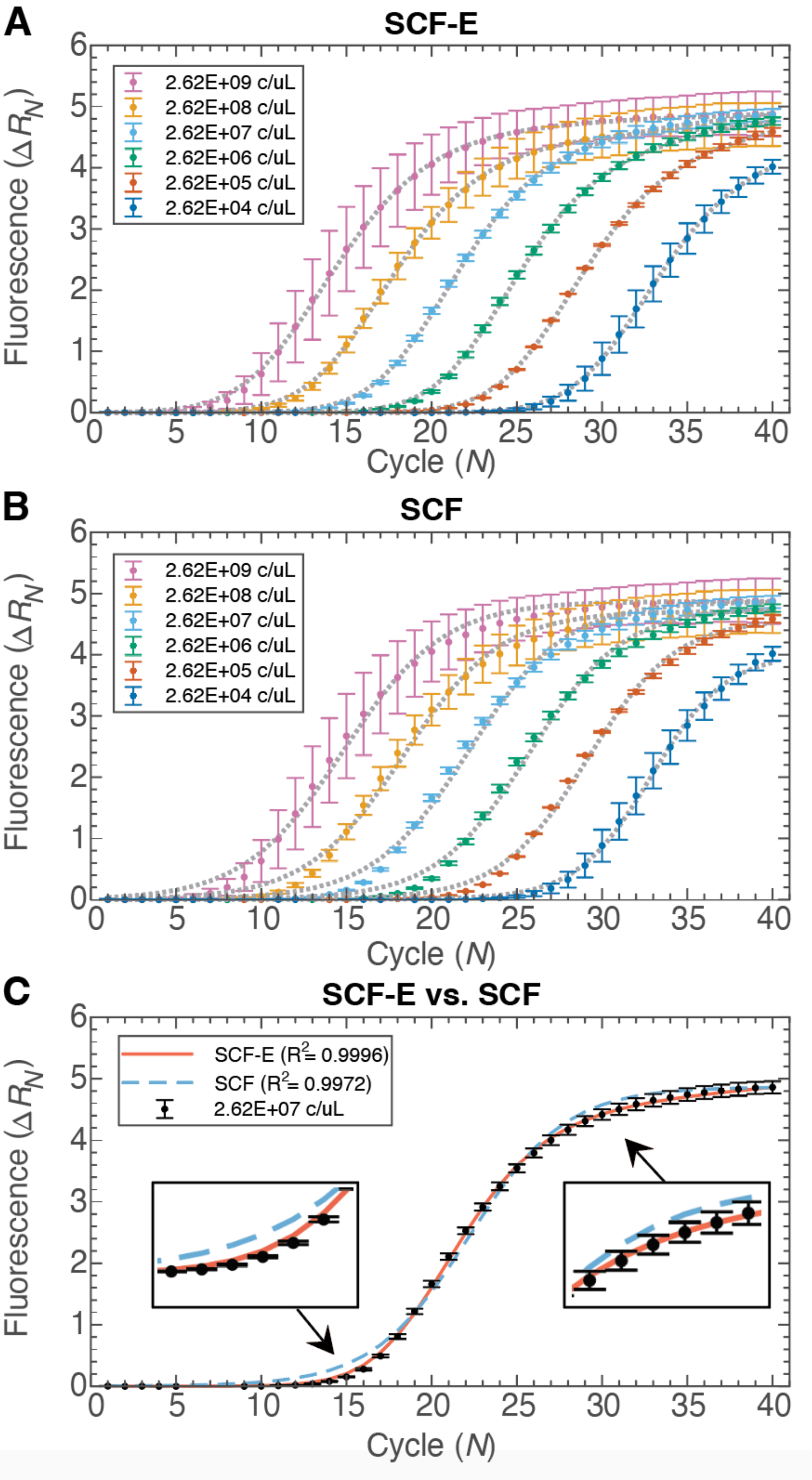


**(Continued on Next Page)**

**Figure S6. Comparison of SCF-E and SCF Methods of Fitting PCR Amplification Curves.** Fluorescence intensities ($\Delta R_N$) of bulk RT-qPCR amplification of six concentrations of IAV M gene RNA (Table S2, 10-fold dilutions from $2.62 \times 10^4$ to $2.62 \times 10^9$ copies/µL) were measured at $N = 1$ to 40. The concentration of each dilution decreases from left to right when viewing the amplification curves. **(A)** Amplification curves generated with the SCF-E model (dotted gray curves) had an average fit of $R^2 = 0.9996$ across all RNA concentrations (Table S3). **(B)** Amplification curves generated with the SCF model (dotted gray curves) had an average fit of $R^2 = 0.9975$ across all RNA concentrations (Table S3). **(C)** For a single M gene RNA concentration ($2.62 \times 10^7$ copies/µL), the SCF-E model provided a better fit ($R^2 = 0.9996$) than SCF ($R^2 = 0.9972$). The difference in fit is most noticeable at the lower ($\Delta R_N < 1$) and higher ($\Delta R_N > 4$) fluorescence values, representing the exponential and plateau phases of PCR amplification (magnified insets). All error bars represent one standard deviation from the average fluorescence measured across three technical replicates.

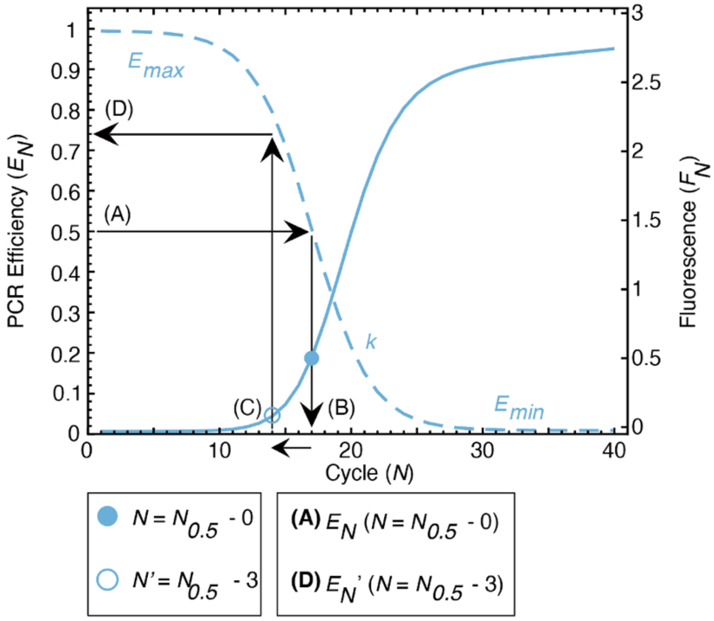


**(Eq. S3)**

$$E_N = \frac{E_{max}}{1 + e^{\left(\frac{N - N_{0.5}}{k}\right)}} + E_{min}$$

**(D)** $E_N'\,(N = N_{0.5}\ - 3)$

$$E_N{}' = \frac{E_{max}}{1 + e^{\left(\frac{-3}{k}\right)}} + E_{min}$$

**Figure S7. Setting a PCR Efficiency Constant (*$E_N$*) for the ACL.** To calculate $E_N$ at $N = N_{0.5} – 3$, we determined $N_{0.5}$ of the SCF-E reference curve, which corresponds to $N$ where $E_N = 0.5$ **(A to B)**. $N_{0.5} – 3$ represents three cycles prior to $N_{0.5}$ **(C)**. We recorded the $E_N$ at $N_{0.5} – 3$ **(D)**. When $N = N_{0.5} – 3$ is inserted into Eq. S3, the term $N_{0.5}$ cancels out, resulting in a constant value of -3 for ($N – N_{0.5}$). The $E_N$ calculated in this step is used in Eq. S13 to determine $m$.

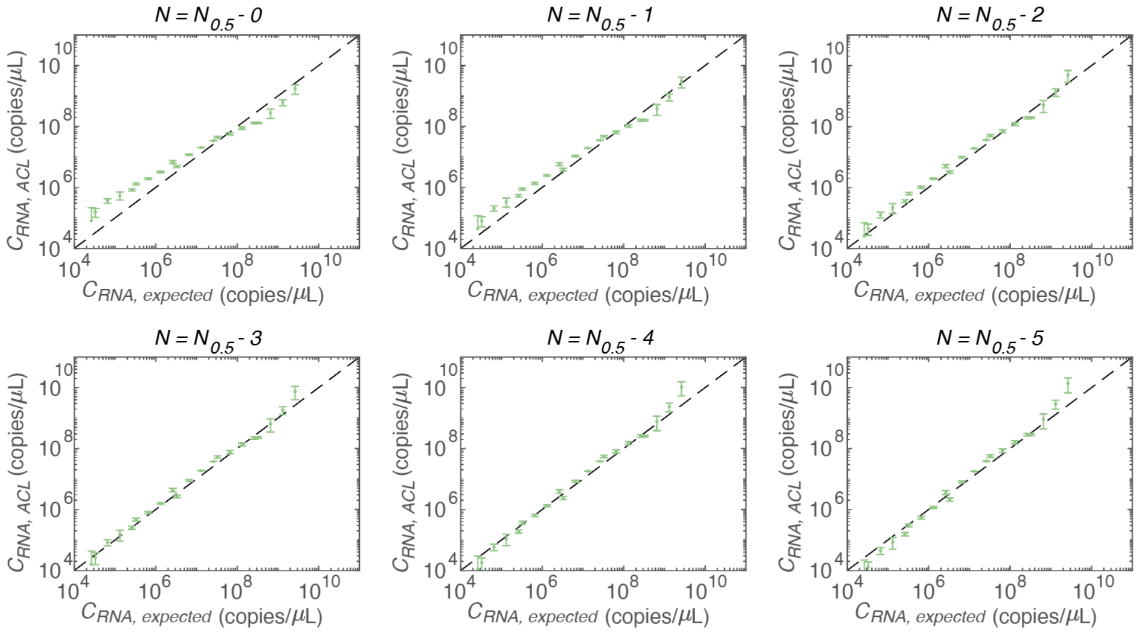


**Figure S8. Linear regression of measured to expected RNA concentrations from Table S4.** We compared cycle numbers (*N*) at which PCR efficiency ($E_N$, Eq. S3) of the reference curve is used to calculate the constant *m* (Eq. S13), for assigning RNA template concentrations to virtual curves in the ACL. We tested the cycle number at which PCR efficiency is equal to 0.5 ($N_{0.5}$) and the five cycles preceding it ($N = N_{0.5}$ *-1 to* $N = N_{0.5}$ *-5*), to construct six ACLs for comparison. From each ACL, a standard curve was constructed at PCR cycle number 20. We evaluated the dynamic range of each standard curve in Table S4. The ACL standard curve with the widest dynamic range and the highest degree of accuracy ($R^2$) between measured ($C_{RNA,\ ACL}$) and expected ($C_{RNA,\ expected}$) M gene RNA concentrations was built using $E_N$ at $N = N_{0.5} - 3$. For this ACL, the dynamic range of the standard curve was 4.70 $\log_{10}$ RNA copies/μL (Table S4, dynamic range of $2.62 \times 10^4$ to $1.31 \times 10^9$ copies/μL) with $R^2 = 0.990$ between $C_{RNA,\ ACL,}$ and $C_{RNA,\ expected}$. Therefore, the $E_N$ at $N = N_{0.5} - 3$ was used to construct all amplification curve libraries in this work.

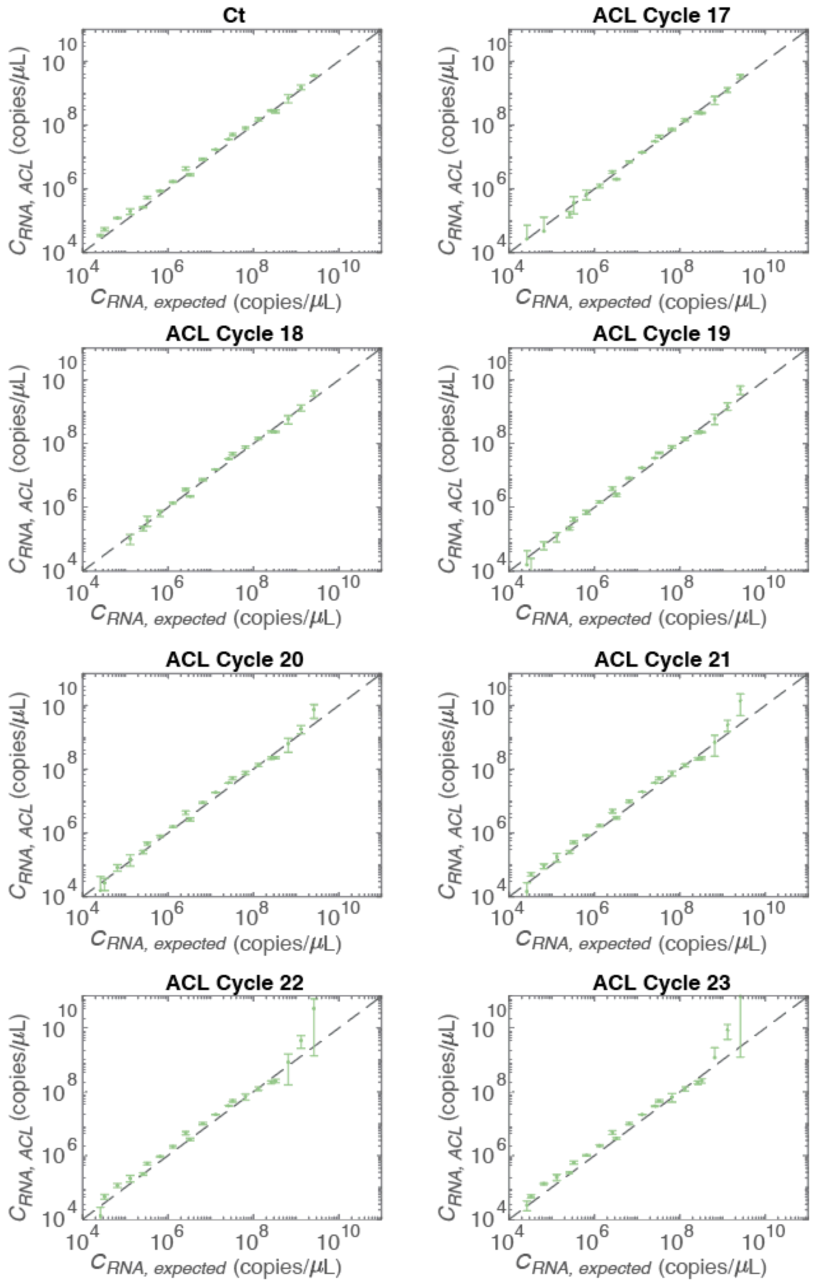


**Figure S9. Linear regression of measured to expected RNA concentrations from Table S5.** We compared the dynamic range of ACL standard curves from different cycle numbers ($N$ = 17 to 23) to the Ct standard curve. Standard curves were used to convert fluorescence to template concentration for a known dilution series of IAV M gene RNA (Table S2, three replicates of 21 concentrations between $2.62 \times 10^4$ to $2.62 \times 10^9$ copies/uL). Standard curve dynamic range and degree of accuracy ($R^2$) between measured ($C_{RNA,\ ACL}$) and expected ($C_{RNA,\ expected}$) M gene RNA concentrations were evaluated in Table S5.

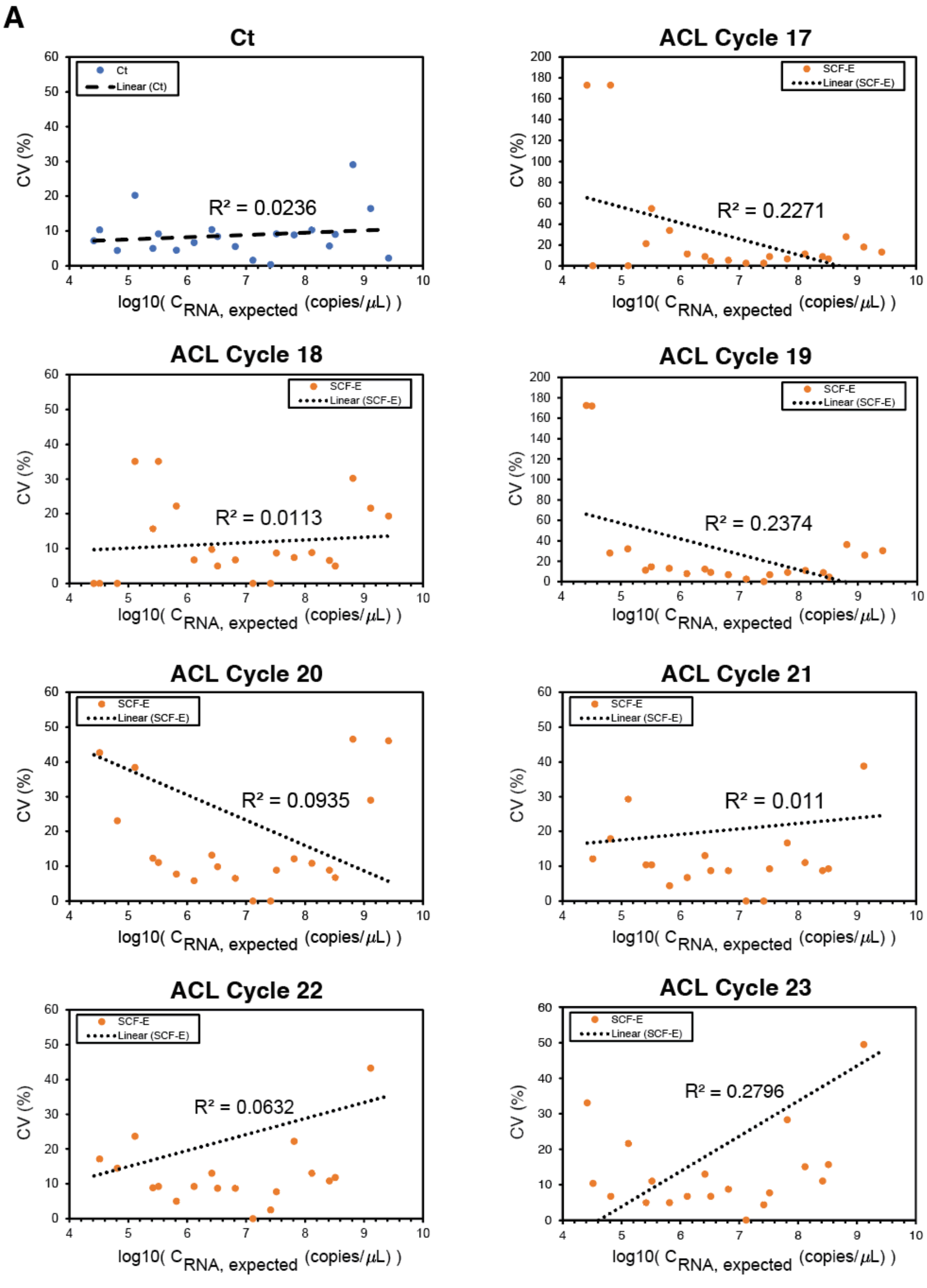


**(Continued on Next Page)**

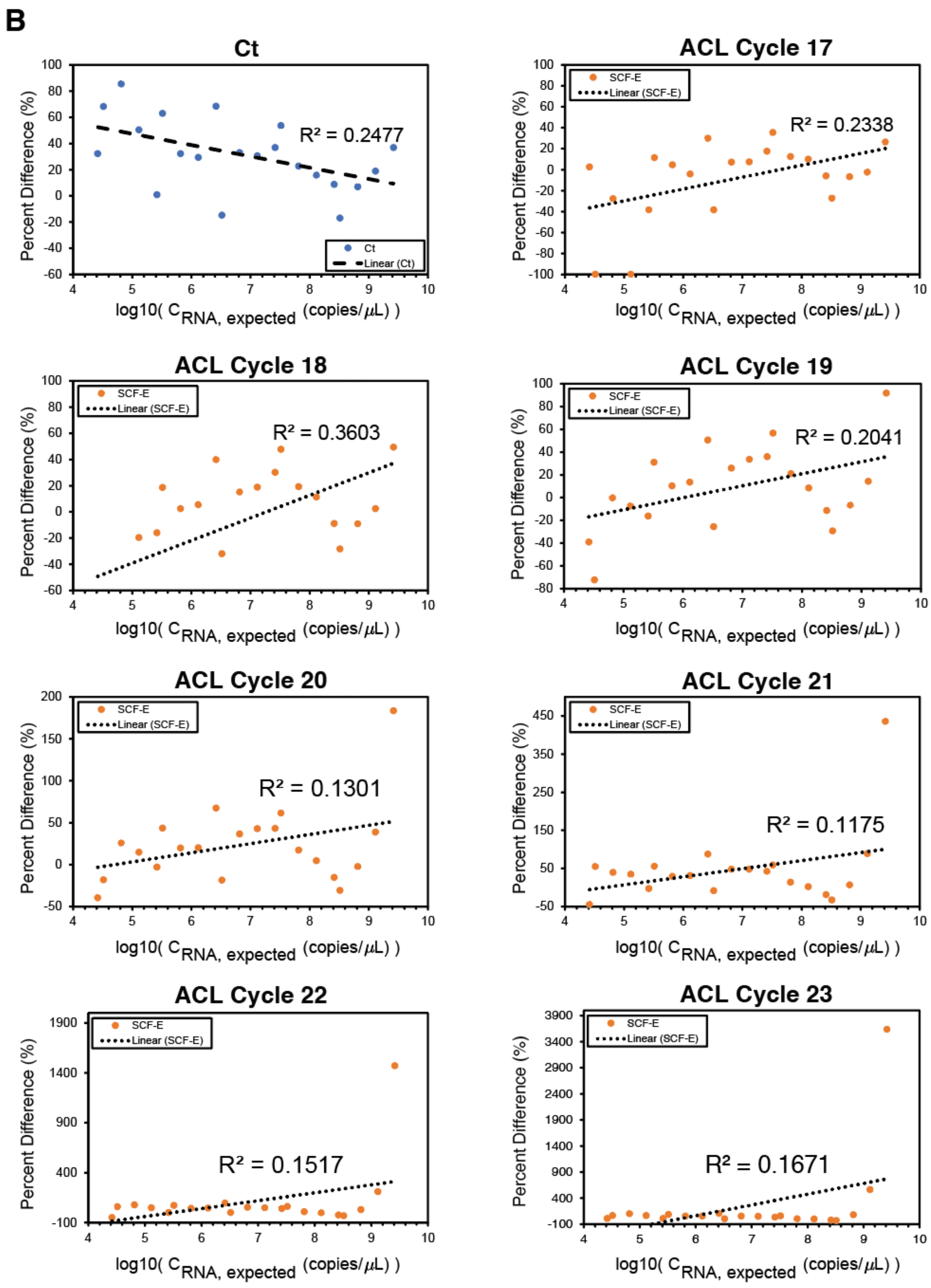


(Continued on Next Page)

**Figure S10. Variability in M gene RNA Concentrations Measured with ACL Standard Curves by PCR Cycle Number.** We examined variability between standard curves obtained from different ACL cycle numbers using two measures: **(A)** coefficient of variation (*CV* %) and **(B)** percent difference (%). Standard curves were constructed from ACL cycle numbers ($N$ = 17 to 23, individual plots) and used to convert fluorescence to RNA concentration ($C_{RNA,\ ACL}$) across 23 IAV M gene RNA concentrations (Table S2). **(A)** *CV* (%) is the ratio of the standard deviation to the mean $C_{RNA,\ ACL}$, and it quantifies the variability *within* replicates of the same M gene RNA concentration. A linear regression (dotted line) was conducted to compare *CV* (%) across M gene RNA concentrations. The results revealed a weak linear relationship ($R^2 < 0.28$) for all cycle numbers. **(B)** Percent difference (*%*) measures the variability *between* $C_{RNA,\ ACL}$ and the expected RNA concentration ($C_{RNA,\ expected}$). Similarly, a weak relationship ($R^2 < 0.36$) was observed between the starting RNA concentration and percent difference (%) for all cycle numbers. To assess the variability in ACL standard curves, a comparison was made with *Ct* standard curves (A and B, first upper left plot), for *CV* (%) and percent difference (%). The analysis indicated no linear response for *CV* (%) ($R^2 = 0.0236$) and a weak linear response for percent difference (%) ($R^2 = 0.2477$) across the M gene RNA concentrations.

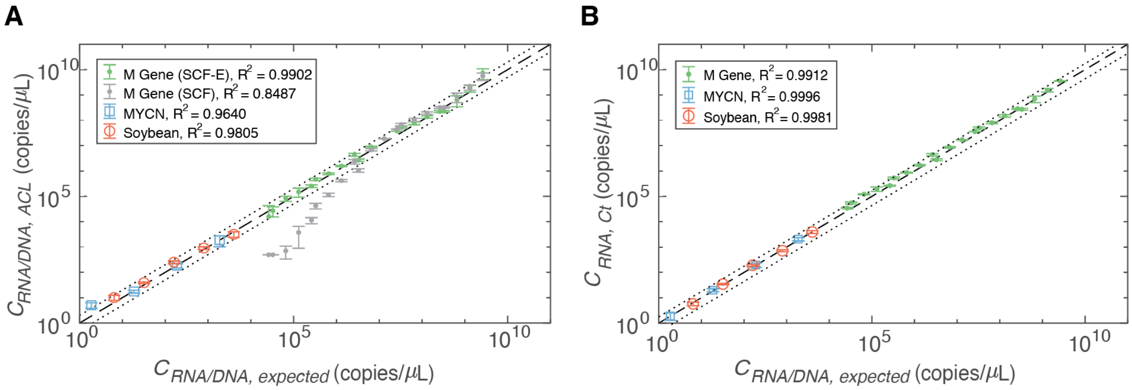


**Figure S11: Validating ACL Standard Curves for Multiple Template Sequences.** Standard curves constructed using the **(A)** ACL method or **(B)** Ct method were used to convert fluorescence to template concentration ($C_{RNA/DNA,\ ACL}$) for a known dilution series of three different template sequences: IAV M gene RNA (green/gray dots, ranging from $2.62 \times 10^4$ to $2.92 \times 10^9$ copies/μL), human MYCN gene DNA (blue squares, ranging from $1.88 \times 10^0$ to $1.88 \times 10^3$ copies/μL), and soybean Lectin endogene DNA (red circles, ranging from $6.4 \times 10^0$ to $4.0 \times 10^3$ copies/μL). Each dilution of the templates had varying numbers of technical replicates: 3 for the M gene, 94 for the MYCN, and 18 for the Lectin dataset. The M gene ACL was built with a $10^7$ copies/μL reference curve fitted with either the SCF-E (green dots) or SCF (grey dots) model. The standard curve from the M gene ACL was constructed at a single cycle number ($N$ = 20). The MYCN libraries were built with a 1,875 copies/μL reference curve, and the Soybean ACL was built with an 800 copies/μL reference curve, both fitted using the SCF-E model. The standard curves from these ACLs were constructed at $N$ = 27 (MYCN) and $N$ = 29 (Lectin). An $R^2$ value quantified the linear response between measured $C_{RNA/DNA,\ ACL}$ to expected concentrations ($C_{RNA/DNA,\ expected}$) across template concentrations. Dotted lines indicate a 2-fold change from $C_{RNA/DNA,\ expected}$, and error bars represent one standard deviation from the mean.

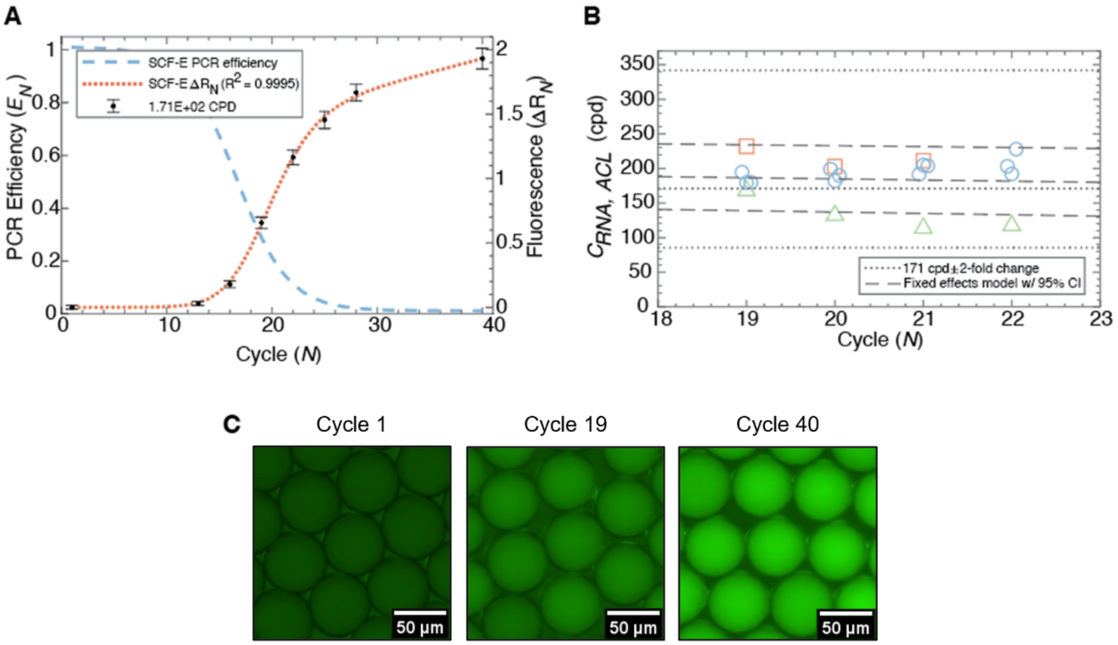


**Figure S12. Droplet Quantitative PCR (dqPCR) of the IAV M gene. (A)** SCF-E reference amplification curve generated for 1.71 x $10^2$ cpd of IAV M gene RNA amplified in 50 µm drops. Drop fluorescence intensity ($\Delta R_N$) was detected at PCR cycle numbers *N* = 1, 13, 16, 19, 22, 25, 28, 40 and displayed as a mean (black dots) and one standard deviation (error bars) (≈1000 or more drops per cycle number). The estimated PCR efficiency ($E_N$) curve (dashed blue line) used during SCF-E produced a well-fitting reference amplification curve (dotted red line, $R^2$ = 0.9995). **(B)** dqPCR was validated for a single concentration of M gene RNA (1.71x$10^2$ cpd) across three biological replicates. The samples were collected at various cycle numbers: rep 1 (red squares) at *N* = 19 to 21 and rep 2 (blue circles) and rep 3 (green triangles) at *N* = 19 to 22. Drop fluorescence was converted to M gene RNA concentration ($C_{RNA,\ ACL}$) within a 2-fold change of the expected mean (dotted lines) for all cycle numbers. Dashed lines indicate a fixed effects model with upper and lower bounds set as the 95% confidence intervals. **(C)** Representative epifluorescence images of 50 µm diameter drops at PCR cycle numbers 1, 19, and 40.

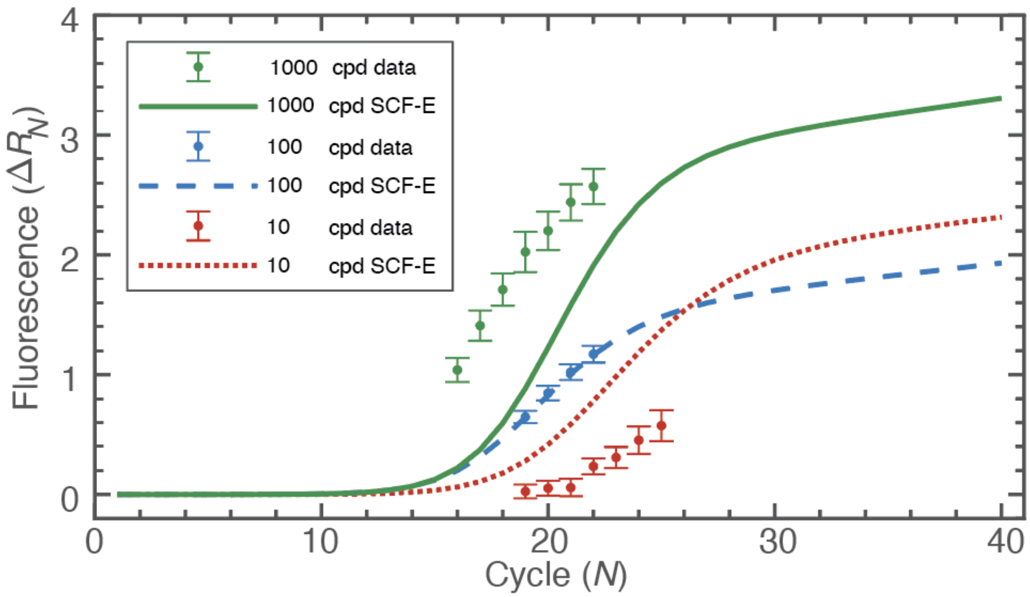


**Figure S13. SCF-E Reference Amplification Curves used in Figure 3B.** Three known concentrations of M gene RNA, 1.71 × $10^1$ cpd (red dots), 1.71 × $10^2$ cpd (blue dots), and 1.71 × $10^3$ cpd (green dots), were amplified in 50 μm drops. Drop fluorescence ($\Delta R_N$) was detected at multiple PCR cycle numbers (Table S6, note that cycles 19-21 in the 10 cpd group were below background and not included in the table) and converted to M gene cpd using dqPCR (Fig. 3B). For each experiment run on different days, a new SCF-E reference curve was generated from the amplification of 1.71 × $10^2$ cpd (red dotted line, blue dashed line, green solid line).

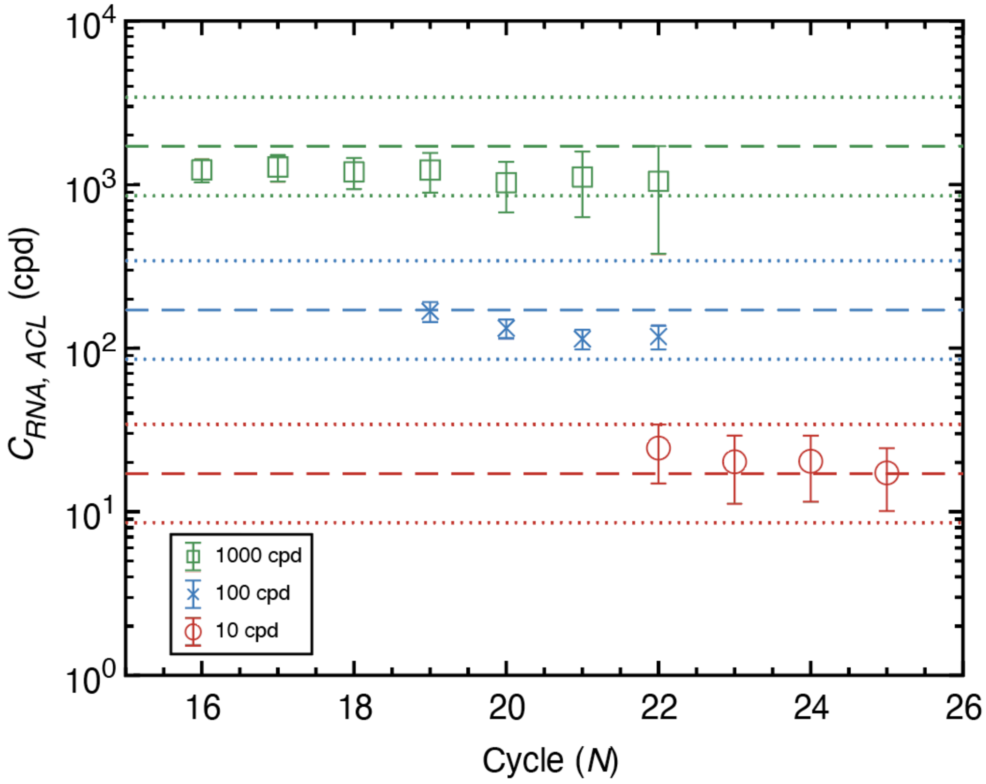


**Figure S14. Using dqPCR to Quantify Three IAV M gene RNA Concentrations at Multiple Cycle Numbers.** Three known IAV M gene RNA concentrations, $1.17 \times 10^1$ cpd (10 cpd, red circles), $1.17 \times 10^2$ cpd (100 cpd, blue crosses), and $1.17 \times 10^3$ cpd (1000 cpd, green squares) were amplified in 50 μm drops. Drop fluorescence ($\Delta R_N$) was detected at multiple PCR cycle numbers (Table S6) and converted to M gene cpd ($C_{RNA,\ ACL}$) using dqPCR. $C_{RNA,\ ACL}$ measurements were pooled together and displayed as a single distribution in Fig. 3B. Dashed lines indicate the expected M gene cpd, and dotted lines represent a 2-fold change from the expected mean.

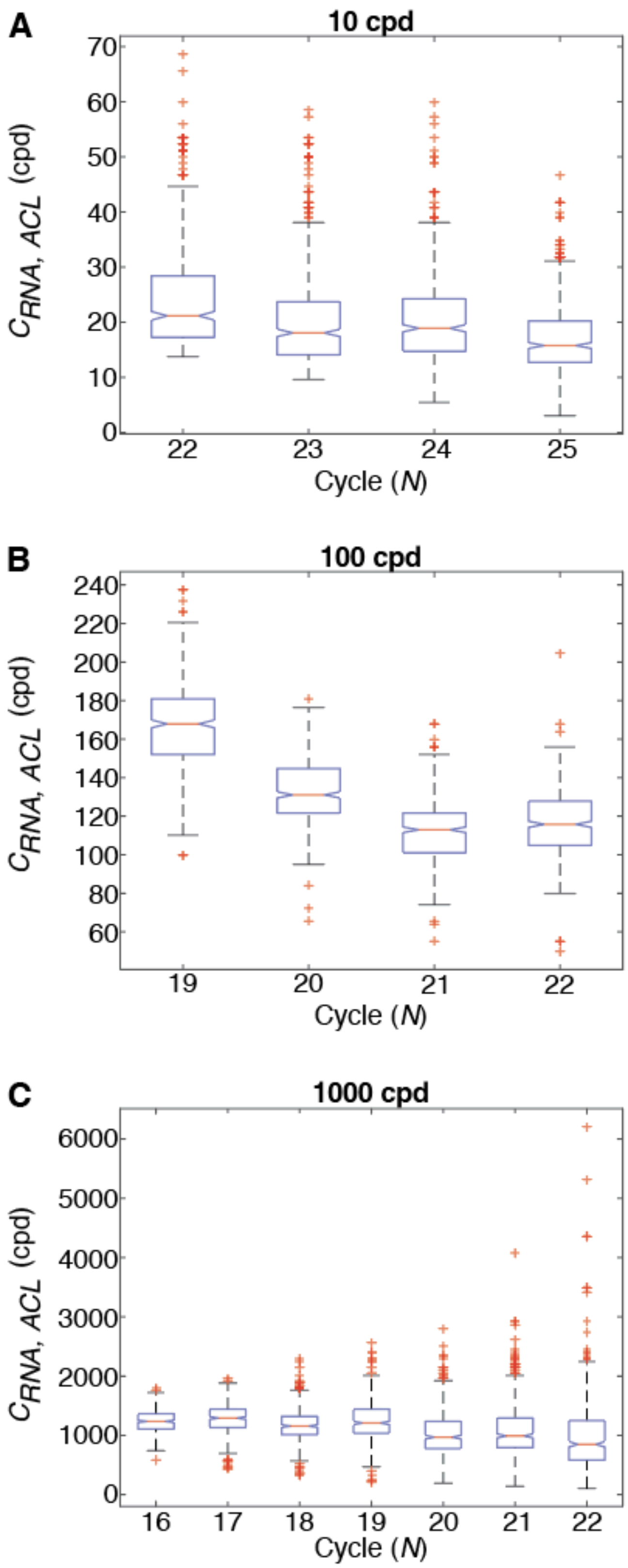


**(Continued on Next Page)**

**Figure S15. Analysis of Variance (ANOVA) for dqPCR of Three IAV M Gene RNA Concentrations.** Three known IAV M gene RNA concentrations, $1.17 \times 10^1$ cpd (10 cpd plot), $1.17 \times 10^2$ cpd (100 cpd plot), and $1.17 \times 10^3$ cpd (1000 cpd plot), were amplified in 50 μm drops. Drop fluorescence ($\Delta R_N$) was detected at multiple PCR cycle numbers (Table S6) and converted to M gene cpd ($C_{RNA,\ ACL}$) using the dqPCR model. $C_{RNA,\ ACL}$ measurements were pooled together and displayed as a single distribution in Fig. 3B. To examine the contribution of error within individual PCR cycle numbers to overall error in the pooled distributions, we performed an Analysis of Variance (ANOVA) test on a random sample of 100 drops from each group. **(A)** The 10 cpd group, quantified at PCR cycle numbers 22 – 25, had mean $C_{RNA,\ ACL}$ values that differed by cycle number (ANOVA p-value <0.05). This variability due to cycle number accounted for 8% of the total error in the pooled distribution. **(B)** The 100 cpd group, quantified at PCR cycle numbers 19-22, had mean $C_{RNA,\ ACL}$ values that differed by cycle number (ANOVA p-value <0.05). This variability due to cycle number accounted for 59% of the total error in the pooled distribution. **(C)** The 1000 cpd group, quantified at PCR cycle numbers 16-22, had mean $C_{RNA,\ ACL}$ values that differed by cycle number (ANOVA p-value <0.05). This variability due to cycle number accounted for 7% of the total error in the pooled distribution.

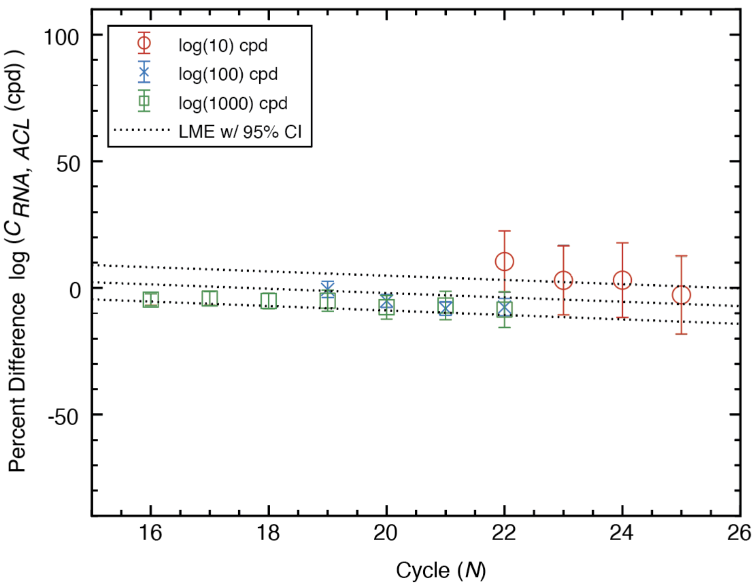


**Figure S16. Linear Mixed Effects Model (LME) for dqPCR of Three IAV M Gene RNA Concentrations.** Three known IAV M gene RNA concentrations, $1.17 \times 10^1$ cpd (10 cpd, red circle), $1.17 \times 10^2$ cpd (100 cpd, blue cross), and $1.17 \times 10^3$ cpd (1000 cpd, green triangle), were amplified in 50 µm drops. Drop fluorescence ($\Delta R_N$) was detected at multiple PCR cycle numbers (Table S6) and converted to M gene cpd ($C_{RNA,\ ACL}$). $C_{RNA,\ ACL}$ measurements were pooled together and displayed as a single distribution in Fig. 3B. To further examine the contribution of error within individual PCR cycle numbers to overall error in the pooled distributions, we calculated the percent difference (%) (Eq. S14) between the measured $C_{RNA,\ ACL}$ values and their expected M gene cpd ($C_{RNA,\ expected}$), across cycle numbers. This analysis informed the validity of pooling conversions measured from multiple PCR cycles in our burst size experiments. Percent differences were compared using a linear mixed effects model (LME) with a 95% confidence interval (CI) (dotted lines). In the LME, we set cycle as the fixed effect variable and $C_{RNA,\ expect}$ as the random effect variable. Both variables were log-transformed during the model fit, as replication cycles correspond to RNA copy numbers on a log scale. The fixed effects coefficients described a negative relationship between the slope and percent difference from the expected RNA concentration of -0.85% per cycle number (p-value <0.05, SE = 0.01). While the p-value indicates significance, the magnitude of the relationship is small (~1%/cycle), maintaining that it is valid to pool dqPCR data from different cycle numbers.

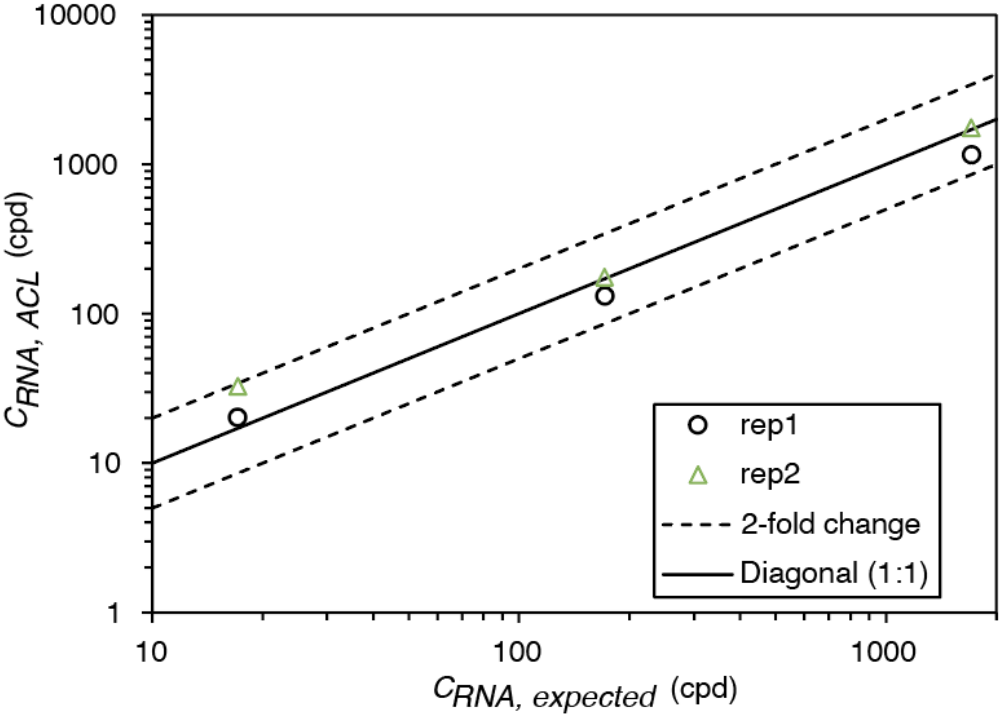


**Figure S17. Using dqPCR to Quantify IAV M gene RNA Concentration, Pooled Cycle Numbers.** Three known IAV M gene RNA concentrations, $1.17 \times 10^1$ cpd, $1.17 \times 10^2$ cpd, $1.17 \times 10^3$ cpd, were amplified in 50 µm drops. Drop fluorescence ($\Delta R_N$) was detected at multiple PCR cycle numbers (Table S6) and converted to M gene cpd ($C_{RNA,\ ACL}$) using dqPCR. $C_{RNA,\ ACL}$ measurements were pooled together and plotted against the expected M gene cpd ($C_{RNA,\ expected}$). Two biological replicates (black circle and green triangle) are shown. The solid line represents a 1:1 relationship between $C_{RNA,\ ACL}$ and $C_{RNA,\ expected}$, dashed lines indicate a 2-fold change from the expected mean.

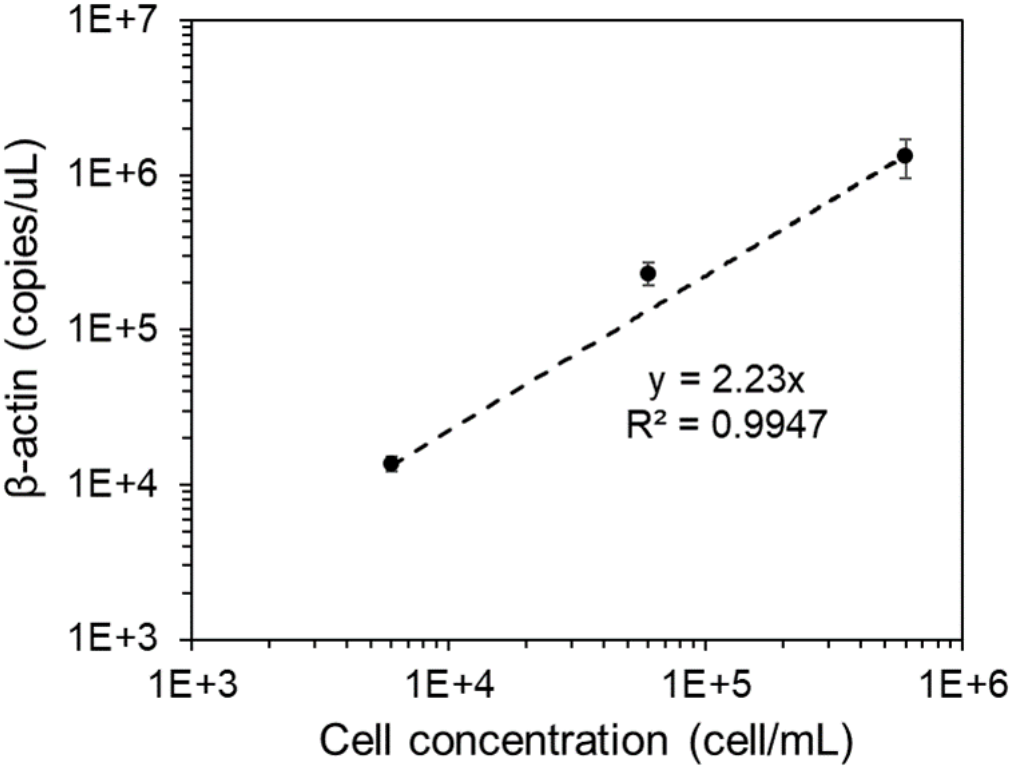


**Figure S18. β-actin mRNA abundance increases with cell concentration.** A bulk RT-qPCR assay was performed to amplify β-actin mRNA from lysed A549 cells after 24 hr of incubation. There was a strong linear relationship between β-actin and cell concentration (dashed black line, $R^2 = 0.995$), validating its use as a reliable marker for the detection of intracellular lysate in drops during our burst size experiments.

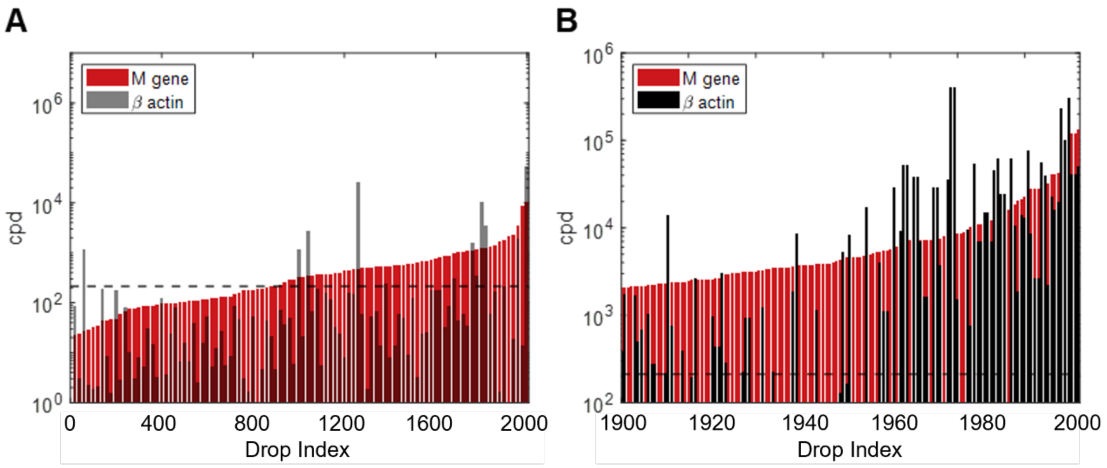


**Figure S19. Multiplexed dqPCR for Detection of IAV M gene and Cellular β-actin. (A)** Both M gene RNA (red) and β-actin mRNA (black) were detected by multiplexed dqPCR from H1N1 infection in drops (n=2000). Data is presented in ascending order of RNA concentration, prior to β-actin filtering (indicated by the dashed black line). **(B)** Showing a zoomed-in region of (A), with drop indexes 1900 to 2000.

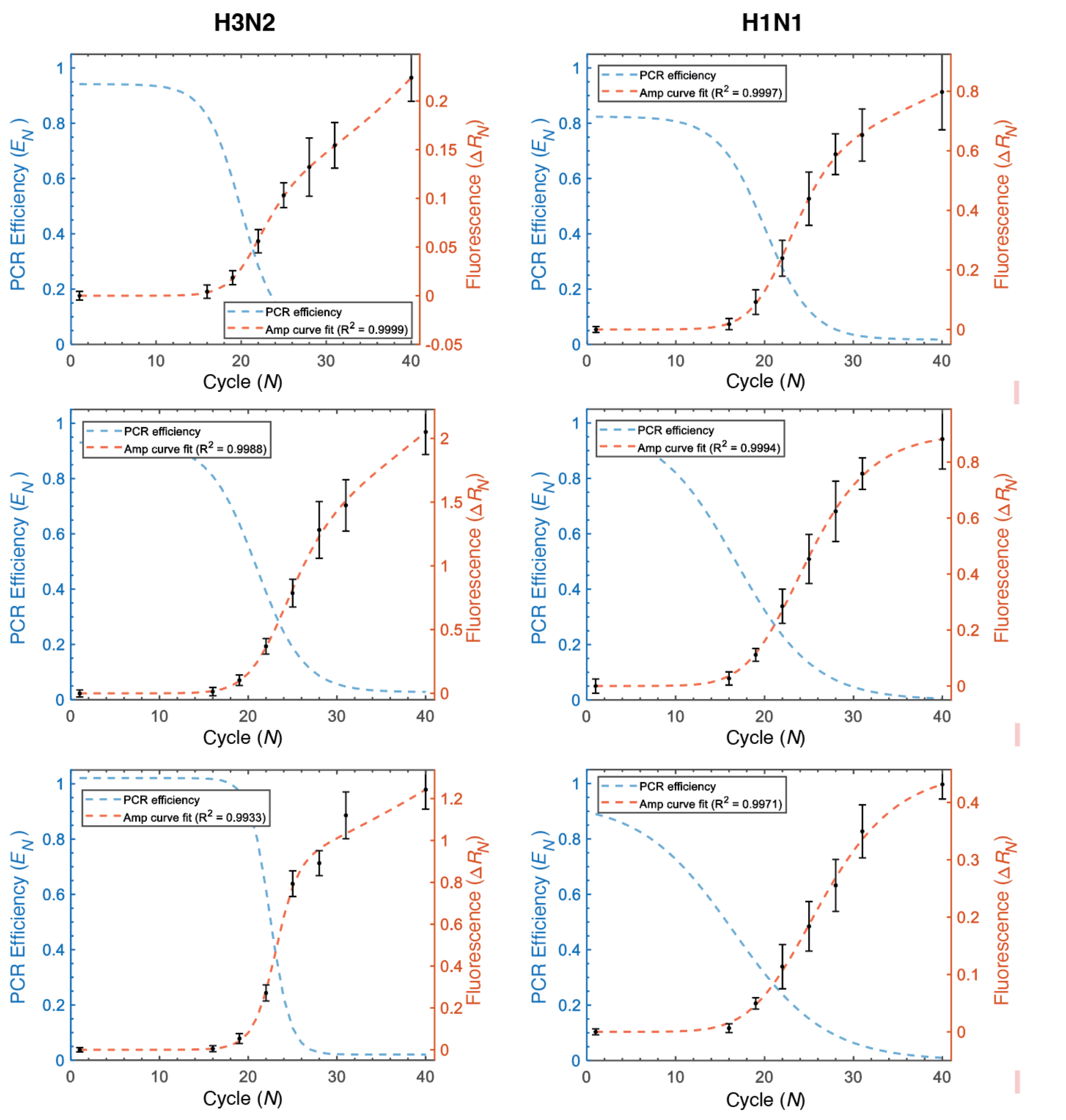


**Figure S20. SCF-E Reference Curves for Burst Size Replicate Experiments.** For each burst size replicate experiment (Table S8), a new reference amplification curve was constructed for dqPCR (Table S9). To generate these curves, a known concentration of M gene RNA (1.71 x $10^2$ cpd) was amplified in 100 μm drops. Drop fluorescence intensity ($\Delta R_N$) was measured at PCR cycle numbers $N$ = 1, 16, 19, 22. 25, 28, 31 and 40. The $\Delta R_N$ of drops from each cycle number are shown as the mean (black dots) with error bars representing one standard deviation. SCF-E was used to create a continuous reference amplification curve (orange dashed line) from discontinuous $\Delta R_N$ measurements, using an estimate for the PCR efficiency (blue dashed line).

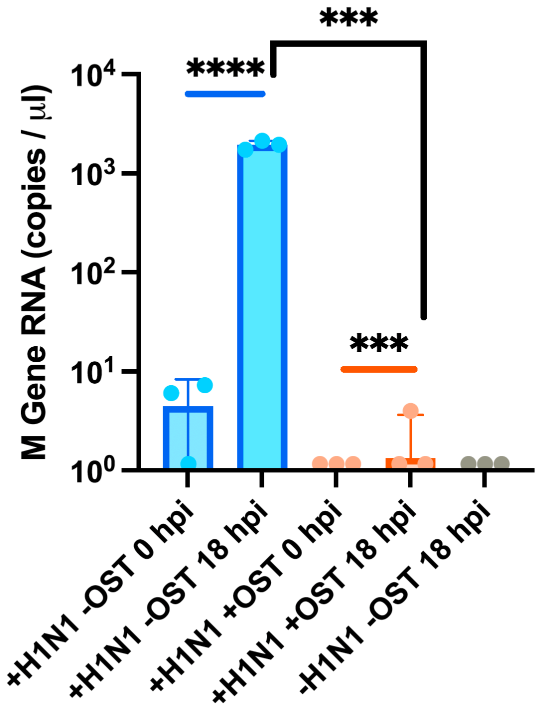


**Figure S21. Oseltamivir (OST) treatment of IAV infected cells during drop infections.** MDCK cells infected with IAV H1N1 were encapsulated into 100 µm diameter microfluidic drops. During encapsulation, infected cells were either suspended in standard droplet infection media (+H1N1 -OST) or in droplet infection media treated with 10 µM concentration of oseltamivir (+H1N1 +OST). Mock infected cells encapsulated in standard infection media (-H1N1 –OST) were used as a negative control. M gene abundance (copies / µL) was measured using a bulk RT-qPCR assay from the supernatant of broken drops sampled at 0 and 18 hpi. Each bar represents the pooled data from three technical replicates. Significance stars (*** $p < 0.001$ and **** $p < 0.0001$) obtained by a two-sample Student's t-test.

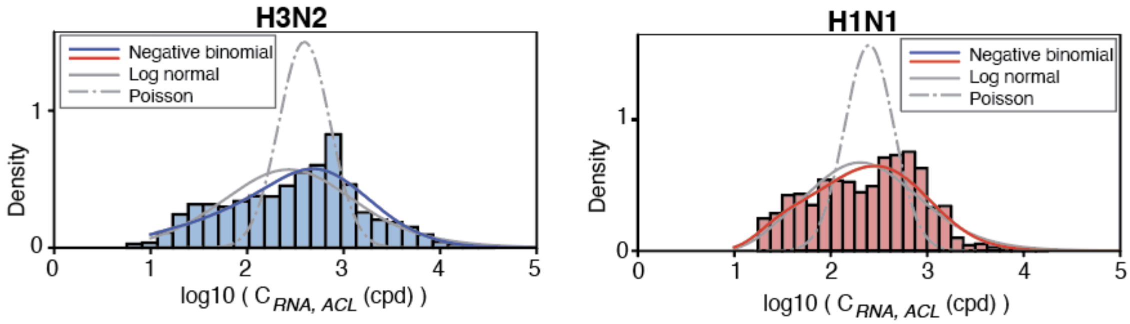


**Figure S22. Best Fit Model to Estimate IAV Burst Size Distributions from Table S10.**

We used a simulation-based approach to estimate the IAV burst distributions shown in Figure 4D. We considered three distribution models: lognormal (solid gray curve), Poisson (dashed gray curve), and negative-binomial distribution (solid blue or red curve, corresponding to H3N2 and H1N1 distributions, respectively). The parameters for each distribution are provided in Table S10. Based on the analysis, burst sizes were found to best fit a negative binomial distribution.

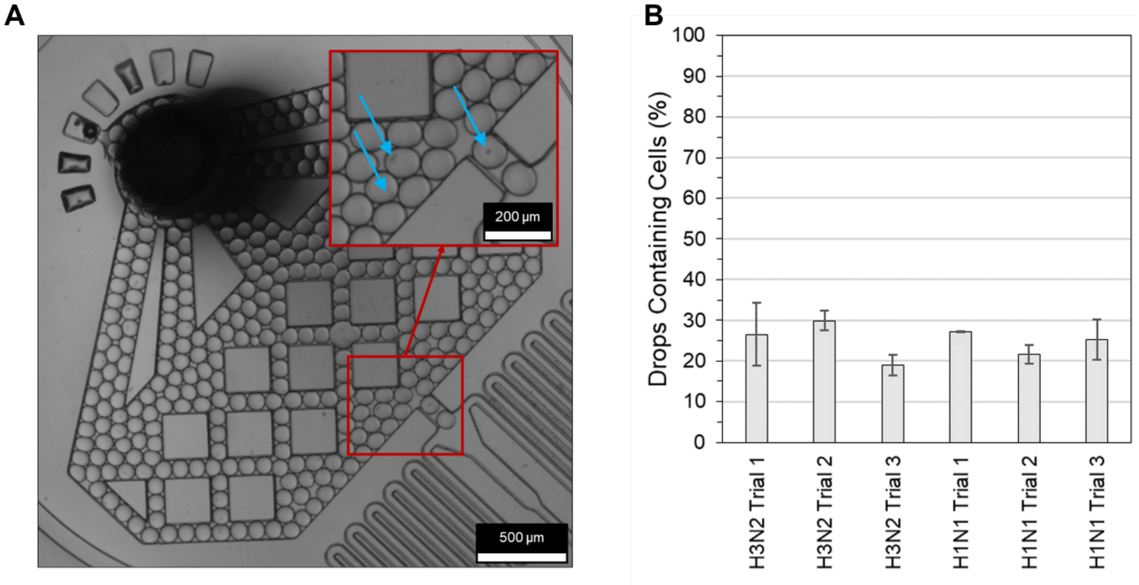


**Figure S23. Cell Loading During Burst Size Replicate Experiments**. **(A)** Images of 100 µm drops re-injected into a microfluidic device were captured at the beginning and end of each burst size replicate experiment to monitor cell loading. Drops containing a single cell are indicated by blue arrows. **(B)** Across all H3N2 and H1N1 replicate experiments, cell loading was determined by counting the percentage of drops containing cells (mean = 25% ± 4%). Cell loading in drops was assumed to follow a Poisson distribution. For a population of drops with 25% containing cells, the estimated Poisson mean ($\lambda$) is 0.29 cells/drop. In this case, ≈75% of drops are empty, ≈21.5% of drops contain one cell, ≈3% contain two cells, and ≈0.5% of drops contain three or more cells. The bar graph displays the drop counts for the replicates from left to right, which are 753, 679, 538, 719, 774, and 582.

878

### Supplementary tables

**Table S1. RT-qPCR Template and Primer Sequences**.

| | **Sequence (5' to 3')** |
|---|---|
| **M gene dsDNA template control** | GTCTAATACGACTCACTATAG*GACCAATCCTGTCACCTCTGAC***TGCAGTCCTCGCTCACTGGGCACG**TGCTTCATCGCGAACTGCTTCGCGGATGCCATCGTCATGGCCACGAGGATATGTAAGAGT*TAGACGATTTGTCCAGAATGCCCT* |
| **M gene forward primer** | GACCRATCCTGTCACCTCTGAC |
| **M gene reverse primer** | AGGGCATTCTGGACAAATCGTCTA |
| **M gene Taqman probe** | /FAM/TGCAGTCCTCGCTCACTGGGCACG/BHQ1/ |
| **β-actin dsDNA template control** | GTGGATGATGATATCGCCGCGCTCGTCGTCGACAACGGCTCCGGCATGTGCAAGGCCGGCTTCGCGGGCGACGATGCCCCCGGGCCGTCTTCCCCTCCATCGTGGGGCGCCCCAGGCACCAGGGCGTGATGGTGGGCATGGGTCAGAAGGATTCCTATGTGGGCGACGAGGCCCAGAGCAAGAGAGGCATCCTCACCCTGAAGTACCCCATCGAGCACGGCATCGTCACCAACTGGGACGACATGGAGAAAATCTGGCACCACACCTTCTACAATGAGCTGCGTGTGCCTCCCGAGGAGCACCCCGTGCTGCTGACCGAGGCCCCCCTGAACCCCAAGGCCAACCGCGAGAAGATGACCCAGATCATGTTTGAGACCTTCAACACCCCAGCCATGTACGTTGCTATCCAGGCTGTGCTATCCCTGTACGCCTCTGGCCGTACCACTGGCATCGTGATGGACTCCGGTGACGGGGTCACCCACACTGTGCCCATCTACGAGGGGTATGCCCTCCCCCATGCCATCCTGCGTCTGGACCTGGCTGGCCGGGACCTGACTGACTACCTCATGAAGATCCTCACCGAGCGCGGCTACAGCTTCACCACCACGGCCGAGCGGGAAATCGTGCGTGACATTAAGGAGAAGCTGTGCTACGTCGCCCTGGACTTCGAGCAAGAGATGGCCACGGCTGCTTCCAGCTCCTCCCTGGAGAAGAGCTACGAGCTGCCTGACGGCCAGGT |

| | |
|---|---|
| | CATCACCATTGGCAATGAGCGGTTCCGCTGCCCTGAGGCACTCTTCCAGCCTTCCTTCCTGGGCATGGAGTCCTGTGGCATCCACGAAACTACCTTCAACTCCATCATGAAGTGTGACGTGGACATCCGCAAAGACCTGTACGCCAACACAGTGCTGTCTGGCGGCACCACCATGTACCCTGGCATTGCCGACAGGATGCAGAAGGAGATCACTGCCCTGGCACCCAGCACAATGAAGATCAAGATCATTGCTCCTCCTGAGCGCAAGTACTCC*GTGTGGATCGGCGGCTCCATC*CTGGCCTCGCTGTCC**ACCTTCCAGCAGATGTGGATC**AG*CAAGCAGGAGTATGACGAGTC*CGGCCCCTCCATCGTCCACCGCAAATGCTTCTAG |
| **β-actin forward primer** | GTGTGGATCGGCGGCTCCATC |
| **β-actin reverse primer** | GACTCGTCATACTCCTGCTTG |
| **β-actin Taqman probe** | /Cy5/ACCTTCCAGCAGATGTGGATC/BHQ2/ |

All sequences are listed in the 5' to 3' direction. The M gene template control was constructed as linear dsDNA containing a T7 promoter (underlined), forward and reverse primer binding sites (*italicized*), a probe binding site (**bold**), and a partial M gene sequence. The M gene dsDNA was *in vitro* transcribed prior to use as a positive control. The M gene TaqMan probe carried a FAM fluorophore with a BHQ1 quencher. The β-actin template control was constructed as a dsDNA plasmid containing forward and reverse primer binding sites (*italicized*), and a probe binding site (**bold**), and a partial β-actin mRNA sequence. The β-actin TaqMan probe carried a Cy5 fluorophore with a BHQ2 quencher.

**Table S2. IAV M Gene RNA Concentrations Amplified by Bulk RT-qPCR and Used to Validate dqPCR.**

| M gene RNA concentration (copies/μL) of positive controls used in bulk RT-qPCR assays |
|---|
| **$2.62 \times 10^4$** |
| $3.28 \times 10^4$ |
| $6.55 \times 10^4$ |
| $1.31 \times 10^5$ |
| **$2.62 \times 10^5$** |
| $3.28 \times 10^5$ |
| $6.55 \times 10^5$ |
| $1.31 \times 10^6$ |
| **$2.62 \times 10^6$** |
| $3.28 \times 10^6$ |
| $6.55 \times 10^6$ |
| $1.31 \times 10^7$ |
| **$2.62 \times 10^7$** |
| $3.28 \times 10^7$ |
| $6.55 \times 10^7$ |
| $1.31 \times 10^8$ |
| **$2.62 \times 10^8$** |
| $3.28 \times 10^8$ |
| $6.55 \times 10^8$ |
| $1.31 \times 10^9$ |
| **$2.62 \times 10^9$** |

**Table S3. SCF-E and SCF Curve Fitting Parameters for Figure S6.**

| M gene RNA concentration (copies/µL) | $E_{min}$ (unitless) | $E_{max}$ (unitless) | $N_{0.5}$ (cycle) | $k$ (cycle) | $R^2$ (SCF-E) | $R^2$ (SCF) |
|---|---|---|---|---|---|---|
| $2.62 \times 10^9$ | 0.002 | 1.000 | 9.1 | 3.31 | 0.9994 | 0.9965 |
| $2.62 \times 10^8$ | 0.003 | 1.000 | 13.1 | 3.03 | 0.9995 | 0.9967 |
| $2.62 \times 10^7$ | 0.005 | 1.000 | 17.1 | 2.97 | 0.9996 | 0.9972 |
| $2.62 \times 10^6$ | 0.009 | 1.000 | 20.8 | 2.83 | 0.9997 | 0.9976 |
| $2.62 \times 10^5$ | 0.010 | 0.993 | 24.4 | 2.90 | 0.9998 | 0.9984 |
| $2.62 \times 10^4$ | 0.010 | 1.000 | 28.9 | 2.83 | 0.9997 | 0.9988 |
| **Average** | **0.007** | **0.999** | **18.9** | **2.98** | **0.9996** | **0.9975** |

**Table S4. Dynamic Range of ACL Standard Curves Built with Different $E_N$ Constants.**

| PCR Cycle ($N$) | $E_N$ | $m$ | $C_{RNA, min}$ (copies/µL) | $C_{RNA, max}$ (copies/µL) | Dynamic Range ($\log_{10}$(copies/µL)) | $R^2$ |
|---|---|---|---|---|---|---|
| $N_{0.5}$ - 0 | 0.514 | 5.55 | $3.28 \times 10^6$ | $1.31 \times 10^8$ | 1.60 | 0.886 |
| $N_{0.5}$ - 1 | 0.597 | 4.92 | $3.28 \times 10^6$ | $2.62 \times 10^8$ | 1.90 | 0.960 |
| $N_{0.5}$ - 2 | 0.675 | 4.46 | $2.62 \times 10^4$ | $2.62 \times 10^9$ | 5.00 | 0.986 |
| $N_{0.5}$ - 3 | 0.745 | 4.14 | $2.62 \times 10^4$ | $1.31 \times 10^9$ | 4.70 | 0.990 |
| $N_{0.5}$ - 4 | 0.805 | 3.90 | $3.28 \times 10^4$ | $1.31 \times 10^9$ | 4.60 | 0.986 |
| $N_{0.5}$ - 5 | 0.853 | 3.73 | $6.55 \times 10^4$ | $6.55 \times 10^8$ | 4.00 | 0.979 |

We compared cycle numbers ($N$) at which PCR efficiency ($E_N$) of the reference curve is used to calculate the constant $m$ (and $E_N$). This constant determines the RNA template concentration assigned to virtual curves in the ACL. $E_N$ is calculated using Eq. S3 and $m$ is calculated using Eq. S13. We used the cycle number at which PCR efficiency is equal to 0.5 ($N_{0.5}$) and the five cycles preceding it ($N = N_{0.5} -1$ to $N = N_{0.5} -5$), to construct six ACLs for comparison. From each ACL, a standard curve was generated at PCR cycle number 20. This standard curve was then used to convert fluorescence ($F_N$) to template concentration ($C_{RNA}$) for a known dilution series of M gene RNA (Table S2, three replicates of 21 concentrations between $2.62 \times 10^4$ to $2.62 \times 10^9$ copies/µL). The minimum ($C_{RNA, min}$) and maximum ($C_{RNA, max}$) concentrations in the dilution series, whose measured RNA concentration fell within a 2-fold change of the expected concentrations, marked the dynamic range of each standard curve. The $R^2$ value quantifies the linear regression (Fig. S8) between the measured ($C_{RNA, ACL}$) and expected ($C_{RNA, expected}$) RNA concentrations. The ACL built with $E_N$ at $N = N_{0.5} – 3$ produced a cycle 20 standard curve with the highest dynamic range and the greatest degree of accuracy ($R^2$). Therefore, an offset of -3 cycles was used to construct all ACLs in this work.

**Table S5. Dynamic Range of ACL Standard Curves from Different ACL Cycle Numbers.**

| Method | Standard Curve PCR Cycle (*N*) | $C_{RNA, min}$ (copies/μL) | $C_{RNA, max}$ (copies/μL) | Dynamic Range ($\log_{10}$(copies/μL)) | $R^2$ |
|---|---|---|---|---|---|
| *Ct* | NA | $2.62 \times 10^4$ | $2.62 \times 10^9$ | 5.00 | 0.991 |
| ACL | 17 | $2.62 \times 10^5$ | $2.62 \times 10^9$ | 4.00 | 0.812 |
| ACL | 18 | $1.32 \times 10^5$ | $2.62 \times 10^9$ | 4.30 | 0.800 |
| ACL | 19 | $6.55 \times 10^4$ | $2.62 \times 10^9$ | 4.60 | 0.989 |
| ACL | 20 | $2.62 \times 10^4$ | $1.31 \times 10^9$ | 4.70 | 0.990 |
| ACL | 21 | $2.62 \times 10^4$ | $1.31 \times 10^9$ | 4.70 | 0.981 |
| ACL | 22 | $3.28 \times 10^5$ | $6.55 \times 10^8$ | 4.30 | 0.962 |
| ACL | 23 | $6.55 \times 10^4$ | $6.55 \times 10^8$ | 4.00 | 0.938 |

We compared the dynamic range of standard curves created using the amplification curve library (ACL) method to that of the cycle threshold (*Ct*) method. Standard curves were used to convert fluorescence to template concentration for a known dilution series of IAV M gene RNA (Table S2, three replicates of 21 concentrations between $2.62 \times 10^4$ to $2.62 \times 10^9$ copies/μL). In the ACL method, multiple standard curves were constructed from individual cycle numbers ($N$ = 17 to 23). On the other hand, in the *Ct* method, a single standard curve is built from multiple *Ct* values, with one *Ct* value corresponding to each concentration of M gene RNA. The dynamic range of each standard curve was determined by identifying the minimum ($C_{\text{RNA, min}}$) and maximum ($C_{\text{RNA, max}}$) concentrations within the dilution series that yielded measured RNA concentration falling within a 2-fold change of the expected concentrations. The $R^2$ value quantifies the linear response (Fig. S9) between the measured ($C_{RNA, ACL}$) and expected ($C_{RNA, expected}$) RNA concentrations.

**Table S6. Using dqPCR to Quantify IAV M Gene RNA Concentrations at Multiple Cycle Numbers.**

| Group | M gene RNA concentration (cpd) | PCR Cycle (*N*) | Mean $C_{RNA, ACL}$ (cpd) | Standard Deviation (cpd) | *CV* (%) | Number of Drops (n) |
|---|---|---|---|---|---|---|
| $10^1$ | 17.1 | 22 | 24.49371 | 9.605813 | 39.21746 | 734 |
| | 17.1 | 23 | 20.18378 | 9.014579 | 44.66249 | 1472 |
| | 17.1 | 24 | 20.33817 | 8.819321 | 43.3634 | 1282 |
| | 17.1 | 25 | 17.26811 | 7.173752 | 41.54335 | 1025 |
| | **17.1** | **Pooled** | **20.2664** | **8.955352** | **44.18817** | **4513** |
| $10^2$ | 171 | 19 | 167.8969 | 23.34381 | 13.90366 | 3479 |
| | 171 | 20 | 132.3851 | 16.98913 | 12.83312 | 4268 |
| | 171 | 21 | 113.7438 | 15.53087 | 13.65426 | 5540 |
| | 171 | 22 | 117.5996 | 19.49319 | 16.57589 | 2612 |
| | **171** | **Pooled** | **131.2311** | **27.83475** | **21.21048** | **15899** |
| $10^3$ | 1710 | 16 | 1228.969 | 196.8349 | 16.01627 | 10107 |
| | 1710 | 17 | 1279.32 | 237.4656 | 18.56186 | 10443 |
| | 1710 | 18 | 1194.993 | 259.0783 | 21.68032 | 9186 |
| | 1710 | 19 | 1226.291 | 334.1291 | 27.24713 | 8743 |
| | 1710 | 20 | 1025.578 | 350.1932 | 34.14592 | 9569 |
| | 1710 | 21 | 1111.12 | 479.4927 | 43.15398 | 9335 |
| | 1710 | 22 | 1046.932 | 669.8305 | 63.98033 | 7878 |
| | **1710** | **Pooled** | **1163.231** | **390.8254** | **33.59827** | **65261** |

**(Continued on Next Page)**

Three known IAV M gene RNA concentrations, $1.71 \times 10^1$ cpd ($10^1$), $1.71 \times 10^2$ cpd ($10^2$), and $1.71 \times 10^3$ cpd ($10^3$), were amplified in 50 μm drops. Drop fluorescence ($\Delta R_N$) was detected at multiple PCR cycle numbers and converted to M gene cpd ($C_{RNA, ACL}$) using dqPCR. $C_{RNA, ACL}$ measurements from different cycle numbers were pooled together and presented as a single distribution in Figure 3B. To assess the variability between individual cycle numbers and the pooled distribution, we analyzed the standard deviation of the mean $C_{RNA, ACL}$ and the coefficient of variation (*CV %*) across the number of sampled drops (*n*) at each *N.*

**Table S7. Maximum Likelihood Estimates for a Gaussian Mixture Model (GMM) Used to Extract Individual Distributions from Mixed Droplet Detection Data.**

| $x_i$ (cpd) | Weights ($\widehat{w_i}$) | Bias ($\widehat{B_i}$) | Standard Deviation ($\widehat{\sigma_i}$) |
|---|---|---|---|
| $1.71 \times 10^1$ | 0.18 | 0.11 | 0.22 |
| $1.71 \times 10^2$ | 0.35 | 0.07 | 0.15 |
| $1.71 \times 10^3$ | 0.47 | -0.01 | 0.30 |

To use dqPCR for measuring IAV burst size in a heterogeneous population of single-cell infections, we validated the method's ability to isolate individual distributions from a sample containing multiple M gene RNA concentrations. We applied a Gaussian mixture model (GMM) on dqPCR data obtained from three known IAV M gene RNA concentrations: $1.17 \times 10^1$ cpd, $1.17 \times 10^2$ cpd, and $1.17 \times 10^3$ cpd, mixed together in a single sample (data shown in Figure 3E). Model assumptions are described in **SI R7**. In the model, $x_i$ represents the number of M gene RNA copies (cpd). Each Gaussian has a different weight ($w_i$) corresponding to the probability that a random drop in the mixture has $x_i$ copies, a predicted mean ($B_i$, representing measurement bias) and variance ($\sigma_i^2$, representing measurement noise).

**Table S8. Burst Size Replicate Experiments.**

| Strain | Replicate | Collected Drops (n) | Filtered Drops (n) | Mean Burst Size (viruses/cell) | Burst Size IQR (viruses/cell) | Gini coefficient (*G*) |
|---|---|---|---|---|---|---|
| H1N1 | 1 | 2141 | 1873 | $4.9 \times 10^2$ | $6.3 \times 10^1$ to $6.9 \times 10^2$ | 0.623 |
| H1N1 | 2 | 2840 | 1932 | $4.8 \times 10^2$ | $1.1 \times 10^2$ to $6.8 \times 10^2$ | 0.520 |
| H1N1 | 3 | 2036 | 1780 | $4.8 \times 10^2$ | $9.9 \times 10^1$ to $5.0 \times 10^2$ | 0.601 |
| **H1N1** | **Pooled** | **7017** | **5585** | $\mathbf{4.8 \times 10^2}$ | $\mathbf{8.8 \times 10^1}$ **to** $\mathbf{6.3 \times 10^2}$ | **0.586** |
| H3N2 | 1 | 4524 | 3955 | $1.6 \times 10^3$ | $1.1 \times 10^2$ to $1.6 \times 10^3$ | 0.724 |
| H3N2 | 2 | 4288 | 3751 | $6.3 \times 10^2$ | $1.1 \times 10^2$ to $8.1 \times 10^2$ | 0.581 |
| H3N2 | 3 | 2193 | 1914 | $3.8 \times 10^2$ | $5.3 \times 10^1$ to $5.8 \times 10^2$ | 0.518 |
| **H3N2** | **Pooled** | **11005** | **9620** | $\mathbf{1.0 \times 10^3}$ | $\mathbf{9.3 \times 10^1}$ **to** $\mathbf{8.9 \times 10^2}$ | **0.705** |

IAV burst sizes were measured from thousands of individual drops sampled at cycle numbers $N$ = 16, 19, 22, 25, and 28 in three biological replicates. The measurements were used to determine the number of IAV M gene RNA and cellular β-actin mRNA (cpd). Drops containing high β-actin cpd (Fig. S19) were assumed to contain non-packaged, intracellular, M gene RNA and were subsequently filtered from the final burst size distributions (Filtered Drops, *SI Appendix* R8). We present the average measured burst size for each replicate, along with the interquartile range (IQR) that represents the lower 25th and upper 75th percentile of the distributions. The heterogeneity of each distribution was examined using the Gini coefficient ($G$). If $G = 0$, the distribution is completely homogeneous, meaning that each cell produces the same number of viruses. Conversely, if $G = 1$, the distribution is completely heterogeneous, where one cell produces all the viruses. Thus, a higher $G$ corresponds to a more heterogeneous distribution.

**Table S9. dqPCR Parameters Used in Burst Size Replicate Experiments.**

| **Strain** | **Replicate** | $\boldsymbol{E_{min}}$ **(unitless)** | $\boldsymbol{E_{max}}$ **(unitless)** | $\boldsymbol{N_{0.5}}$ **(cycle)** | $\boldsymbol{k}$ **(cycle)** | $\boldsymbol{N_{0.5} - 3}$ | $\boldsymbol{E_N}$ | $\boldsymbol{m}$ |
|---|---|---|---|---|---|---|---|---|
| H1N1 | 1 | 0.0172 | 0.807 | 20.0 | 2.62 | 17.0 | 0.629 | 4.72 |
| H1N1 | 2 | 0 | 0.993 | 16.8 | 4.38 | 13.8 | 0.660 | 4.54 |
| H1N1 | 3 | 0 | 0.934 | 16.5 | 5.17 | 13.5 | 0.599 | 4.91 |
| H3N2 | 1 | 0.0414 | 0.907 | 20.0 | 2.03 | 17.0 | 0.779 | 4.00 |
| H3N2 | 2 | 0.0276 | 0.903 | 20.9 | 2.66 | 17.9 | 0.710 | 4.29 |
| H3N2 | 3 | 0.0207 | 1 | 22.4 | 1.14 | 19.4 | 0.954 | 3.44 |

The parameters used to create reference standard curves using SCF-E (Fig. S20, orange dotted curves) for each burst size experiment.

**Table S10. Best Fit Model to Estimate IAV Burst Size in Fig. 4D.**

| Strain | Burst size distribution | Parameter estimates | | Mean Burst Size (cpd) | AIC |
|---|---|---|---|---|---|
| H3N2 | Poisson | mean = 359.4 cpd | | 359 | 81541.5 |
| **H3N2** | **Negative Binomial** | **shape = 0.48** | **prob = 0.9993** | **709** | **19938.2** |
| H3N2 | Log-Normal | mean = 5.57 cpd | std= 1.70 | 1120 | 20302.4 |
| H1N1 | Poisson | mean = 222.4 cpd | | 222 | 25394.9 |
| **H1N1** | **Negative Binomial** | **shape = 0.49** | **prob = 0.9986** | **358** | **8689.0** |
| H1N1 | Log-Normal | mean = 5.24 cpd | std= 1.44 | 534 | 9044.2 |

We used a simulation-based approach to estimate the IAV burst distributions shown in Figure 4D. We considered three possible distributions: lognormal, Poisson, and negative-binomial, all with unknown parameters. First, we simulated the viral burst size based on the assumed distribution. For each simulated value of burst size $x$, we introduced measurement noise by assuming a log-normal distribution. The mean of the log-normal distribution was determined by the bias function $B(x)$, and the standard deviation was determined by $\sigma(x)$. Next, we computed a density function using a kernel density estimation with a Gaussian kernel to represent the resulting distribution. We then used this density function to calculate the log-likelihood of the observed data. We estimated the parameters for each distribution and for each dataset (H3N2 or H1N1) by maximizing the likelihood of observations reported in Fig. 4D.
